# Personalized CRISPR Knock-In Cytokine Gene Therapy to Remodel the Tumor Microenvironment and Enhance CAR T Cell Therapy in Solid Tumors

**DOI:** 10.1101/2025.04.25.650703

**Authors:** Michael Launspach, Julia Macos, Shoaib Afzal, Janik Hohmann, Stefanie Beez, Maximilian Pilgram, Emily Ohlendorf, Andrii Yevchuk, Casper F T van der Ven, Chahrazad Lachiheb, Lena Andersch, Marvin Jens, Marc Lukas Appis, Felix Zirngibl, Jonas Kath, Maria Stecklum, Kathleen Anders, Dimitrios Laurin Wagner, Ralf Kühn, Angelika Eggert, Annette Künkele

## Abstract

The immunosuppressive tumor microenvironment (TME) remains a central barrier to effective immunotherapy in solid tumors. To address this, we developed a novel gene therapeutic strategy that enables localized remodeling of the TME via tumor-intrinsic cytokine expression. Central to this approach is CancerPAM, a multi-omics bioinformatics pipeline that identifies and ranks patient-specific, tumor-exclusive CRISPR-Cas9 knock-in sites with high specificity and integration efficiency. Using neuroblastoma—a pediatric solid tumor with a suppressive TME—as a model, we applied CancerPAM to sequencing data from cell lines and patients to identify optimal integration sites for pro-inflammatory cytokines (CXCL10, CXCL11, IFNG). CRISPR-mediated CXCL10 knock-in into tumor cells significantly enhanced CAR T cell infiltration and antitumor efficacy both in vitro and in vivo. In vivo, CXCL10-expressing tumors showed significantly increased early CAR T cell infiltration and prolonged survival compared to controls. CancerPAM rankings correlated strongly with target-site specificity and knock-in efficiency, validating its predictive performance. Our findings establish CancerPAM as a powerful tool for safe and effective CRISPR-based interventions and provide a conceptual framework for integrating cytokine-driven TME remodeling with cellular immunotherapies. This personalized strategy holds promise for enhancing CAR T cells and other immunotherapies across immune-refractory solid tumors.

## Introduction

The emergence of CRISPR-Cas9 technology has transformed genome engineering by enabling precise gene editing. Guided by single-guide RNAs (sgRNAs), the Cas9 nuclease induces site-specific double-strand breaks, allowing targeted gene knockouts and insertions. This versatility has established CRISPR-Cas9 as an indispensable tool in both genetic research and therapy development^1,2^. Harnessing CRISPR for cancer therapies necessitates highly specific target selection to ensure both safety and efficacy. Automated bioinformatics pipelines integrating multi-omics (genomic, transcriptomic and epigenomic) data have been developed either to advance understanding of cancer biology^3,4^ or to facilitate optimal CRISPR gRNA design, to maximize therapeutic precision^5^. However, tools that effectively combine both, are still lacking.

Despite advances in precision oncology, solid tumors remain a major global health burden^6^. Directly targeting tumor cells through cytotoxic gene therapy is hindered by the necessity to deliver the therapy to every cancer cell to prevent relapse, a requirement current delivery vehicles fail to meet^7–9^. Immunotherapies have emerged as a promising alternative, leveraging the immune system’s ability to recognize and eliminate tumors. Oncolytic viruses, neoantigen-based vaccines and chimeric antigen receptor (CAR) T cell therapy have shown considerable potential^10–12^. CAR T cells, genetically modified to express synthetic receptors that combine tumor antigen recognition with intracellular signaling domains, directly recognize target antigens and mount an cytotoxic response^13^. However, the immunosuppressive tumor microenvironment (TME) in many solid tumors presents a significant hurdle to CAR T cell therapy. Tumors deploy various immune evasion mechanisms, including secretion of immunosuppressive cytokines and recruitment of regulatory T cells, which collectively hinder CAR T cell infiltration and cytotoxic activity^14,15^.

Neuroblastoma is a pediatric solid tumor of neuroectodermal origin having an immunosuppressive TME. Despite aggressive multimodal treatment regimens, high-risk neuroblastoma is associated with poor long-term survival^16,17^. CAR T cell therapies targeting antigens on neuroblastoma cells, such as L1CAM or GD2 have been explored^18,19^. While CAR T cell therapy against GD2 has shown some promise (NCT03373097), other clinical trials, including those targeting L1CAM (NCT02311621), have reported limited efficacy due to antigen escape and poor infiltration and persistence in the hostile TME^20–22^. *MYCN*-amplification, occurring in ∼25% of neuroblastomas, downregulates IFNG signaling and creates a T cell-excluding TME^23,24^.

Here, we propose a novel gene therapeutic approach that remodels the TME by inducing tumor-intrinsic expression of immunostimulatory cytokines such as CXCL10, CXCL11 and IFNG. CXCL10 and CXCL11, ligands of CXCR3 expressed on activated T and NK cells, have been shown to enhance immune cell infiltration and improve responses to immunotherapies in solid tumors^25–28^. IFNG, a potent immunostimulatory cytokine, augments anti-tumor immunity and induces CXCL10 and CXCL11 expression, further amplifying immune cell recruitment and activity^29,30^. To implement this concept with precision, we developed CancerPAM—an integrative multi-omics pipeline that identifies and ranks tumor-specific CRISPR knock-in sites based on specificity, efficiency, and safety criteria. Using neuroblastoma as a model, we demonstrate that CancerPAM enables efficient, site-specific integration of cytokine transgenes into tumor genomes to improve CAR T cell infiltration and efficacy **(Fig. 1).** Functional validation in vitro and in vivo confirms that tumor-intrinsic CXCL10 expression enhances CAR T cell infiltration and improves tumor control. These findings highlight tumor-intrinsic cytokine expression as a promising strategy to enhance solid tumor immunotherapy and establish CancerPAM as a powerful tool for CRISPR-based precision interventions.

**Fig. 1.**
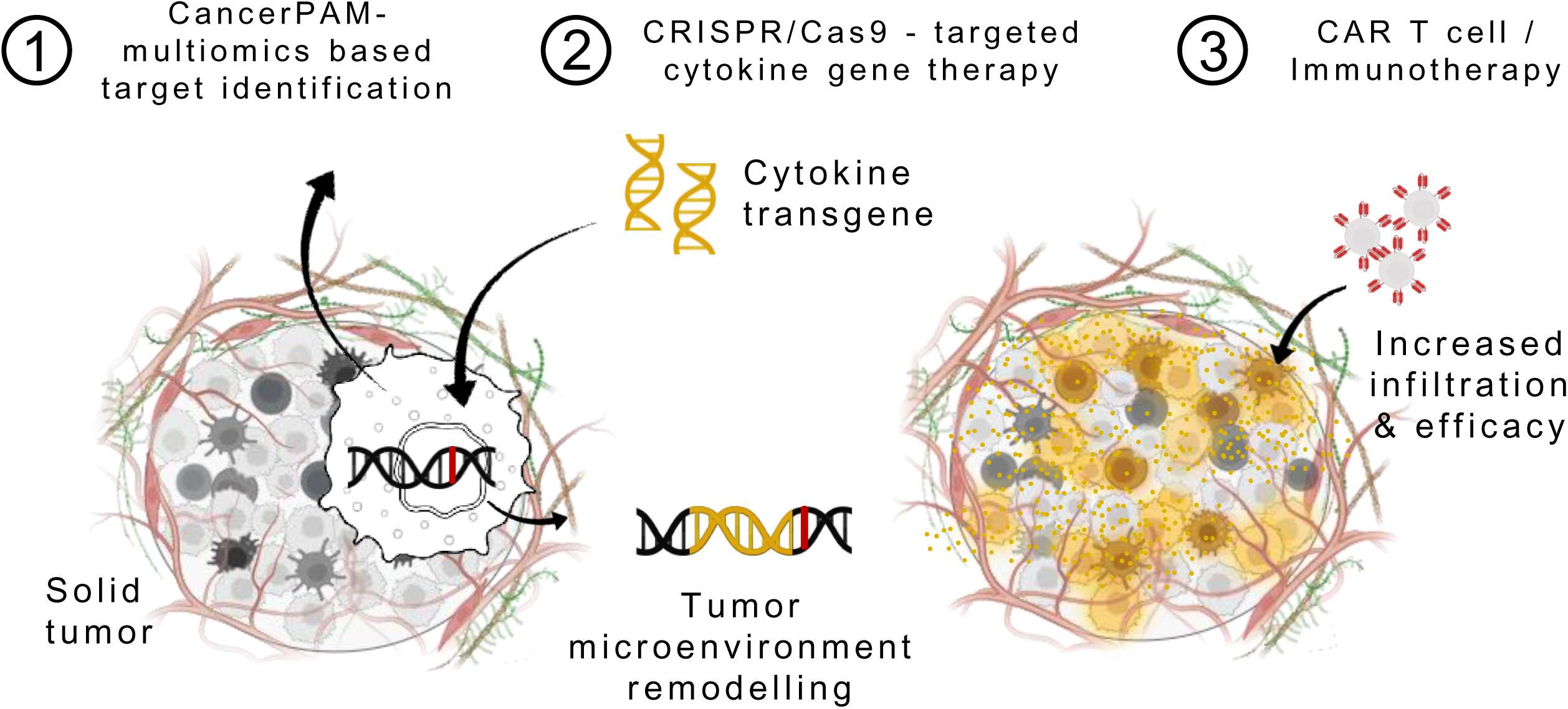
Therapeutic concept. Schematic of an augmentative approach combining the CancerPAM multiomics-based target identification pipeline with CRISPR/Cas9-targeted cytokine gene therapy, followed by CAR T cell or other immunotherapies. CancerPAM enables the identification of promising tumor-specific CRISPR knock-in target sites in solid tumors with an immunosuppressive microenvironment. At these sites, cytokine transgenes are integrated, leading to beneficial tumor biology changes that enhance subsequent CAR T cell or other immunotherapies.

## Results

*The CancerPAM Pipeline Identifies and Ranks Tumor-Specific Novel PAM Sites* Identification of tumor-specific CRISPR target sites is crucial to develop precise and efficient gene-editing strategies. We established a manual step-by-step process to identify and select single-nucleotide variants (SNVs) in tumor sequencing data that form novel 5’-NGG-3’ sequences, where N is any base, as protospacer adjacent motifs (PAM) recognized by the Cas9 nuclease (for simplicity, we refer to this specific type as *PAM site* throughout the manuscript) to effectively cleave target sequences in tumors. We tested this manual process on whole-exome sequencing (WES) datasets from neuroblastoma cell lines and identified promising novel PAM sites for subsequent knock-in experiments (**Supplementary Fig. S1a, Supplementary Table 1**). To automate and improve this selection process, we developed CancerPAM. CancerPAM is a Python-based modular pipeline that integrates WES- or whole-genome sequencing (WGS) data with multi-omics annotation to identify and rank novel PAM sites (**Fig. 2a**). The corresponding gRNA sequence is automatically determined, then the gene expression, copy number, gene dependency, Doench^5^ and Moreno^31^ CRISPR cutting efficiency scores, cutting frequency determination (CFD)^5^ and Massachusetts Institute of Technology (MIT)^32^ specificity scores are annotated. By incorporating these key biological and CRISPR-associated parameters, CancerPAM prioritizes optimal editing sites using both feasibility and safety criteria in a weighted ranking algorithm (**Supplementary Fig. S1b**). We hypothesized that several factors contribute to novel PAM site optimality, beyond gRNA sequence-based CRISPR efficiency scores. Specifically, we posited that higher gene expression, which correlates with chromatin accessibility^33^, and increased copy number would enhance knock-in efficiency. High specificity scores would reduce off-target activity, thereby improving safety, while low gene dependency would mitigate potential selection disadvantages. Higher expression levels of the target are also expected to reduce the risk of CRISPR editing causing pro-tumorigenic effects, such as inadvertently targeting a functional tumor suppressor gene - a risk already mitigated by the presence of the selected PAM-creating mutation, which may itself impair gene function. To implement these considerations, we applied a ranking algorithm that weighted the CFD specificity score twice to enhance safety. All other feature values are considered equally for ranking, except for the Doench and Moreno CRISPR cutting efficiency scores, which, given their limited predictive accuracy^34^, are compiled as a single predictor value (mean of both scores). Applying CancerPAM to neuroblastoma datasets, we identified a substantial number of novel PAM sites in both neuroblastoma cell lines and tumor samples from patients (matched tumor/healthy tissue data from diagnosis). In 14 neuroblastoma cell lines, a median of 327 novel PAM sites were identified from 1,470 SNVs (23% [95% CI: 22–24%]) in exonic regions. In tumor data from 54 patients, a median of 12 novel PAMs were identified from 82 SNVs (17% [95% CI: 13–22%]) in exonic regions, with 130 from 1,190 SNVs (14% [95% CI: 12–15%]) across the entire transcriptome (**Fig. 2b, Supplementary Note 1**). To evaluate CancerPAM pipeline accuracy and sensitivity, we conducted manual curation of findings for five neuroblastoma cell lines and cross-validated CancerPAM predictions. The pipeline demonstrated 99% accuracy, with nearly all manually identified novel PAM sites being successfully detected, confirming its reliability (**Fig. 2c**). Consistent with exonic data analysis, CancerPAM results revealed that tumor-specific PAMs were unevenly distributed across the genome, clustering in gene-dense regions (i.e. chromosomes 11, 17 and 19; **Fig. S1d, Supplementary Fig. S2f, Supplementary Notes 2 and 3**)^35^. CancerPAM accurately annotated key feature values, allowing data-driven selection of the most viable CRISPR target sites (**Fig. 2d, Supplementary Fig. S5a-g, Supplementary Table 2**). To validate the weighted ranking algorithm, we assessed correlations between PAM rank and various biological and CRISPR-associated features. Higher PAM rank strongly positively correlated with CRISPR specificity scores and moderately correlated with CRISPR efficiency scores and gene dependency (**Supplementary Fig. S6a-g**), but did not correlate with gene expression or copy number. Tumor sample groups bearing many or few PAM sites (above and below median) were compared regarding annotated features. Top-ranked PAM sites in tumor genomes harboring many PAM sites exhibited significantly higher specificity scores that strongly positively correlated with the total PAM site count in these tumor genomes (**Fig. 2e**). We determined a potential feasibility and safety threshold using the total novel PAM sites detected in a tumor sample. Tumor samples with at least 86 novel PAM sites (72% [39/54] of the cohort) had a >90% probability that their top three ranked PAM sites possessed a CFD specificity score >90 and MIT specificity score >80 (**Fig. 2f, Supplementary Notes 4 & 5**). To sum up, CancerPAM achieves 99% accuracy in identifying tumor-specific novel PAM sites from sequencing data, and uses feasibility and safety criteria to rank sites for optimal CRISPR target site prioritization.

**Fig. 2.**
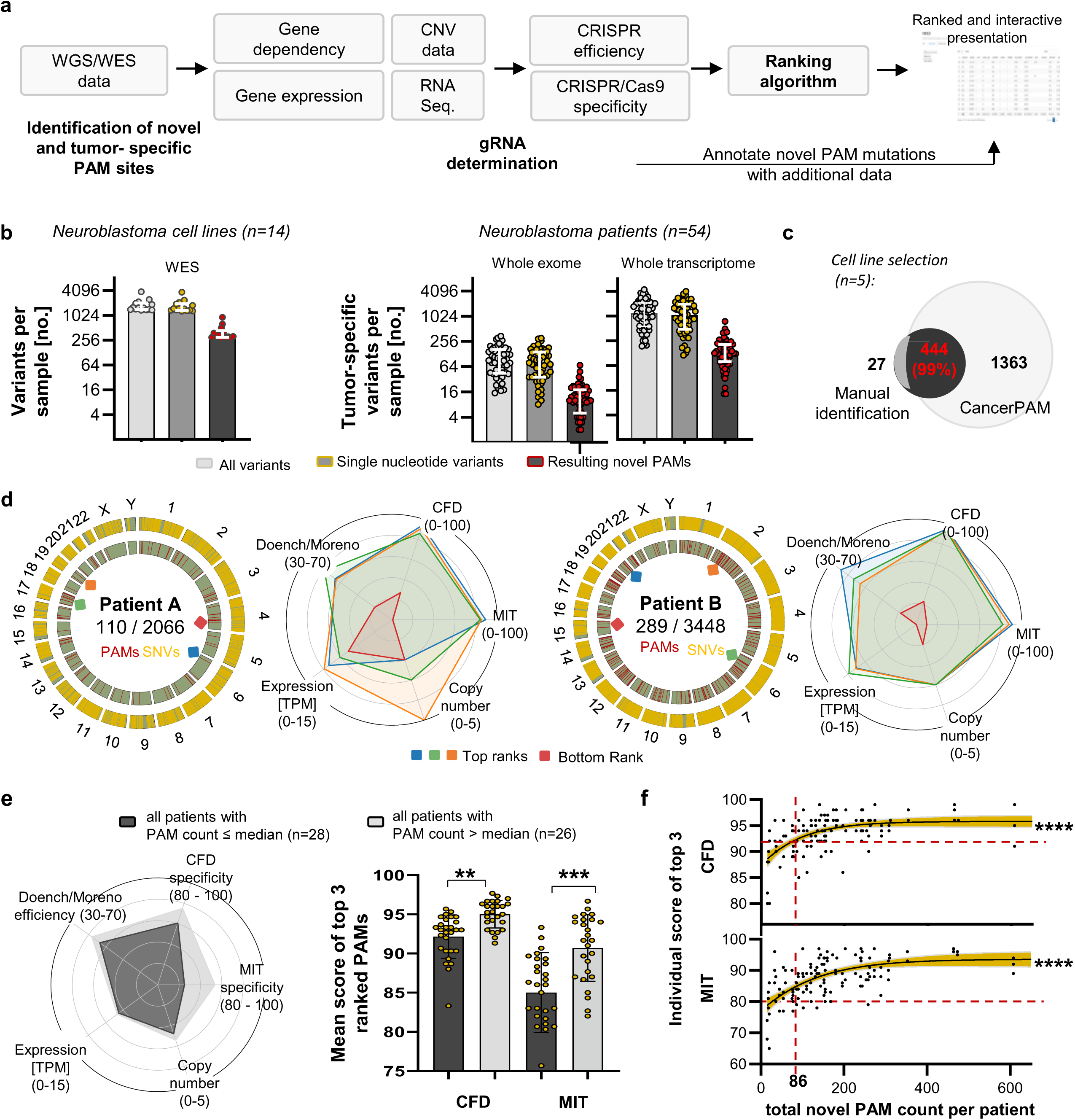
The CancerPAM multiomics-based automated pipeline identifies and ranks targetable tumor-specific PAM sites in cancer cell lines and patient samples. **(a)** Overview of the CancerPAM pipeline, which integrates whole-genome/whole-exome sequencing (WGS/WES) data analysis, gRNA identification, multiomics feature annotation, to identify and rank tumor-specific novel PAM sites. **(b)** Quantification of identified variants and tumor-specific PAM sites in neuroblastoma cell lines (from WES data) and neuroblastoma patients (from WGS data filtered for whole transcriptome or exonic variants). The *all variants* group includes insertions, deletions, InDels (<300 bp), and single nucleotide variants (SNVs). **(c)** Overlap between novel PAM sites identified by manual screening of five neuroblastoma cell lines and those identified using the CancerPAM pipeline. **(d)** Circos plots showing the chromosomal distribution of SNVs and novel tumor-specific PAM sites and radar plots visualizing annotated feature characteristics for the top three highest- and the lowest-ranked PAM site in two patients (A and B). **(e)** Circos plot illustrating mean annotated feature values of the top three ranked PAM sites, averaged across patients with a high (>median; n = 26) or low (≤median; n = 28) PAM count. Statistical comparison of these two groups is shown for the mean CFD and MIT specificity scores. **(f)** Correlation analysis of individual scores for the top three ranked novel PAM sites across patients (n = 162) versus total tumor-specific PAM count per patient. Red dashed lines indicate potential safety and feasibility thresholds. *Data presentation: (b,e) Means ± SD. Statistical tests: (e) Kruskal-Wallis with Dunn’s post hoc test, (f) logistic regression for curve fitting and Spearman correlation. p values: *<0.05, **<0.01, ***<0.001, ****<0.0001; n.s.: not significant*.

### CRISPR-Mediated Knock-In of Cytokine Transgenes at Tumor-Specific Target Sites is Feasible and Efficient

We selected two neuroblastoma cell lines (SK-N-BE2c: *MYCN*-amplified, SK-N-AS lacking *MYCN* amplification) in which to evaluate the feasibility of site-specific cytokine transgene integration. The most promising 9 novel PAM sites identified during manual selection were chosen, and also identified by CancerPAM **(Fig. 3a, Supplementary Fig. S1a, Supplementary Table 1)**. Sanger sequencing confirmed the presence of the selected novel PAM sites in 7 cases, while also verifying target site absence in the respective cell line where it was not present in WES data. Mutation frequencies in the respective cell line (harboring the novel PAM site) calculated from conformational Sanger sequencing varied between 0-20% for novel PAM sites located in *SH3BP1* and *SNX18*; 30–60% for *CHD1, RBM12, SCAF11, AP1M1* and *CHST11*; 66% for *RPLP0* (2 of 3 alleles) and 100% for *IGSF9B* (2 of 2 alleles) **(Fig. 3c)**. These novel PAM sites were targeted with specific Cas9/gRNA ribonucleoproteins (RNP) (**Supplementary Note 6**) while co-delivering homology-directed repair template (HDRT) for CXCL10, CXCL11 or IFNG transgene knock-in (**Fig. 3b, Supplementary Fig. S9c,d**). Cytokine transgenes contained a custom-designed EF1a-derived shortened promoter and Q8 epitope tag for flow cytometric detection (**Supplementary Fig. S10a, Supplementary Note 7**). Successful knock-in at the target sites was flow cytometrically confirmed 28 days after RNP/HDRT transfer **(Fig. 3d)**, and was most efficient at the *IGSF9B* locus in SK-N-BE2c and *RPLP0* locus in SK-N-AS (**Supplementary Note 8**). With the exception of the *IGSF9B* locus (knock-in occurred in both cell lines), gRNAs facilitated relevant knock-in only in cell lines harboring the novel PAM mutation (**Fig. 3d**). An alternative PAM site was present adjacent to the novel PAM site in *IGSF9B* in the SK-N-AS cell line, which lacks the novel *IGSF9B* PAM site, potentially explaining the unexpected integration events. Knock-in efficiency strongly positively correlated with copy number and gene expression, and weakly positively correlated with Doench and Moreno CRISPR efficiency scores (**Fig. 3e**). Site-specific cytokine transgene knock-in was validated by digital PCR. Specificity was confirmed and quantified using In/In and Out/In fluorescent probe-based assays that distinguish precise knock-in from random integrations or free-floating DNA. The results demonstrated robust integration of CXCL10, CXCL11 and IFNG at target loci (**Fig. 4a-c, Supplementary Fig. S12a, Supplementary Note 9**). IFNG, however, was integrated at significantly lower rates (**Fig. 4d**), in line with flow cytometry findings 28 days after knock-in (**Supplementary Fig. S10c**). Culturing untreated SK-N-BE2c cells with IFNG showed signs of IFNG-mediated toxicity and growth impairment (**Supplementary Fig. S12d**), which could indicate early cell loss following IFNG knock-in (before DNA isolation for digital PCR) or impairment of homology directed repair (HDR). To further validate pipeline ranking and CRISPR specificity score predictive value, we performed a knock-in experiment using primary T cells isolated from two healthy donors and the same gRNA set. High-throughput flow cytometry demonstrated low, but significant, unspecific knock-in for gRNAs targeting the *IGSF9B*, *SCAF11*, *SH3BP1*, *AP1M1* and *RPLP0* sites. No unspecific knock-in was observed for gRNAs targeting the CHD1, RBM12, SNX18 and CHST11 sites. (**Fig. 4e,f, Supplementary Fig. S13a, Supplementary Note 10**). However, only unspecific knock-in in the cases of *IGSF9B* and *RPLP0* resulted in CXCL10 levels exceeding variable baseline levels (**Supplementary Fig. S13b**). While not statistically significant, correlation analysis revealed a trend toward inverse correlation between unspecific knock-in rates and CancerPAM ranking scores. The highest-ranked PAM sites demonstrated the lowest frequency of off-target integration, further supporting the utility of the CancerPAM algorithm in prioritizing safe and effective gene-editing targets (**Fig. 4g,h**). In conclusion, successful CRISPR-mediated site-specific cytokine transgene integration was achieved and confirmed by digital PCR. Knock-in efficiency correlated strongly with gene expression and copy number, with minimal off-target effects. CancerPAM ranking was associated with reduced unspecific knock-in, validating its predictive accuracy for safe gene editing. Novel tumor-specific PAM site discovery and ranking by CancerPAM provide optimal support for experimental design and efficient CRISPR-mediated cytokine transgene integration.

**Fig. 3.**
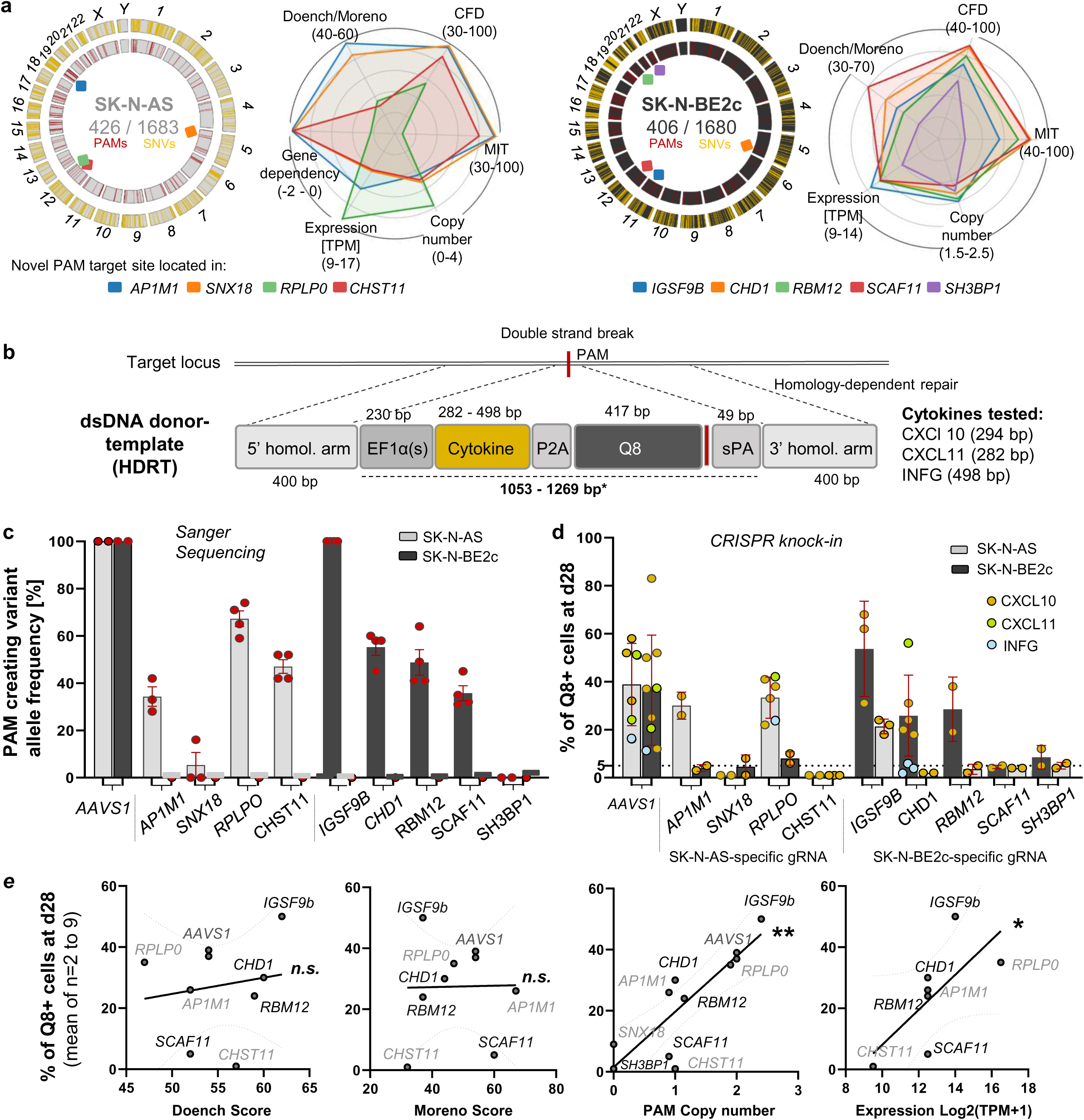
Gene therapeutic CRISPR knock-in of cytokine transgenes is efficient and specific for top-ranked novel PAM targets. **(a)** Circos plots showing the chromosomal distribution of SNVs (yellow) and novel PAM sites (red) in SK-N-BE2c and SK-N-AS neuroblastoma cell lines. PAMs tested in knock-in experiments are marked with colored squares. Radar plots visualize annotated features for these PAMs, named after their respective genes. **(b)** Schematic of CRISPR/Cas9-mediated transgene knock-in by homology-directed repair (HDR) following ribonucleoprotein (RNP) electroporation. The linear double-stranded (ds) homology-directed repair donor template (HDRT) consists of 5′ and 3′ homologous arms (400 bp), a custom EF1a-derived promoter, the cytokine transgene linked by a P2A self-cleaving peptide to a stainable Q8 reporter (CD34 epitope, CD8 transmembrane domain), followed by a stop codon and synthetic poly(A) (sPA) sequence. **(c)** PAM-creating mutation allele frequency analyzed by Sanger sequencing of selected targets in SK-N-BE2c and SK-N-AS. **(d)** Knock-in efficiency of three cytokines (CXCL10, CXCL11, IFNG) at different target sites, measured by Q8 antigen expression by flow cytometry at day 28 post-electroporation. **(e)** Correlation of knock-in rate (Q8 antigen expression at day 28) with PAM-annotated features, including CRISPR efficiency scores (Doench and Moreno), PAM copy number, and expression of the gene containing the PAM for selected targets. *Data presentation: (c, d) Means ± SD. Statistical analysis: (e) Linear regression for curve fitting and Spearman correlation. p values: *<0.05, *<0.01; n.s., not significant*.

**Fig. 4.**
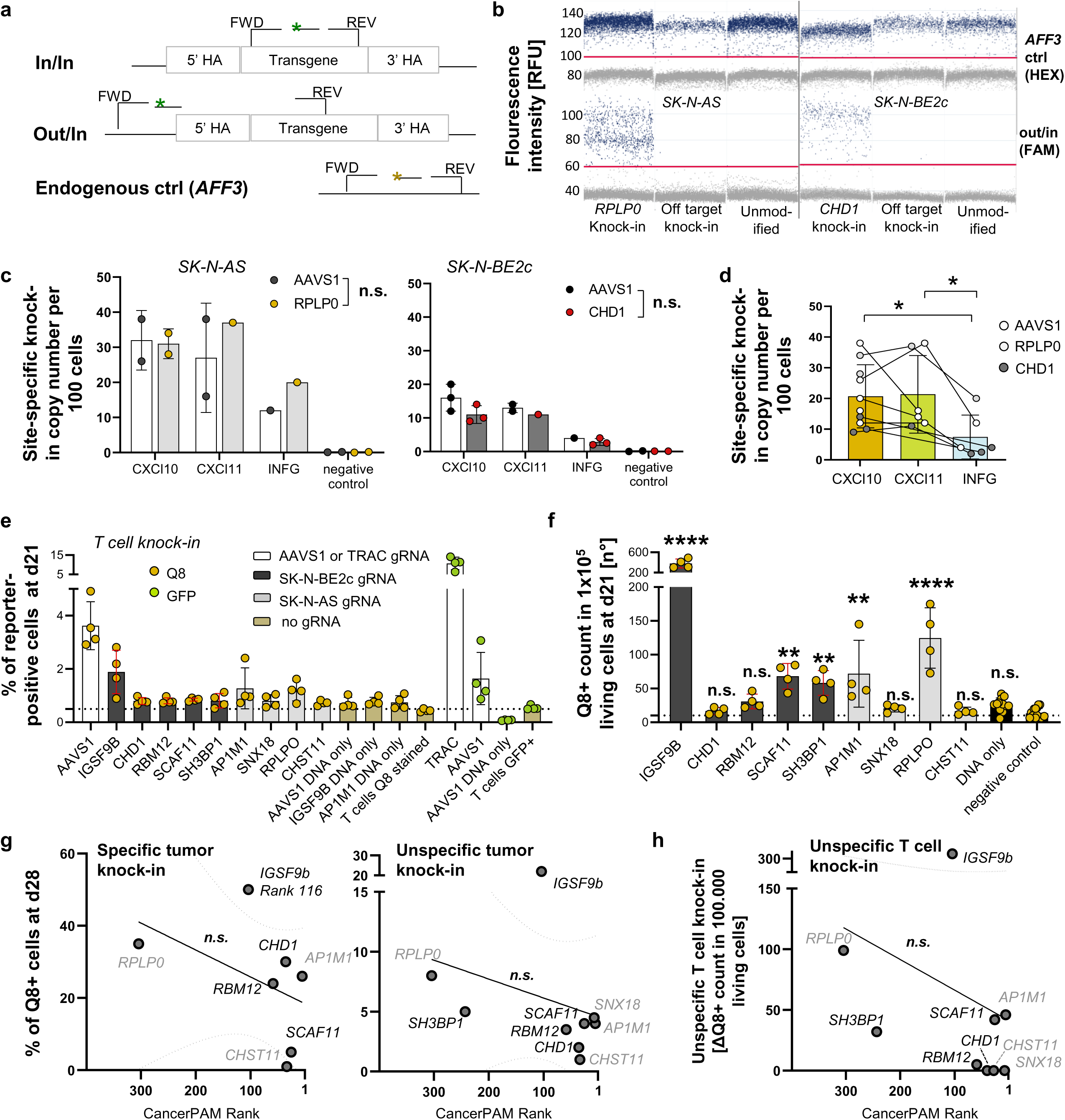
The CancerPAM pipeline accurately identifies novel PAM sites with a low unspecific knock-in risk. **(a)** Schematic of digital PCR assays for site-specific CRISPR knock-in confirmation. “In/In” refers to a fluorescence probe-based PCR assay with primers binding inside the transgene, while “Out/In” uses a forward primer upstream of the homologous arm. An AFF3 probe-based assay served as an endogenous control. **(b)** Digital PCR raw data showing positive partitions (blue) for knock-in and control samples using the Out/In or *AFF3* control assay. **(c)** Site-specific knock-in copy number per 100 cells, 7 days after RNP/HDRT electroporation in SK-N-AS and SK-N-BE2c for different transgenes and loci. **(d)** Cumulative site-specific knock-in copy numbers for CXCL10, CXCL11, and IFNG. **(e)** Comparison of reporter (Q8/GFP) positive cells 21 days post-CRISPR knock-in in primary T cells from two donors using gRNAs targeting neuroblastoma-specific novel PAM sites or AAVS1 and TRAC controls. **(f)** High-throughput flow cytometry (>300,000 cells analyzed per sample) to compare unspecific knock-in rates for Q8-reporter HDRTs at neuroblastoma-specific PAM sites. **(g)** Correlation of specific knock-in rates (in novel PAM-harboring neuroblastoma cells) and unspecific knock-in rates (in cells lacking the novel PAM) with CancerPAM-assigned ranks. **(h)** Correlation of unspecific CRISPR/Cas9-mediated T cell knock-in rates, as determined in (f), with CancerPAM ranks. *Data presentation: (c-f) Means ± SD. Statistical analysis: (c, d) Two-way ANOVA with Tukey test; (f) Kruskal-Wallis with Dunn’s post hoc test; (g, h) Linear regression for curve fitting and Spearman correlation. p values: *<0.05, **<0.01, ***<0.001, ***<0.0001; n.s., not significant*.

### Tumor Cytokine Secretion Achieved by CRISPR-mediated Knock-in Improves CAR T cell Infiltration and Efficacy in Vitro and in Vivo

To investigate the effects of cytokine secretion on CAR T cells, we enriched transgenic neuroblastoma cell lines using fluorescence-associated cell sorting (FACS) for PE-labelled Q8-reporter-positive cells. Flow cytometry and digital PCR (dPCR) confirmed stable transgene expression and site-specific knock-in, except in the case of IFNG into SK-N-BE2c (**Fig. 5a, Supplementary Fig. S14, Supplementary Note 11**). Elevated levels of CXCL10, CXCL11 and IFNG were confirmed in these cell lines by ELISA-based quantification (**Fig. 5b**). CAR T cells demonstrated significantly better tumor growth control over 96 hours compared to non-transduced cells (effector-to-target ratio, 1:5; cytokine-expressing and enriched SK-N-AS neuroblastoma cells co-cultured with L1CAM-targeting CAR T cells; **Fig. 5c**). However, complete tumor eradication was not achieved under any condition. Cytotoxicity in IFNG-expressing tumor cells was pronounced at 24 hours, but no longer statistically significant by 72 hours (**Fig. 5c, Supplementary Fig. S16a, Supplementary Note 12**). Conditioned media from CXCL10- and CXCL11-expressing tumor cells significantly enhanced CAR T cell migration compared to media from control tumor cells using two distinct transwell migration assays (**Supplementary Fig. S18a-e**). In vitro 3D bioprinted tumor models^36^ were used to further investigate transendothelial migration and tumor infiltration. A substantial increase in CAR T cell tumor infiltration into CXCL10- and CXCL11-expressing tumors, both in the presence and absence of an additional endothelial layer was confirmed in 12-hour infiltration experiments, after relevant CAR T cell proliferation (measured by KI67 positivity) within 12 hours post-co-culture initiation was excluded as a confounder in a preliminary experiment (**Fig. 5d, Supplementary Fig. S19c**). We conclude that tumor-secreted CXCL10 and CXCL11 enhances CAR T cell migration. In vivo luciferase-expressing L1CAM-targeting CAR T cell trafficking and expansion was monitored by longitudinal bioluminescence imaging (BLI) in mice subcutaneously xenografted with either transgenic or unmodified tumor cell lines in the flank. Tumors with low (SK-N-AS) or high (SK-N-BE2c) L1CAM antigen expression were evaluated (**Fig. 6a, Supplementary Figs. S20-S23, Supplementary Note 13**). Overall tumor-localized CAR T cell mass - including infiltrated and expanded T cells - was quantified as the flank BLI area under the curve (AUC) from day 1 until the final BLI measurement prior to sacrifice. A positive trend in tumor-localized CAR T cell mass was observed for mice bearing cytokine-expressing tumors compared to those with unmodified tumors (+37% in SK-N-AS and +18% in SK-N-BE2c), although this did not reach statistical significance. Notably, mice bearing cytokine-expressing tumors exhibited significantly higher BLI signals at day 4 post-injection, indicative of enhanced CAR T cell infiltration (SK-N-AS: +221%, *p*=0.026; SK-N-BE2c: +132%, *p*=0.038; **Fig. 6b,c, Supplementary Fig. S24c-d)**. In terms of therapeutic efficacy, no tumor remission was observed in mice bearing L1CAM-low (SK-N-AS) tumors, regardless of CXCL10 expression **(Fig. 6d)**. CAR T cell efficacy and tumor control in mice bearing L1CAM-high (SK-N-BE2c) tumors, however, varied across treatment groups. L1CAM-knockout tumors rapidly grew by day 11 (median survival: 7 days), while partial remission occurred in three animals bearing unmodified tumors, (assessed until day 22, median survival: 11 days; **Fig. 6d, Supplementary Fig. S24e**). Regression was enhanced for CXCL10-secreting tumors, with three animals being tumor-free by day 22 then relapsing after day 26, as the CAR T cell BLI signal disappeared (median survival: 14 days, i.e. +27% compared to unmodified tumor harboring animals; **Fig. 6d, Supplementary Figs. S20a,b,e, Supplementary Note 14**). Overall, our in vitro and in vivo experiments showed that tumor-secreted CXCL10 enhanced CAR T cell migration and tumor infiltration.

**Fig. 5.**
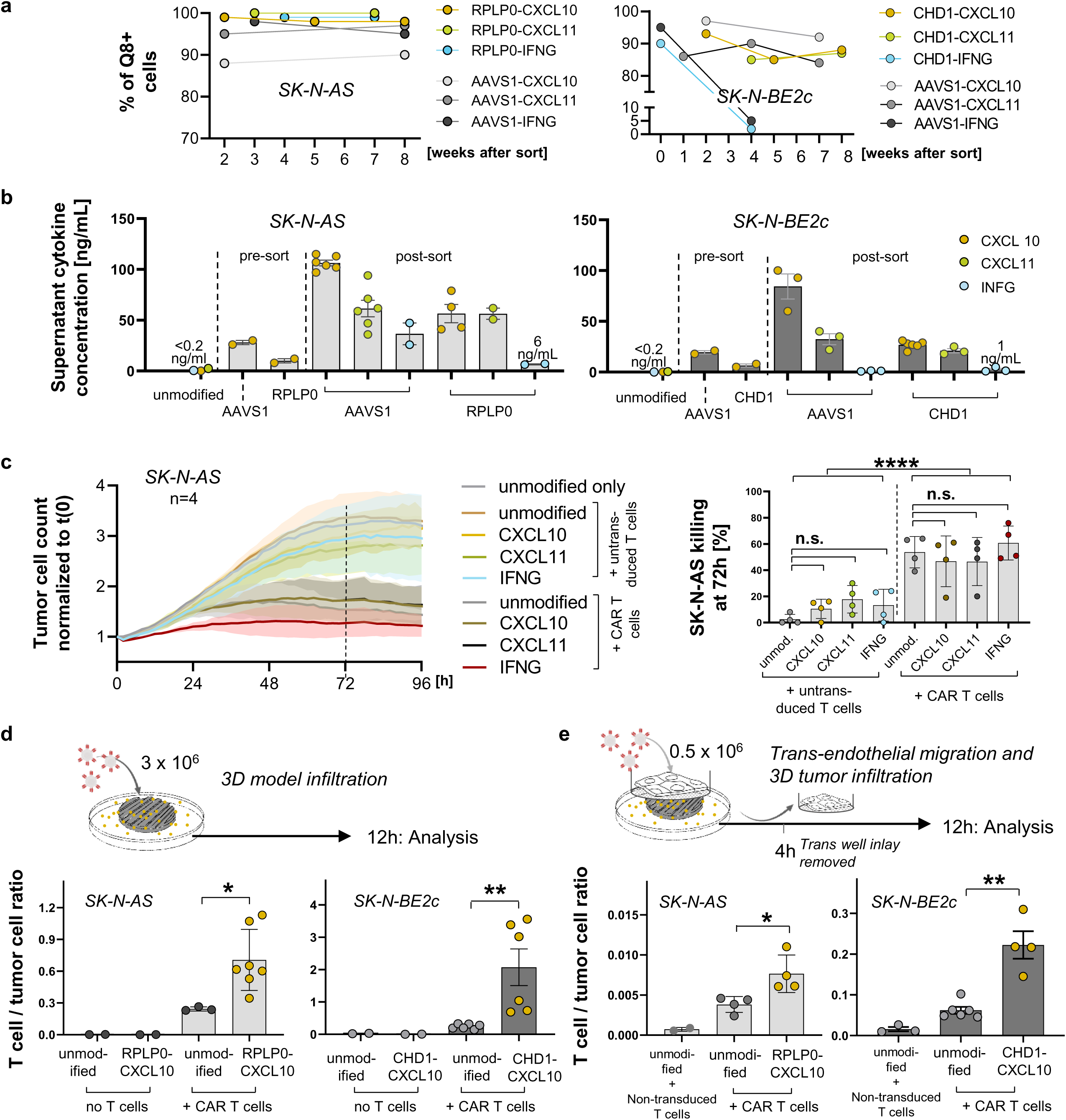
Gene therapeutically induced tumoral CXCL10 secretion increases CAR T cell migration and infiltration in vitro. **(a)** Post-enrichment transgene expression rates measured by Q8 positivity in flow cytometry over 8 weeks for transgenic SK-N-AS and SK-N-BE2c cell lines. **(b)** Supernatant cytokine concentrations for different cytokine-target locus combinations before (pre-sort) and after enrichment (post-sort) for Q8L cells, determined by ELISA. **(c)** CAR T cell killing dynamics for transgenic and enriched cytokine-expressing SK-N-AS cell lines co-cultivated with either non-transduced T cells or L1CAM-targeting CD4L and CD8L CAR T cells at an effector-to-target ratio of 1:5. Killing dynamics were tracked over 96 hours by Incucyte live imaging. The dotted line marks 72 hours, where statistical comparisons were conducted. Killing at 72 hours was determined as the ratio of the t(0) normalized tumor cell count of the treated cell line against the untreated unmodified control. **(d)** Bioprinted 3D neuroblastoma models were used to analyze CAR T cell infiltration into 3D tumors 12 hours post-co-culture. 3D tumor CAR T cell infiltration, represented by the T cell-to-tumor cell ratio (flow cytometry), was compared between enriched cytokine-expressing and unmodified SK-N-AS and SK-N-BE2c cells. (**e)** Trans-endothelial migration and 3D tumor infiltration assays were performed using a HUVEC monolayer with a Boyden transwell insert on a bioprinted 3D neuroblastoma model. Four hours after adding L1CAM-targeting CAR T cells, the insert was removed, and 3D tumor infiltration was measured by flow cytometry 8 hours later, following the same procedure as in (d). *Data presentation: (b-g) Means ± SD. Statistical analysis: (b) Logistic regression curve fitting; (b, d) Kruskal-Wallis with Dunn’s post hoc test; (d) Two-way ANOVA for CAR T cell vs. non-transduced T cell comparison; (e, f) Mann-Whitney test. p values: *<0.05, **<0.01; n.s., not significant*.

**Fig. 6.**
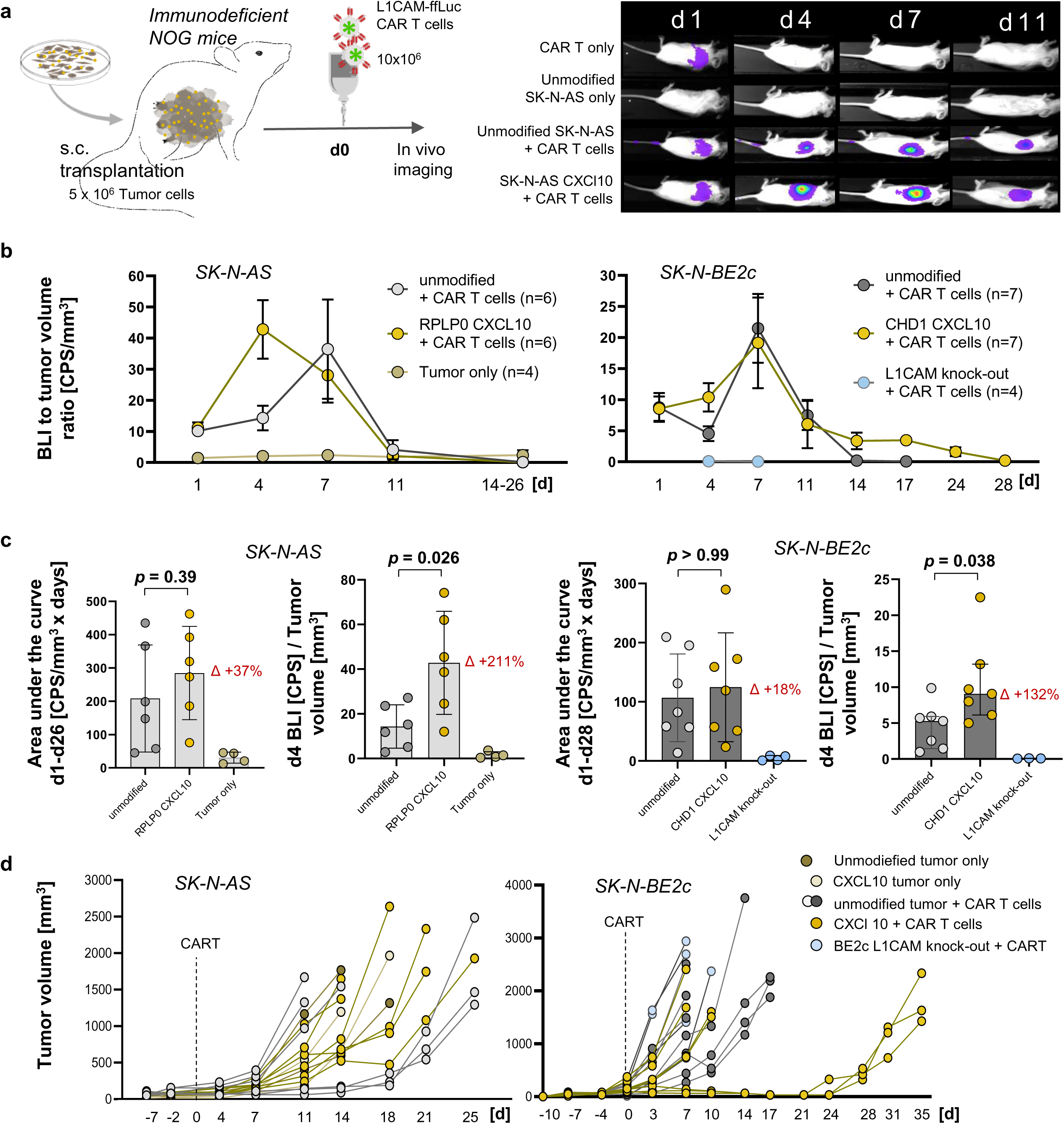
Gene therapeutically induced tumoral CXCL10 secretion increases CAR T cell tumor infiltration in vivo. **(a)** Schematic of the xenograft mouse model using immunodeficient NOG mice with subcutaneous transplantation of unmodified SK-N-AS and SK-N-BE2c cells, RPLP0/CHD1 CXCL10 Q8L enriched knock-in cell lines, or a SK-N-BE2c L1CAM knock-out cell line in the left flank. After tumor engraftment, mice were treated with firefly luciferase-expressing L1CAM-targeting CD3L CAR T cells by tail vein injection, followed by repetitive bioluminescence imaging analysis. **(b)** Tumoral CAR T cell infiltration over time, represented as the bioluminescence signal in the left flank relative to the tumor size measured on the same day for different treatment groups. The *SK-N-AS tumor only group* includes four animals that were excluded from CAR T cell treatment due to tumor sizes smaller than 50 mm³ (Fig. S24a). **(c)** Comparison of total tumoral CAR T cell infiltration and expansion, represented by the area under the curve (AUC) of the bioluminescence-to-tumor volume ratio, and comparison of early infiltration based on the bioluminescence-to-tumor volume ratio on day 4 between treatment groups. **(d)** Individual tumor growth curves for all treatment groups. *Data presentation: (b, c) Means ± SD. Statistical analysis: (c) Mann-Whitney test*.

## Discussion

We present a novel gene therapeutic strategy that remodels the tumor microenvironment (TME) through tumor-intrinsic cytokine expression to enhance immune cell infiltration and immunotherapy efficacy, enabled by CancerPAM - a bioinformatics pipeline that identifies patient-specific, tumor-selective CRISPR knock-in target sites. CancerPAM leverages multi-omics datasets to systematically select optimal CRISPR targets, prioritizing specificity and safety while facilitating efficient transgene integration. Applied to data from neuroblastoma samples, with a low mutational tumor burden relative to adult malignancies^37,38^, CancerPAM identified a median of 130 tumor-specific PAM sites across the entire transcriptome. Using these data, we successfully implemented CRISPR-mediated insertion of pro-inflammatory cytokine genes (CXCL10, CXCL11, IFNG) to enhance immune cell infiltration and function. CancerPAM’s integrative approach, incorporating genomic, transcriptomic and epigenomic data, provides an automated and systematic method to identify tumor-specific PAM sites and effectively minimize off-target risks. The pipeline substantially accelerates the identification process while maintaining high accuracy compared to manual selection. Its ranking algorithm prioritizes sites based on specificity, predicted efficiency and biological relevance to ensure robust therapeutic target selection. Certain limitations remain. Sequencing dataset biases and tumor heterogeneity may influence reproducibility and the CRISPR efficiency scores used require further improvement. CancerPAM is currently optimized for 5’-NGG-3’ PAM sequences (related to the *Streptococcus pyogenes*-derived Cas9 used), limiting compatibility with alternative Cas9 variants. Future pipeline iterations will focus to broaden PAM recognition, refining ranking criteria and further integrating functional validation.

CAR T cell recruitment and tumor control in vivo were enhanced through CXCL10 expression by engineered tumor cells. These findings highlight tumor-intrinsic cytokine expression as a promising strategy to reprogram the TME and overcome key barriers to immunotherapy in solid tumors. By enabling localized, sustained secretion of chemokines such as CXCL10 and CXCL11 directly from tumor cells, this approach promotes immune cell recruitment and trafficking while avoiding systemic cytokine exposure, potentially minimizing off-target effects and toxicity. Unlike conventional systemic cytokine therapies or exogenous TME modulation, this strategy reprograms the tumor from within, enhancing its susceptibility to immunotherapy. In combination with CAR T cell therapy, we observed improved immune infiltration and tumor control in vitro and in vivo, supporting this synergistic paradigm. While our study focuses on neuroblastoma, the underlying principle may be applicable across a range of immune-excluded solid tumors. Future work should explore multiplexed or inducible cytokine circuits and combinations with checkpoint blockade or antigen-unmasking strategies to further enhance this therapeutic platform.

Clinical translation of our strategy depends on improving CRISPR knock-in efficiency and tissue-specific in vivo delivery. Optimized non-viral knock-in approaches, such as modified single-stranded DNA templates and DNA nanostructures have been validated to significantly enhance homology-directed repair efficiency^39^. Novel CRISPR delivery strategies like lipid nanoparticles^40^, evolved viral particles^41^ or peptide-based delivery of CRISPR components^42^, show promise for in vivo translation. Particularly, lipid nanoparticle-mediated mRNA delivery has been shown to allow transient CRISPR expression and selective organ-targeting for tissue-specific delivery^40,43,44^. Adeno-associated viral (AAV) vectors, widely used for stable gene delivery, require further refinement to mitigate immunogenicity and insertional mutagenesis risks^45^. Combining AAV and nanoparticle-based approaches could enhance safety and efficiency^46^. Notably, in vivo administration of CAR T cell-encoding mRNA lipid nanoparticles enabled functional CAR T cell production without ex vivo engineering, suggesting a potential adaptation for CRISPR-based TME remodeling to improve immunotherapy in solid tumors^47^.

Our approach—combining CancerPAM-guided CRISPR editing with tumor-intrinsic cytokine expression—offers a versatile and conceptually novel framework to enhance immunotherapy across solid tumors. By tailoring immune-stimulatory cues within the tumor itself, this strategy has the potential to convert immune-excluded tumors into immune-permissive environments, thereby augmenting existing therapies such as CAR T cells or immune checkpoint inhibitors. While our proof-of-concept focuses on neuroblastoma, the platform is broadly applicable to other malignancies with immunosuppressive TMEs. Continued advances in CRISPR delivery technologies and knock-in efficiency will be essential to realizing clinical translation. Nevertheless, our findings establish a foundation for precision-engineered TME modulation, highlighting the synergy between computational target selection and therapeutic genome engineering to unlock new avenues in personalized cancer immunotherapy.

## Material & Methods

### CancerPAM - Data Sources and Sample Information

Unpublished data from several neuroblastoma cell lines (terminate NB, n=14) and a patient cohort (n=54) with respective disease information were analyzed from our own collaborative research network. Whole exome sequencing was performed for the cell lines, while whole genome sequencing was performed for the patients for tumor and healthy tissue. Expression data was generated for both cell lines and patients from RNA sequencing data. Additionally, public sequencing data from neuroblastoma cell lines (n=48) from the Cancer Dependency Map Portal (DepMap 23Q4, 22Q2, 20Q4) were included in the analysis^3,48–52^. These datasets consist of mutational data generated through high-coverage sequencing, read-depth analysis and RNA sequencing (WES) with regular new releases. Gene dependency data, gene expression and copy number was obtained from DepMap as well. Somatic variants were identified using MuTect2 (version) to call variants between tumor-normal pairs for patients and against the reference genome GRCh38.p14 (release 44) for DepMap and terminate NB cell lines. For comprehensive gene annotation, the reference genome GRCh38.p14 (release 44) was used for all data sets.

### CancerPAM - Identification and Analysis of Novel PAMs

CancerPAM (Python) was designed to process the called variants (csv-files) based on their genome builds (e.g., hg38) and identify somatic variants that produce novel NGG PAMs. The 5′ and 3′ genomic sequences surrounding the somatic variants were retrieved through an API and the UCSC Genome Browser. This analysis was conducted to assess whether novel Cs were adjacent to existing Cs or novel Gs to existing Gs. The output included information on the somatic variant, the potential sgRNA sequence, the novel PAM and whether the novel PAM was located on the plus or minus strand of the genome. Novel PAMs were analyzed for CRISPR efficiency and specificity using CRISPOR^53^. Access to the platform was facilitated through an API, which provided the relevant efficiency and specificity values. CancerPAM was designed using Snakemake to automate and streamline the analysis pipeline^54^.

### CancerPAM - Features and Ranking of Novel PAMs

In this study, four CRISPOR-derived scores were utilized alongside gene dependency, expression and copy number data. Two scores, MIT and CFD, assess on-target specificity by estimating the guide RNA’s genomic uniqueness and considering mismatches with the target DNA. These scores, ranging from 0 to 100, account for the number, position and distribution of mismatches in a sequence-dependent manner, with higher values indicating reduced off-target effects. Notably, the CFD score correlates more strongly with the total off-target cleavage fraction of a guide than the MIT score^5,32,55^. To predict the on-target efficiency the Doench and Moreno-Mateos scores were chosen. The higher the efficiency score, the more likely cleavage occurs at this position. The Moreno-Mateos score is based on CRISPRscan and predicts gRNA activity by analyzing molecular features such as guanine enrichment, adenine depletion, nucleotide truncation and 5’ mismatches, effectively capturing the sequence determinants influencing CRISPR/Cas9 activity in vivo^31^. The Doench score, developed from a large-scale CRISPR study, evaluates sequence traits linked to high or low guide RNA activity, creating a scoring algorithm based on desirable nucleotide patterns. These scores also range from 0 - 100^5^. Beside that gene dependency data was used to estimate how essential a gene is for cell survival and the lethality of its knock-out. Highly negative dependency scores indicate critical gene functions, with values below -0.5 signifying depletion in most cell lines and scores below -1 representing strong lethality, corresponding to the median of all essential genes. A value of 0 reflects non-essential genes^3^. To calculate a single score for each novel PAM, they were first ranked for each feature individually. PAMs lacking data for a given feature were assigned the lowest rank for that feature. In the case of ties in feature values, the corresponding PAMs shared the same rank. Subsequently a final score for each novel PAM site was determined by summing the weighted ranks of all features. The Doench and Moreno scores (which assess cutting efficiency but not knock-in efficiency) were averaged. In contrast, the CFD score was assigned a weighting factor of 2, to increase specificity and hence safety. The final score was calculated as follows: 2* rank CFD + rank MIT + 0.5*(rank Doench + rank Moreno) + rank Dependency + rank expression + rank copy number. Scores were then normalized within each dataset to a range of 0–100, where a score of 100 represents the most promising target and a score of 0 the least promising. Finally, PAMs were ranked from highest to lowest score in the output table. In the case of ties the corresponding PAMs shared the same rank.

### Cell Lines and Cell Culture

Neuroblastoma cell lines SK-N-BE(2)c and SK-N-AS cells were obtained from Prof. Michael Claus V Jensen at Seattle Children’s Hospital. HEK293T cells (ATCC CRL-3216) and neuroblastoma cell lines were cultured in DMEM high glucose medium (Gibco) with 10% heat-inactivated fetal calf serum (FCS) and 1% Penicillin/Streptomycin (Pen/Strep, Gibco). Human Umbilical Vein Endothelial Cells (HUVECs, ATCC CRL-1730) were cultured in Endothelial Cell Growth Medium 2 (PromoCell, Cat# C-22111/39211), supplemented with EGF, IGF, Vascular Endothelial Growth Factor (VEGF), FGF2, 2% Fetal Bovine Serum (FBS) and 1% Pen/Strep. All cell lines were maintained at 37°C with 5% CO2. Lymphocytes were isolated from fresh primary adult blood cells from anonymous healthy human donors using EasySep isolation kits for CD3+, CD4+ or CD8+ T cells (StemCell Technologies). Isolated T cells were cultured at an initial density of 10^6^ cells per ml in X-VIVO 15 medium (Lonza) supplemented with human serum (5%, Gemini), penicillin–streptomycin (1%, Gibco), interleukin (IL-7; 5LngL/ml, Miltenyi) and IL-15 (5LngL/ml, Miltenyi). After isolation, cells were stimulated for 2 days with anti-human CD3/CD28 magnetic Dynabeads (Thermo Fisher Scientific) using a 1:1 bead-to-cell ratio.

### Homology-Directed Repair Template Design and Preparation

Sequences for all HDRTs used are provided in **Supplementary Table 3**. Human cytokine coding sequences were obtained from the NCBI genome viewer (https://www.ncbi.nlm.nih.gov/gdv/). Q8 tag and sPA were used as published^56,57^. DNA fragments were acquired from Twist Bioscience after codon optimization. We used homology-based seamless cloning NEBuilder HiFi DNA Assembly, NEB) to insert DNA fragments into a high-copy-number pUC-based plasmid vector, containing a ColE1 origin and Ampicillin resistance (AmpR) before transformation of ultracompetent XL10-Gold (Agilent) E-coli cells. Transformed cells were selected on LB agar plates containing ampicillin. Positive clones were identified by colony PCR and their plasmid DNA was isolated using the ZymoPURE Plasmid Miniprep Kit (ZYMO RESEARCH) before Sanger sequencing (LGC Genomics) was conducted for comprehensive analysis. Linear dsDNA homology-directed repair templates (HDRTs) were produced by PCR amplification using KAPA HiFi DNA Polymerase (Roche) in a 400 μl reaction. Following amplification, the HDRTs were purified using AMPure XP Beads (Beckman Coulter) and their correct size confirmed by gel electrophoresis. For vector expression testing the Effectene transfection reagent (Qiagen) was used in HEK293T cells at a dose of 1µg DNA per 1x10^6^ cells.

### CRISPR Cutting and Knock-in

CRISPR cutting and knock-in experiments were conducted using the Lonza 4D-Nucleofector electroporation system with 16-well Nucleocuvette™ strips as described previously^58^. Ribonucleoprotein (RNP) complexes were assembled using chemically synthesized single-guide RNAs (sgRNAs) (Synthego) and Alt-R™ S.p. Cas9 Nuclease V3 (IDT) with or without homology-directed repair templates (HDRTs) and polyethylene glycol (PGA). Electroporation was performed using the X-unit of the 4D-Nucleofector (Lonza, Basel, Switzerland), applying pulse programs DN110 for SK-N-BE2c and FF104 for SK-N-AS cells. For CRISPR-Cas knock-in experiments, the RNP complex was assembled immediately before nucleofection. Per reaction, a gRNA to Cas9 protein molar ratio of 2:1 was used (80 pmol gRNA (2,5 µg) and 40 pmol Cas9 enzyme (6,67 μg) per 1x10^6^ cells) with 50 μg polyethylene glycol (PGA) were mixed with sterile water. The complex was incubated at room temperature for 15 minutes. HDRT dsDNA (2.0 μg per 1x10^6^ tumor cells and 1.0 µg per 1x10^6^ T cells) was then added. Tumor cells were trypsinized, counted, washed twice (200 × g, 5 minutes, room temperature) and resuspended in SF buffer containing Supplement 1 (1:4.5 dilution). T cells were resuspended in supplement containing buffer P3. For each reaction, 20 μL of the cell suspension was added to the corresponding RNP-HDRT mixture and electroporation was carried out using Lonza 4D-Nucleofector 16-well strips. Following nucleofection, cells were recovered in antibiotic-free RPMI medium supplemented with 10% FBS, incubated at 37°C with 5% COL and transferred to appropriate cell culture plates. For T cell knock-ins Alt-R HDR Enhancer V2 (IDT) at 1 µM final concentration was added. CRISPR knock-in experiments targeting tumor cells did not include an Alt-R HDR Enhancer V2 (IDT). After 24 hours, culture conditions were adjusted as necessary, including the removal of HDR Enhancer (IDT) when applicable. For CRISPR editing efficiency assessment, custom PCR assays were designed using the GeneGlobe tool (Qiagen) to amplify the cut sites. Out/out PCR reactions were performed using the QIAprep & CRISPR Kit (Qiagen) with AllTaq Master Mix. Sanger sequencing was conducted and the resulting chromatograms were analyzed using the ICE (Synthego) and EditR (http://baseeditr.com/) tools to quantify indel formation.

### Digital PCR for Copy Number Variation Analysis

To assess the efficiency and stability of genetic modifications in transgenic cell lines, digital PCR (dPCR) was performed using the QIAcuity digital PCR system (QIAGEN). All primer and probe sequences are available in **Supplementary Table 4**. Copy number variation (CNV) analysis was conducted using a duplex probe-based approach to quantify the integrated transgenes relative to a reference gene. Genomic DNA was extracted from stable transgenic and wild-type control cell lines. The dPCR reaction mix contained the QIAcuity Probe PCR Master Mix (QIAGEN), gene-specific primers and probes and fragmented template DNA. The total reaction volume was 12 µL for a 96-well nanoplate format and 40 µL for a 24-well format. The final reaction composition included: 1× QIAcuity Probe PCR Master Mix, 800 nM forward and reverse primers for the target transgene, 400 nM for the reference gene (AFF3) primers, 400 nM target-specific hydrolysis probe (FAM-labeled) and 200 nM AFF3 probe (HEX-labeled). To optimize template accessibility, 0.05 U/µL XbaI (New England Biolabs) was added. Samples were pipetted into the QIAcuity Nanoplate (QIAGEN) and sealed with a QIAcuity Nanoplate Seal (QIAGEN). The dPCR thermal cycling protocol was performed on the QIAcuity One digital PCR system (QIAGEN) with the following conditions for In/In transgene-specific assays: Enzyme activation at 95°C for 2 minutes, denaturation at 95°C for 15 seconds, primer annealing and elongation at 58°C for 30 seconds. This cycle was repeated for 40 cycles, followed by an imaging step for fluorescent signal detection. For Out/In site-specific knock-in assays annealing at 58°C for 30 seconds was followed by an additional elongation step for 1 minute at 72° and cycle count was increased to 55 cycles followed by a 2-minute extension at 72°C to enhance signal resolution. Following amplification, the QIAcuity software (QIAGEN) analyzed partitioned fluorescence signals to determine absolute DNA copy numbers per microliter using Poisson statistics. Each sample was processed as a duplex reaction, normalizing transgene copy number to the AFF3 reference to account for variations in DNA input and to calculate cell counts. The software generated graphical representations of positive and negative partitions, scatterplots and quantitative tables. Negative control wells (no-template controls) were included to validate specificity and fluorescence thresholding was manually reviewed to ensure correct partition classification. Final CNV values were expressed as copies per 100 cells.

### CAR T Cell Generation

The generation of CAR T cells was conducted as described previously and under the ethical approval EA2/262/20 from Charité – Universitätsmedizin Berlin^18^. PBMCs were isolated from the blood of healthy donors using a density gradient centrifugation method. Briefly, blood was diluted 1:1 with phosphate-buffered saline (PBS) and carefully layered over Ficoll-Paque (Sigma-Aldrich) in a centrifuge tube. Following centrifugation at 300 × g for 20 minutes without brake, the mononuclear cell layer was collected. Red blood cells were lysed using a hemolysis buffer and the remaining cells were washed with PBS and counted. T cells were isolated from PBMCs the CD3+ Pan-T Cell Isolation Kit (Miltenyi), followed by magnetic-activated cell sorting (MACS) as per the manufacturer’s instructions. Isolated T cells were seeded at a concentration of 1 × 10^6^ in a 24-well plate and activated with anti-CD3/CD28 beads at a 1:1 cell-to-bead ratio. Depending on the chimeric antigen receptor (CAR) construct, T cells were transduced with lentiviral vectors (SIN epHIV7) propagated in 293T cells on Day 1 post-activation at a multiplicity of infection (MOI) of 1. non-transduced T cells served as negative controls. Transduced T cells were cultured in T cell medium (RPMI + 10% FBS + 1% GlutaMAX (Gibco) supplemented with 0.5 ng/mL interleukin-15 (IL-15, Miltenyi)) and 5 ng/mL interleukin-7 (IL-7, Miltenyi), with medium and cytokine replenishment every 2–3 days. Twelve days post-transduction, transduction efficiency was assessed by immunostaining for epidermal growth factor receptor truncated (EGFRt) and analyzed by flow cytometry. EGFRt-positive cells were enriched using MACS. Cells were stained with a PE-labeled anti-EGFRt antibody, followed by incubation with magnetic anti-PE beads and subsequently separated magnetically. The enriched T cells were then cryopreserved for future use. Prior to experimental use for in vitro experiments, cryopreserved CAR T cells were thawed and subjected to an expansion protocol. This involved co-culturing the CAR T cells with freshly thawed, irradiated (80 Gy) PBMCs and irradiated (35 Gy) EBV-transformed lymphoblastoid feeder cells (TM-LCL), in the presence of OKT3 (Miltenyi) CD3 activating antibody complex, IL-7 and IL-15. The culture medium was refreshed every 2–3 days with the addition of 0.5 ng/mL IL-15 and 5 ng/mL IL-7. Experiments were conducted 12–15 days after the initiation of the second stimulation. For in vivo use CAR T cells were not enriched using MACs and not but used directly after thawing.

### Cytotoxicity Assays

To assess the cytotoxic activity of L1CAM targeting CAR T cells against tumor cells, real-time imaging was performed using the IncuCyte® Live-Cell Analysis System (Sartorius). SK-N-AS transgenic or control tumor cells were seeded in 48-well plates at a density of 0.05 × 10L cells per well and cultured in RPMI + 10% FBS + 1% P/S for 16 hours prior to co-culture. To enable live-cell imaging, tumor cells were fluorescently labeled using Vybrant DiO Cell-Labeling Solution (Thermo Fisher) at 1:500 dilution for membrane staining and IncuCyte Nuclight Rapid Red Dye (Sartorius) at 1:4000 dilution for nuclear staining. Following a 30-minute incubation at room temperature, tumor cells were washed, resuspended in fresh culture medium and transferred to the IncuCyte chamber. After tumor cell adherence and spreading, CAR T cells were introduced at an effector-to-target (E:T) ratio of 1:5 in a final volume of 400 μL per well. CAR T cells were cultured in RPMI + 10% FBS + 1% P/S + 1% GlutaMAX. Live-cell imaging was conducted every 2 hours over a 96-hour period using a 20× objective, capturing phase-contrast, green and red fluorescence images to monitor tumor cell viability. The number of viable tumor cells was determined based on nuclear staining intensity and cytotoxicity was quantified as the percentage of tumor cell loss over time relative to untreated controls. For experiment reproducibility, biological duplicates and technical triplicates were performed per condition. To account for donor variability, CAR T cells from at least two independent donors were tested, with repeated experiments performed on a separate day using the same donors.

### 3D Bioprinted Tumor Infiltration Model

To assess CAR T cell infiltration into 3D neuroblastoma tumors, bioprinted tumor models were generated as previously reported^36^. Bioprinting was performed by Cellbricks GmBH (Berlin, Germany) using a hydrogel-based bioink optimized for neuroblastoma culture, allowing the creation of standardized cylindrical tumor constructs with precise size and volume. Each cylindrical tumor model had a diameter of 3 mm, a height of 1 mm and a total volume of 7.07 mm³, enabling controlled cell distribution and tumor architecture. SK-N-AS and SK-N-BE2c neuroblastoma cells, either unmodified or cytokine-expressing, were embedded within these constructs to model solid tumors. L1CAM-targeting CAR T cells were then co-cultured with the 3D tumors for 12 hours and infiltration efficiency was quantified by flow cytometry, using the T cell-to-tumor cell ratio as a readout. To further model physiological barriers to T cell migration, a trans-endothelial migration and tumor infiltration assay was developed by incorporating a HUVEC monolayer within a Boyden transwell insert, positioned atop the bioprinted tumor constructs. This system simulated vascular endothelium, requiring CAR T cells to migrate through an endothelial barrier before reaching the tumor mass. Vybrant™ DiO Cell-Labeling Solution (Thermo Fisher, Cat# V22886) was used to confirm HUVEC layer formation before T cell addition. After four hours of CAR T cell migration, the insert was removed and CAR T cell infiltration into the tumor mass was assessed 8 hours later by flow cytometry.

### Housing and Handling of Animals

All mouse experiments were approved by the regulatory agency (Landesamt für Gesundheit und Soziales Berlin, approval number: Anz.Ther.: Reg E0023-23) and were carried out in compliance with the German Law of Animal Rights. Mice were kept in a 12 h light/dark cycle, at a housing temperature of 23°C. Furthermore, food and water were available ad libitum. The animal welfare was checked twice daily. Body weights, tumor volume and general health conditions were recorded throughout the whole study.

### Xenograft Mouse Model, in Vivo CAR T cell Transplantation and Bioluminescence Imaging

For transplantation 5x10^6^ tumor cells were mixed with Matrigel (1:1) and transplanted subcutaneously in a final volume of 100 µl into 6–8 weeks old female CIEA NOG mouse® (nomenclature: NOD.Cg-Prkdcscid Il2rgtm1Sug/JicTac; genotype: sp/sp;ko/ko; Taconic Biosciences, Inc.). After engraftment (palpable tumor), tumor size was measured at least twice a week with a digital caliper. Tumor volume was calculated with the formula V = (length × width 2)/2. CAR T cell transplantation was performed at a tumor volume of at least 50 mm³. For in vivo 10 mio. CAR T cells were injected into the tail vein (i.v.) in 200 µl PBS by slow injection. For BLI mice were anesthetized with Isoflurane (Baxter, San Juan, Puerto Rico) and received intraperitoneally 150 mg/kg D-Luciferin (Biosynth, Staad, Switzerland) dissolved in PBS. BLI was performed with the NightOwl II LB983 in vivo imaging system. The IndiGO 2.0.5.0 software is used for initial analysis, color-coding of the signal intensity and quantification. BLI was performed d1, d4, d7, d11 and d14-26 for SK-N-AS and d1, d4, d7, d11, d17, d24 and d28 for SK-N-BE2c transplanted animals. Mice were sacrificed after reaching a tumor volume > 1,5cm^3^ or other ethical endpoints. Blood samples were collected after retrobulbar venous plexus puncture in MiniCollect® tubes containing Lithium Heparin for serum samples and processed concerning manufacturer’s instruction. Serum samples were stored at -80°C. For tumor tissue collection, mice were sacrificed. Tumors were removed, their weights determined and subsequently divided into two pieces, one of which was formalin-fixed and paraffin-embedded and one as a snap frozen sample.

### Statistical Analysis

All statistical analyses were performed using GraphPad Prism (version 10.2.0). Data are presented as mean ± standard deviation (SD) unless stated otherwise. For comparisons of more than two groups, Kruskal-Wallis tests with Dunn’s post hoc correction were applied. Pairwise comparisons were conducted using Mann-Whitney tests. Two-way analysis of variance (ANOVA) with Tukey’s post hoc test was used for multiple comparisons across conditions and time points. Linear regression was used for correlation curve fitting and Spearman test for correlation analyses. Logistic regression was used for growth curve fitting and growth coefficients k were used for proliferation rate comparison (Y=YM*Y0/((YM-Y0)*exp(-k*x) +Y0). For survival analysis in in vivo studies, Kaplan-Meier curves were generated. Bioluminescence to tumor volume ratios and total CAR T cell infiltration over time were analyzed using area under the curve (AUC) calculations followed by Mann-Whitney tests. For dPCR-based quantifications, two-way ANOVA was applied. Statistical significance was defined as *p* < 0.05 (*p* values: *<0.05, **<0.01, ***<0.001, ****<0.0001; n.s., not significant). Exact *p* values of key analyses from in vivo experiments are given in **Figure 6**, all other *p* values are available in the source data file. Sample sizes for each experiment are indicated in the respective figures and figure legends, as well as in the source data file.

Additional methods are described in the supplementary methods section. Equipment, consumables, antibodies and software used are listed in **Supplementary tables 5-7**.

## Author contributions

ML conceived and designed the study. ML, JH and MJ created the CancerPAM pipeline. ML, JM, SA, SB, MP, EO, AY, CV, CL, LA, MLA, JK and MS performed the experiments. ML, JM, SA, JH, SB, MP, EO and CV analyzed the data. ML, MJ, CV, LA, FZ, JK, MS, KA, DLW, RK and AK developed the methodology. ML wrote the manuscript with input from all authors. RK, AE and AK supervised the study.

## Data Sharing Statement

All data collected and analyzed in this study, including detailed test statistics, are available in the supplementary source data file.

Raw sequencing data will be available through the German Human Genome-Phenome Archive (NFDI-GHGA). The data will be made public upon publication.

The CancerPAM bioinformatics pipeline used in this study is available in a private GitHub repository for peer review at https://github.com/MichaelLaunspach/CancerPAM using the following credentials:

Username: CancerPAM-Reviewer

Password: #CancerPAM25

The repository will be made publicly accessible upon publication.

## Acknowledgments

This work was supported by the Berliner Krebsgesellschaft (grant number LAFF202008 to M.L.) and KINDerLEBEN e.V. Berlin (to M.L.). Additional support was provided by Charité - Universitätsmedizin Berlin and the Berlin Institute of Health (BIH) at Charité – Universitätsmedizin Berlin (AdHoc Booster Grant to D.W. and M.L.). M.L. participates in the BIH Charité Clinician Scientist Program, funded by Charité - Universitätsmedizin Berlin and the BIH. A.K. participates in the BIH Charité Advanced Clinician Scientist Pilot Program, also funded by Charité - Universitätsmedizin Berlin and the BIH.

The authors thank Silke Schwiebert and Anika Winkler for technical support with the experiments; Michael C. Jensen for providing the L1CAM-CAR T cell construct; Nikolaus Rajewsky for his support as a scientific mentor within the framework of the BIH Charité Clinician Scientist Program; and Kathy Astrahantseff for manuscript proofreading and editorial advice. Parts of Figures 1, 5, 6 and Supplementary Figure S16 were created with BioRender.com.

## Conflict of interest disclosure

ML has received reagents and services related to CRISPR–Cas gene editing and digital PCR from Qiagen. DLW is named as an inventor on patent applications related to genome editing and cell therapies. DLW is a co-founder and holds equity in TCBalance Biopharmaceuticals GmbH. DLW’s laboratory at Charité has received reagents and services related to CRISPR– Cas gene editing from Integrated DNA Technologies and GenScript Inc.

None of the aforementioned companies were involved in the design, execution, or interpretation of the study. All other authors declare no competing interests.

