## Supplement for "Personalized CRISPR Knock-In Cytokine Gene Therapy to Remodel the Tumor Microenvironment and Enhance CAR T Cell Therapy in Solid Tumors"

### CancerPAM: A Multi-Omics Pipeline for Personalized CRISPR Knock-In Cytokine Gene Therapy to Remodel the Tumor Microenvironment and Enhance CAR T Cell Therapy in Solid Tumors

- Supplement -

Supplementary Fig. S10. In vitro CRISPR knock-in. 19
Supplementary Fig. S11. Site-specific knock-in confirmation via Sanger sequencing and PAM frequency analysis. 20
Supplementary Fig. S12. Site-specific knock-in analysis via dPCR and IFNG toxicity testing. 21
Supplementary Fig. S13. Unspecific T cell knock-in. 22
Supplementary Fig. S14. dPCR analysis after FACS enrichment. 23
Supplementary Fig. S15. Characterization of enriched transgenic tumor cell lines. 24
Supplementary Fig. S16. CAR T cell activation and exhaustion in tumor cell line–CAR T cell co-culture experiments. 25
Supplementary Fig. S17. Cytokine secretion in tumor cell line–CAR T cell co-culture experiments. 26
Supplementary Fig. S18. Cytokine-mediated T cell migration in vitro. 27
Supplementary Fig. S19. 3D tumor infiltration assay. 28
Supplementary Fig. S20. In vivo CAR T cell infiltration model development. 29
Supplementary Fig. S21. In vivo CAR T cell infiltration experiment: pre-transplant and pre-treatment analysis. 30
Supplementary Fig. S22. Bioluminescence imaging of CAR T cell treated SK-N-AS xenograft neuroblasoma models. 31
Supplementary Fig. S23. Bioluminescence imaging of CAR T cell treated SK-N-BE2c xenograft neuroblasoma models. 32
Supplementary Fig. S24. In vivo CAR T cell infiltration in CXCL10-secreting xenograft tumors. 33
Supplementary Fig. S25. Immunofluorescence staining of tumor sections. 34
Supplementary Tables 35
Supplementary Table 1. Exemplary patient data for top and bottom rank novel PAM sites identified via CancerPAM 35
Supplementary Table 2. Neuroblastoma cell line target loci 35
Supplementary Table 3. HDRT sequences 36
Supplementary Table 4. Primer and probe sequences 37
Supplementary Table 5. Equipment and Consumables used 38
Supplementary Table 6. Antibodies used for flow cytometry and IF Staining 42
Supplementary Table 7. Software used 43

#### List of Abbreviations

AAV – Adeno-associated virus
AUC – Area under the curve
BLI – Bioluminescence Imaging
CAR – Chimeric antigen receptor
CFD – Cutting frequency determination
cDNA – Complementary DNA
CI – Confidence Interval
CMV – Cytomegalovirus (used as a promoter in gene expression)
CRISPR – Clustered regularly interspaced short palindromic repeats
DepMap – Cancer Dependency Map
dPCR – Digital polymerase chain reaction
dsDNA – Double-stranded DNA
EF1a – Elongation factor 1 alpha (used as a promoter in gene expression)
ELISA – Enzyme-linked immunosorbent assay
EGFRt – Truncated epidermal growth factor receptor
FACS – Fluorescence-activated cell sorting
FBS – Fetal bovine serum
FMO – Fluorescence minus one
GFP – Green fluorescent protein
gRNA – Guide RNA
HDR – Homology-directed repair
HDRT – Homology-directed repair template
HUVEC – Human Umbilical Vein Endothelial Cells (used in the trans-endothelial migration assay)
IFNG – Interferon-gamma
IL-2 – Interleukin-2
IL-7 – Interleukin-7
IL-15 – Interleukin-15
L1CAM – L1 cell adhesion molecule
LNP – Lipid nanoparticle
LTR – Long terminal repeat
MACS – Magnetic-activated cell sorting
MFI – Mean fluorescence intensity
MIT – Massachusetts Institute of Technology (used in CRISPR specificity scoring)
NGG – Most common protospacer adjacent motif (PAM) recognized by SpCas9
NK – Natural killer (cells)
NSG – NOD-scid IL2Rγnull (immunodeficient mouse model)
PAM – Protospacer adjacent motif
PBS – Phosphate-buffered saline
PBMCs – Peripheral blood mononuclear cells
PCR – Polymerase chain reaction
PE – Phycoerythrin
Pen/Strep - Penicillin/Streptomycin
qPCR – Quantitative polymerase chain reaction
Q8 – Custom reporter epitope tag (CD34 epitope and CD8 transmembrane domain)
REP – Rapid expansion protocol
RNP – Ribonucleoprotein
RPMI – Roswell Park Memorial Institute (medium)
SD – Standard deviation
sgRNA – Single-guide RNA
SNV – Single nucleotide variant
SORT – Selective organ targeting
sPA – Synthetic poly(A) sequence
TAE – Tris-acetate-EDTA (buffer for electrophoresis)
TME – Tumor microenvironment
TRAC – T cell receptor alpha constant
UV – Ultraviolet
VEGF – Vascular Endothelial Growth Factor
WES – Whole-exome sequencing
WGS – Whole-genome sequencing

#### Supplementary Notes

##### Supplementary Note 1

The increased frequency of novel PAM sites in neuroblastoma cell lines was associated with a higher prevalence of mutations toward cytosine (C) or guanine (G), as observed in these datasets **(Supplementary Fig. S2c).**

##### Supplementary Note 2

The FCGBP gene harbored five or more novel PAMs across all terminate NB cell lines, yet it did not exhibit a similar prominence in patient samples. Notably, there was no overlap between the top 10 genes with the highest novel PAM counts in cell lines and those in patient-derived data **(Fig. S1E)**. In patient exome data, DLG2 (Chr. 11) emerged as the gene with the highest prevalence of at least one novel PAM site, identified in 13 out of 54 patients **(Supplementary Fig. S1f)**. Across all patient samples, 17 novel PAM sites were recurrently identified in two individuals.

##### Supplementary Note 3

To further investigate discrepancies between cell lines and patient data, we applied CancerPAM to publicly available whole-exome sequencing (WES) datasets from 48 neuroblastoma cell lines (including eight from our dataset) available on the DepMap portal. In DepMap 23Q4, a median of 25 novel PAMs were identified from 155 SNVs (19% [95% CI: 17–22%]), a value more consistent with patient-derived data compared to terminate NB cell line data **(Supplementary Fig. S2a)**. However, cross-comparison of different DepMap versions (20Q4, 22Q2 and 23Q4) for the same 48 cell lines revealed inconsistencies. DepMap 23Q4 reported fewer SNVs and novel PAMs, with only partial overlap between versions **(Supplementary Fig. S2b,c)**. Additionally, DepMap 23Q4 exhibited a significantly lower percentage of SNVs relative to the terminate NB dataset (Supplementary Fig. S2d). Surprisingly, the concordance of SNVs between DepMap datasets and terminate NB was low, with only 46% of SNVs from DepMap 23Q4 and 45% from DepMap 20Q4/22Q2 overlapping with terminate NB, despite terminate NB reporting nearly 10-fold more SNVs than DepMap 23Q4 (median 1,470 vs. 155 SNVs) **(Supplementary Fig. S2e,g, S3)**. As expected, a gene-level analysis of chromosomes 1 and 19 revealed a higher proportion of genes with at least two novel PAM sites in terminate NB, reinforcing the increased mutation burden in these datasets. Only three genes on Chr. 1 and none on Chr. 19 harbored at least two novel PAMs shared across both datasets **(Supplementary Fig. S4).**

##### Supplementary Note 4

When analyzing the correlation of other features besides CRISPR specificity scores with the total PAM count per patient, a similar, albeit weaker, effect was observed for CRISPR efficiency scores, while no significant correlation was detected for copy number or gene expression. However, the copy number of top-ranked PAM sites was positively correlated with the total number of PAMs detected per patient, suggesting a potential structural genomic influence on PAM site availability **(Supplementary Fig. S7).**

##### Supplementary Note 5

To explore possible confounding factors, we examined the correlation between PAM feature characteristics and MYCN amplification status. Analysis of DepMap 23Q4 data indicated a slightly higher proportion of novel PAM sites located in non-essential genes in MYCN non-amplified cell lines. However, the observed effect size was marginal (median difference of 1.1%), and the findings could not be reproduced in the eight terminate NB cell lines with available dependency data, indicating that MYCN amplification status is unlikely to be a major determinant of PAM site distribution **(Supplementary Fig. S8).**

##### Supplementary Note 6

Preliminary CRISPR cutting efficiency experiments demonstrated relevant cutting activity (>5%) at all loci, except for *SCAF11*, *SH3BP1* and *SNX18*. The measured relative cutting frequency exhibited only a weak positive correlation with the Doench score and showed no correlation with the Moreno score. However, the strongest correlation - albeit not statistically significant - was observed for higher copy number regions **(Supplementary Fig. S9a, b).**

Supplementary Note 7
Optimization of HDRT constructs for knock-in revealed that a custom-designed EF1a-derived shortened promoter in combination with a Q8 (CD8 transmembrane + CD34 Qbend epitope) reporter exhibited the highest knock-in efficiency. In contrast, HDRTs incorporating a larger hCMV promoter or fluorophores such as RFP and GFP showed reduced efficiency, likely due to their increased genetic payload. We also tested a multicistronic vector encoding two cytokines, which demonstrated stable knock-in at the AAVS1 locus but with significantly lower efficiency.

Supplementary Note 8
We also evaluated a combinatorial knock-in strategy using two HDRTs (differing homology arms) and two gRNAs (targeting distinct tumor-specific loci) simultaneously, without increasing total DNA or gRNA dosage. This approach led to increased reporter expression after 28 days **(Supplementary Fig. S10b).** However, despite its efficacy, this strategy was not pursued in subsequent experiments due to the potential risk of chromosomal translocation. Lastly, we observed significantly lower knock-in rates for IFNG, particularly in SK-N-BE2c, suggesting locus- or transgene-specific effects on knock-in efficiency **(Fig. 3d, Supplementary Fig. S10c).**

Supplementary Note 9
Additional Sanger sequencing of Out/In PCR products provided definitive proof of on-target knock-ins **(Supplementary Fig. S11a).** Further analysis using Out/Out preferential amplification followed by Sanger sequencing and PAM site frequency determination confirmed the loss of the novel PAM site post-knock-in on a quantitative level **(Supplementary Fig. S11c)**. Analysis of weak-positive dPCR signals indicated reduced performance and sensitivity of the Out/In dPCR assay used for investigating AAVS1 knock-ins, which explains the discrepancy between higher knock-in rates observed in flow cytometry compared to dPCR for AAVS1 knock-ins **(Supplementary Fig. S12c).**

Supplementary Note 10
Unspecific knock-in, was assumedly not entirely due to CRISPR/Cas9-related effects, as transfection of high doses of Q8-HDRT alone also led to an increase in Q8+ positivity after 21 days - although this was not statistically significant- when compared to Q8+ positivity of control cells transfected with a GFP-HDRT. After subtracting the mean Q8+ count, only *IGSF9B* and *RPLP0* remained as gRNAs significantly associated with CRISPR-mediated unspecific knock-in (**Fig. 4e, Supplementary Fig. S13c**). The observed trend for inverse correlation between unspecific knock-in rates and CancerPAM ranking scores (**Fig.4g,h**) persisted after subtracting mean DNA-only transfection Q8+ cell counts **(Supplementary Fig. S13d**).

Supplementary Note 11
We did not observe any growth advantage or disadvantage in the enriched transgenic tumor cell lines, regardless of cell line, target locus, or cytokine combination **(Fig. 5C, Supplementary Fig. S14a,b).**

Supplementary Note 12
During in vitro CAR T cell SK-N-AS tumor co-culture, we did not observe any differences in the expression of activation markers (CD25, CD137) or exhaustion markers (LAG3, TIM3, PD1), as measured via flow cytometry, among the different cytokine-expressing tumor cell lines **(Supplementary Fig. S16b,c)**. Interestingly, co-culture with tumor cells resulted in elevated CXCL10 and CXCL11 levels in all CAR T cell plus unmodified or transgenic tumors, except when co-cultured with IFNG-expressing tumors. Regarding IFNG release, we observed that CAR T cell-mediated IFNG secretion was only a fraction of the levels produced by IFNG-transgenic tumors. CAR T cell IL-2 secretion remained unaffected by additional cytokine secretion from tumor cells **(Supplementary Fig. S17).**

Supplementary Note 13
After confirming similar engraftment across all treatment groups, comparison of day 1 BLI data post-CAR T cell injection in the lung showed no significant differences between tumor groups **(Supplementary Figs. S24a,b).**

#### Supplementary Note 14

Serum cytokine level analysis via ELISA confirmed CXCL10 expression in mice harboring transgene-positive tumors, both pre- and post-treatment **(Supplementary Fig. S21a, S24f).** Additionally, post-mortem immunofluorescence analysis of tumor sections confirmed persistent L1CAM positivity and low-abundance CD3+ T cells within tumors **(Supplementary Fig. S25).**

#### Supplementary Methods

##### Genomic DNA Sample Isolation

Genomic DNA was extracted from cultured cells using the QIAamp DNA Mini Kit (QIAGEN) following the manufacturer's guidelines with minor optimizations. Briefly, up to 5 × $10^6$ cells were lysed using Proteinase K and buffer AL at 56°C, followed by ethanol precipitation. The lysate was passed through a QIAamp Mini Spin Column via centrifugation to bind DNA, followed by sequential washes with AW1 and AW2 buffers to remove contaminants. Purified DNA was eluted in Buffer AE and stored at -20°C until further use.DNA concentration and purity were assessed using a NanoDrop 2000 spectrophotometer (Thermo Fisher Scientific) at 260 nm absorbance, ensuring concentrations ≥10 ng/µL for downstream applications.

##### Standard PCR

PCR was performed using KAPA polymerase (Roche) to amplify DNA sequences for homology-directed repair templates (HDRTs), target sequencing and qualitative knock-in analysis. Primers were custom-designed using SnapGene (GSL Biotech LLC) and Primer3Plus (https://www.primer3plus.com/) and Primer-BLAST (NCBI). PCR reactions were carried out using a C1000 Touch Thermal Cycler (Bio-Rad) under optimized cycling conditions. For sequence analysis, amplified fragments purified using the QIAquick PCR Purification Kit (QIAGEN) and sequenced by LGC Genomics. PCR product specificity and efficiency were confirmed by agarose (1-2%) gel-electrophoresis using TAE buffer-based gels, with GelRed (Sigma-Aldrich) staining for visualization. A detailed list of primers and oligonucleotides used in the study is provided in **Supplementary Table 4**.

##### Evaluation of Cell Proliferation and Viability Following CRISPR/Cas9-Mediated Gene Integration

Genome-edited SK-N-BE2c and SK-N-AS neuroblastoma cell lines, along with their wild-type counterparts, were used for proliferation and viability assessments. Cell proliferation was monitored over seven days using the IncuCyte® Live-Cell Analysis System (Sartorius). Cells were seeded in 96-well flat-bottom plates at 1 × $10^4$ cells per well in biological duplicates and technical triplicates. Phase-contrast imaging was performed at regular intervals to generate quantitative growth curves. Viability was assessed using IncuCyte® Cytotox Green reagent (Sartorius, catalog #4633) at a final concentration of 250 nM, added during seeding. Green fluorescence was monitored using IncuCyte® software to quantify non-viable cells with compromised membrane integrity.

##### Cytokine Quantification via ELISA

Cytokine levels were quantified using enzyme-linked immunosorbent assays (ELISA) following the manufacturer's protocols. CXCL10 and CXCL11 concentrations were determined using the Human CXCL10/IP-10 and CXCL11/I-TAC DuoSet® ELISA kits (R&D Systems, Cat# DY266 and DY392), while IL-2 and IFNG levels were measured using the BD OptEIA™ ELISA sets (BD Biosciences). 96-well microplates (NUNC) were coated overnight at 4°C with the respective cytokine-specific capture antibodies. After washing, plates were blocked with 10% fetal calf serum (FCS) for IL-2 and IFNG ELISAs or 1% bovine serum albumin (BSA) for CXCL10 and CXCL11 ELISAs. Frozen samples were thawed and analyzed at multiple dilution factors to ensure accurate quantification: 1:10, 1:100, or undiluted. Detection was performed using biotinylated secondary antibodies, streptavidin-HRP and TMB substrate, with signal development for 20 min at room temperature in the dark. The reaction was stopped with 2N $H_2SO_4$ (50 µL/well). Absorbance was measured at 450 nm (reference: 570 nm) using an Epoch™ microplate spectrophotometer (Gene5 software). Standard curves were generated using four-parameter linear regression in Microsoft Excel.

##### Flow Cytometry and Fluorescence-Activated Cell Sorting (FACS)

Flow cytometry was used to analyze transgene expression, cell surface markers and viability, while fluorescence-activated cell sorting (FACS) was performed to enrich successfully modified cells. All flow cytometric analyses were conducted on BD FAC LSRFortessa X-20 (Lasers for Detection: 405 nm (violet), 488 nm (blue), 561 nm (yellow-green), 640 nm (red)), with sorting performed using a BD FACSAria III. Data acquisition and analysis were performed using FlowJo v10. A comprehensive list of all antibodies, fluorochromes is provided in **Supplementary Table 6** and detailed gating strategies are illustrated in **Supplementary Figures S10, S19 and S21**. Cells were collected, washed twice with PBS and resuspended in 100 µL staining solution containing the respective fluorochrome-conjugated antibodies. For CD34-epitope detection and sorting, cells were labeled with PE-conjugated anti-CD34 monoclonal antibody (clone QBEND/10, Invitrogen, dilution 1:10) and incubated at 4°C for 30 min in the dark. For samples requiring CAR expression analysis, cells were pre-stained with anti-mouse F(ab')2 and blocked with mouse serum (MilliporeSigma) before antibody labeling. FACS was used to enrich CD34-positive genome-edited cells, ensuring a homogeneous population. Cells were harvested, stained for CD34 expression and sorted using BD FACSAria III. The sorted fraction was collected in FBS-coated tubes, resuspended in culture medium and expanded for further analyses. Fluorescence signals were compensated using single-stained CD8+ T cells.

#### Boyden chamber and Clearview trans well migration assays

The migration potential of L1CAM-specific CAR T cells was assessed using a 24-well Boyden chamber transwell system with 8 µm pore size polycarbonate membranes. Inserts were pre-coated with 0.5% bovine serum albumin (BSA) in PBS for 1 hour at room temperature. CAR T cells were resuspended at $5 \times 10^6$ cells/mL at day 12-15 after expansion initiation seeded into the upper transwell chamber (100 µL per well). The lower chamber contained 500 µL of conditioned medium from cytokine-expressing neuroblastoma cells or control medium. Migration was monitored for 4 hours using the IncuCyte S3 live-cell imaging system and data were analyzed using the IncuCyte S3 software. Data were normalized to initial seeding density and analyzed using GraphPad Prism with migration parameters compared via Gompertz-Laird model fitting and statistical tests. The ClearView Transwell Migration Assay was performed using the IncuCyte ClearView Cell Migration Plate in a 96-well format with a pore size of 8 µm. The membrane surface was coated with 1% BSA solution before being incubated with Protein G (20 µg/mL, Thermo Fisher, #101200) for 1 hour at 37°C, followed by an additional 40 µL wash with Protein G solution. The membrane was then coated with ICAM-1 (5 µg/mL, Sino Biological, #10346-H03H) for 2 hours at 37°C before both sides were blocked with PBS + 1% BSA for 30 min at room temperature. CAR T cells (50,000 per well) were seeded at day 12-15 after expansion initiation in 60 µL RPMI + 0.5% FCS and allowed to attach for 45-60 min before the addition of 200 µL chemoattractant solution prepared in RPMI + 0.5% FCS to the lower chamber.

#### Generation of L1CAM knock-out SK-N-BE2c

SK-N-BE2c cells were transfected with a plasmid encoding Cas9 and a L1CAM-targeting guide RNA and a puromycin resistance gene. Specifically, $1 \times 10^6$ cells were seeded in a 6-well plate containing 3 mL of culture medium. The following day, the medium was replaced and cells were transfected using Effectene Transfection Reagent (QIAGEN) according to the manufacturer's protocol, utilizing 1 µg of plasmid DNA per transfection. On day two post-transfection, the medium was refreshed and on day three, selection commenced with 0.5 µg/mL puromycin. A control plasmid lacking the puromycin resistance gene was used as a negative control. By day ten, knockout efficiency was assessed via flow cytometry. Subsequently, to create a cell line with complete L1CAM knockout, single-cell clones were isolated and expanded.

#### Immunohistochemistry (IHC)

Immunohistochemistry was performed on paraffin-embedded tumor tissues. Tumors were fixed, dehydrated and embedded in paraffin before sectioning into 6 µm slices using a microtome. Sections were mounted on positively charged glass slides and subjected to deparaffinization in xylene (twice, 5 min each), followed by rehydration in a graded ethanol series (100%, 95%, 70% and 50% ethanol, 5 min each) and rinsing in distilled water. Antigen retrieval was performed in citrate buffer (10 mM tri-sodium citrate, 0.05% Tween-20, pH 6.0, Abcam ab93678) using a pressure cooker (2 min at 125°C, cooling to 90°C, then under running water for 10 min). After an additional 5 min incubation in distilled water, sections were washed in PBS (5 min). To prevent non-specific binding, blocking was performed using 3% normal serum (corresponding to the species of the secondary antibody) in TBS-T for 1 hour in a humidity chamber. For immunofluorescence staining, primary antibodies were applied in 3% serum in TBS-T and incubated overnight at 4°C. The following primary antibodies were used: rabbit anti-human/mouse CD3 (Abcam, ab16669, clone SP7, 1:100), mouse anti-human L1CAM (Invitrogen, MA1-46044, clone UJ127.11, 1:500). The next day, slides were washed five times in TBS-

T before incubation with fluorescently labeled secondary antibodies in TBS-T and 3% serum for 1 hour at room temperature. The following secondary antibodies were used: donkey anti-rabbit Alexa 594 (Invitrogen, Lot# 1938375, 1:300), donkey anti-mouse Alexa 488 (Invitrogen, Lot# 1975519, 1:300), goat anti-rabbit Alexa 594 (Invitrogen, 1:500), goat anti-mouse Alexa 488 (Invitrogen, Lot# A11001, 1:500). Following secondary antibody incubation, slides were washed 3 times in TBS-T and counterstained with Hoechst (1:2000 in TBS-T, 2–5 min). After a final wash in distilled water (5 min), samples were mounted using Eukitt quick-hardening mounting medium (Sigma-Aldrich, Lot# BCBP7505V). Immunofluorescence images were acquired using a fluorescence microscope (Olympus BX43) and analyzed with Olympus cellSens software.

#### Supplementary Figures

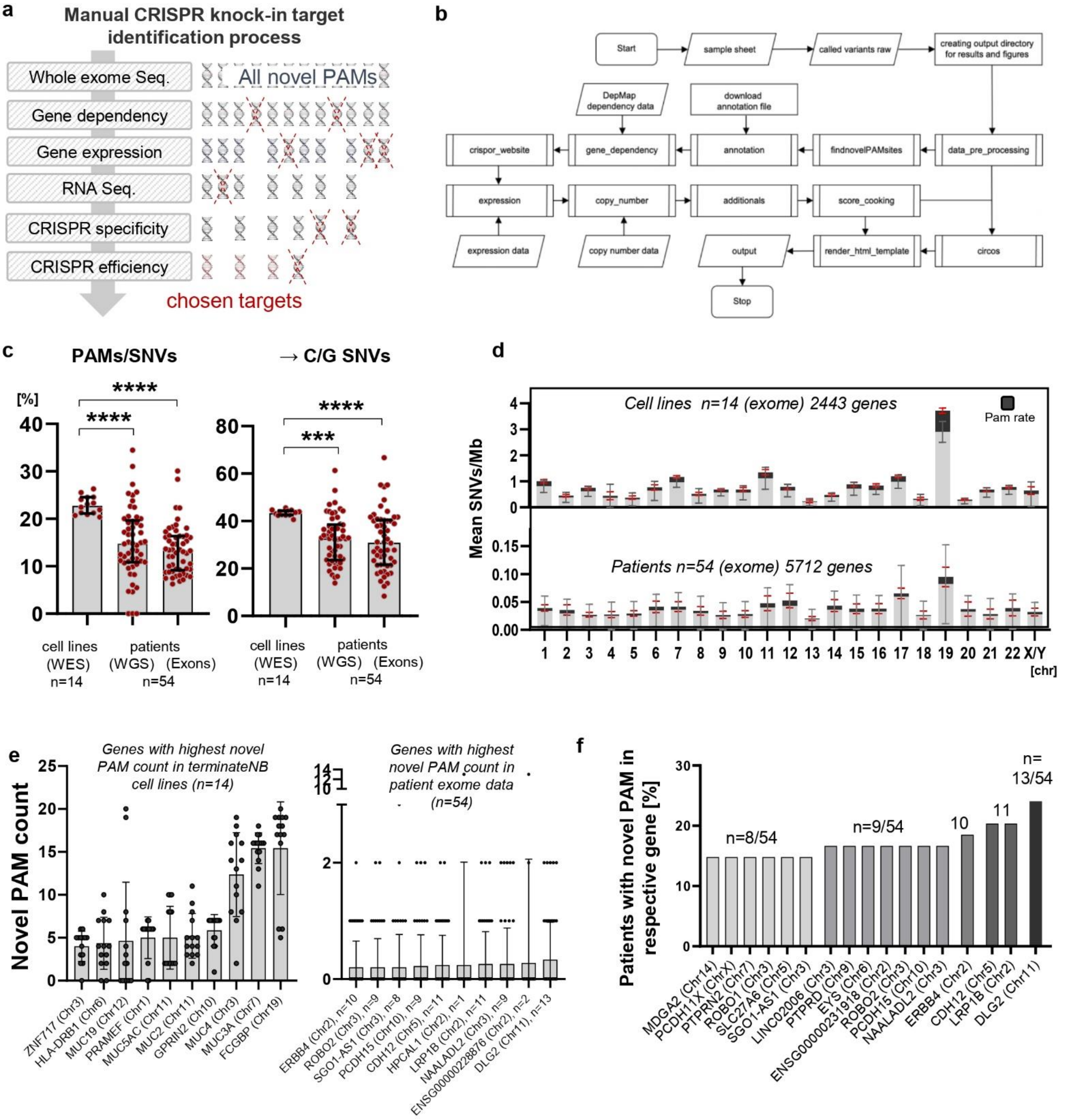

**Supplementary Fig. S1. CancerPAM multiomics-based automated pipeline. (a)** Manual multi-step CRISPR knock-in target identification process preceding the CancerPAM pipeline, where novel PAM sites were excluded as potential targets if they fell below defined threshold values at each step. **(b)** Visualization of the CancerPAM pipeline data flow. **(c)** Comparison of the proportion of novel PAMs within all SNVs and mutations to C or G (→ C/G) in cell line versus patient data. **(d)** Chromosomal distribution of exonic mutations in neuroblastoma cell lines and patient samples(F-H) Proportion of patients with novel PAM counts in the same gene, focusing on the 16 genes most frequently recurring across patients. **(e)** Quantification of novel PAM counts in the top 10 genes with the highest novel PAM counts in cell line or patient data. **(f)** Proportion of patients with novel PAM counts in the same gene, focusing on the 16 genes most frequently recurring across patients. *Data presentation: (c,d) Means ± SD. Statistical analysis: (c) Kruskal-Wallis with Dunn's post hoc test. p values: *<0.05, **<0.01, ***<0.001, ****<0.0001; n.s.: not significant.*

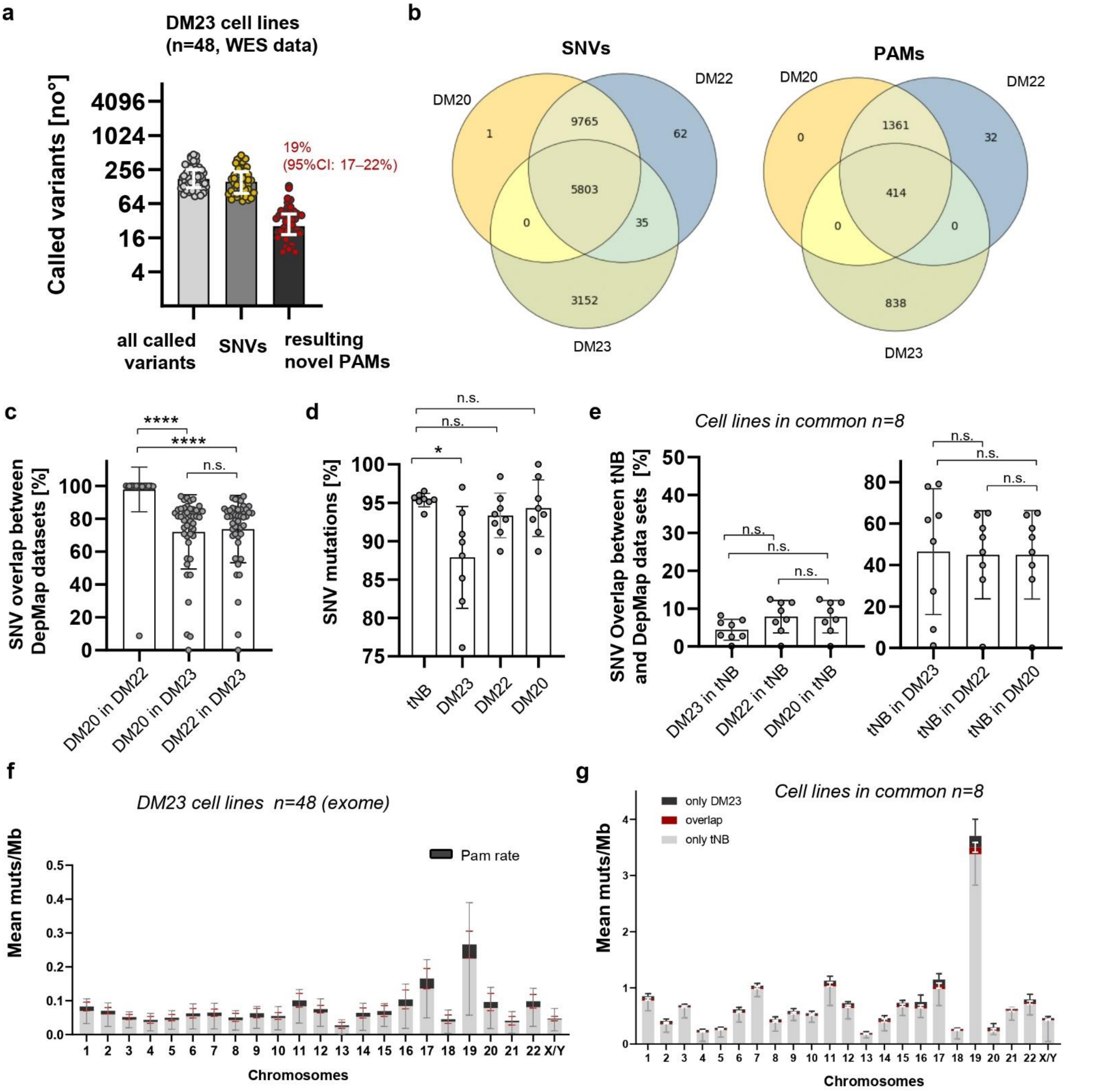

**Supplementary Fig. S2. Dataset comparison between Terminate NB and DepMap. (a)** Quantification of identified variants and tumor-specific PAM sites in neuroblastoma cell lines from the DepMap 23Q4 (DM23) dataset (WES data) identified using the CancerPAM pipeline. **(b)** Overlap of SNVs and novel PAMs identified in neuroblastoma cell lines across the three DepMap datasets: DepMap 20Q4 (DM20), DepMap 22Q2 (DM22) and DepMap 23Q4 (DM23). **(c)** Overlap of identified SNVs between DepMap datasets, where bars represent the percentage of SNVs in the first dataset that were also found in the second dataset. **(d)** Comparison of the percentage of identified SNVs between terminate NB (tNB) and DepMap datasets. **(e)** Overlap of identified SNVs between terminate NB and DepMap datasets for all cell lines present in all datasets, with bars showing the percentage of SNVs in the first dataset also identified in the second dataset. **(f)** Chromosomal distribution of exonic mutations in neuroblastoma cell lines from the DM23 dataset. **(g)** Overlap of identified SNVs and novel PAMs between terminate NB and the DM23 dataset per chromosome. *Data presentation: (a,d-e) Means ± SD. Statistical analysis: (c-e) Kruskal-Wallis with Dunn's post hoc test; p values: *<0.05, **<0.01, ***<0.001, ****<0.0001; n.s.: not significant.*

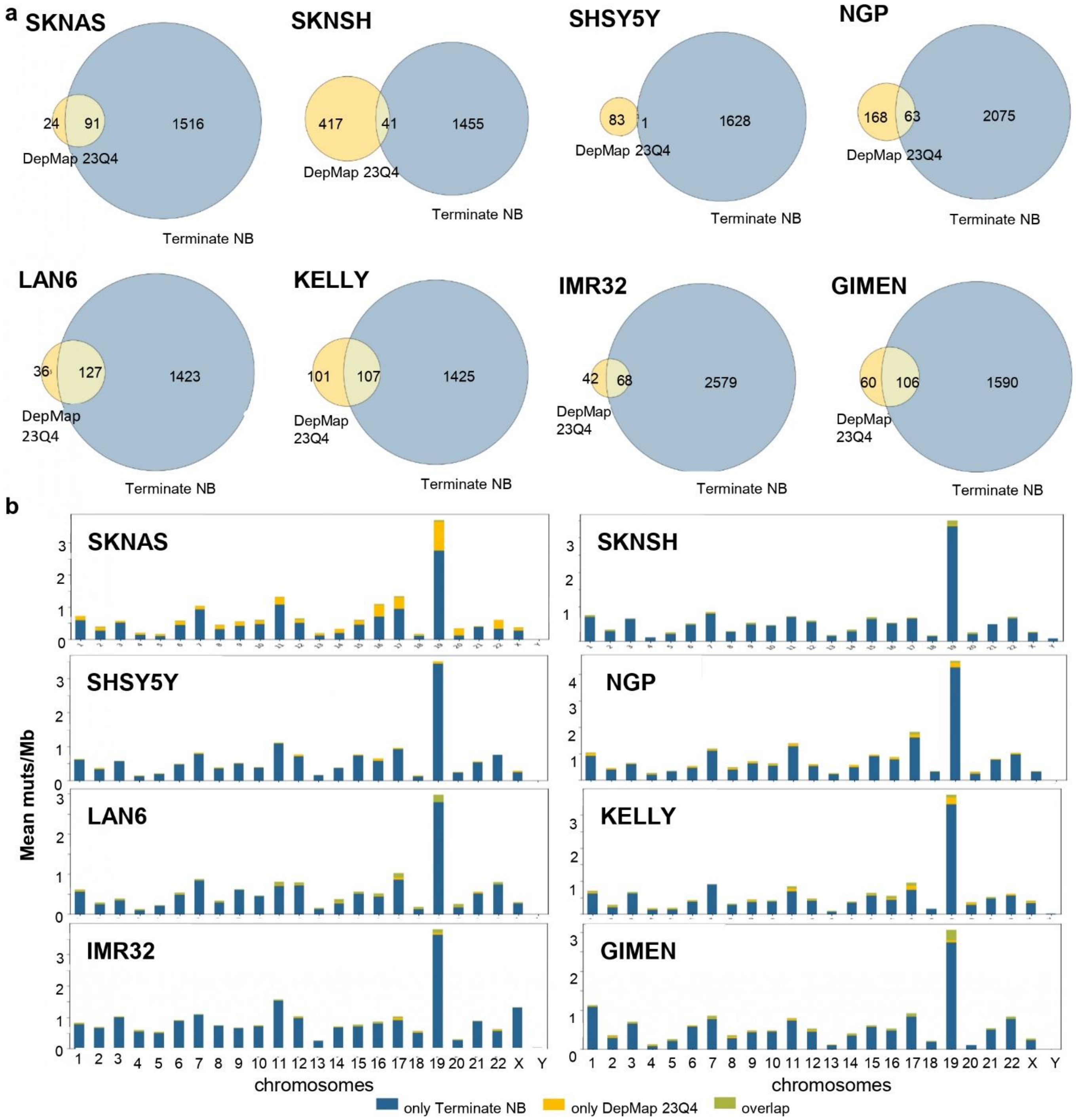

**Supplementary Fig. S3. Cell line data comparison between Terminate NB and DepMap data sets. (a)** Overlap of SNVs identified in neuroblastoma cell lines across the terminate NB and DepMap 23Q4 data sets. **(b)** Overlap of identified SNVs and novel PAMs between terminate NB and the DepMap 23Q4 dataset per chromosome for all cell lines included in both data sets.

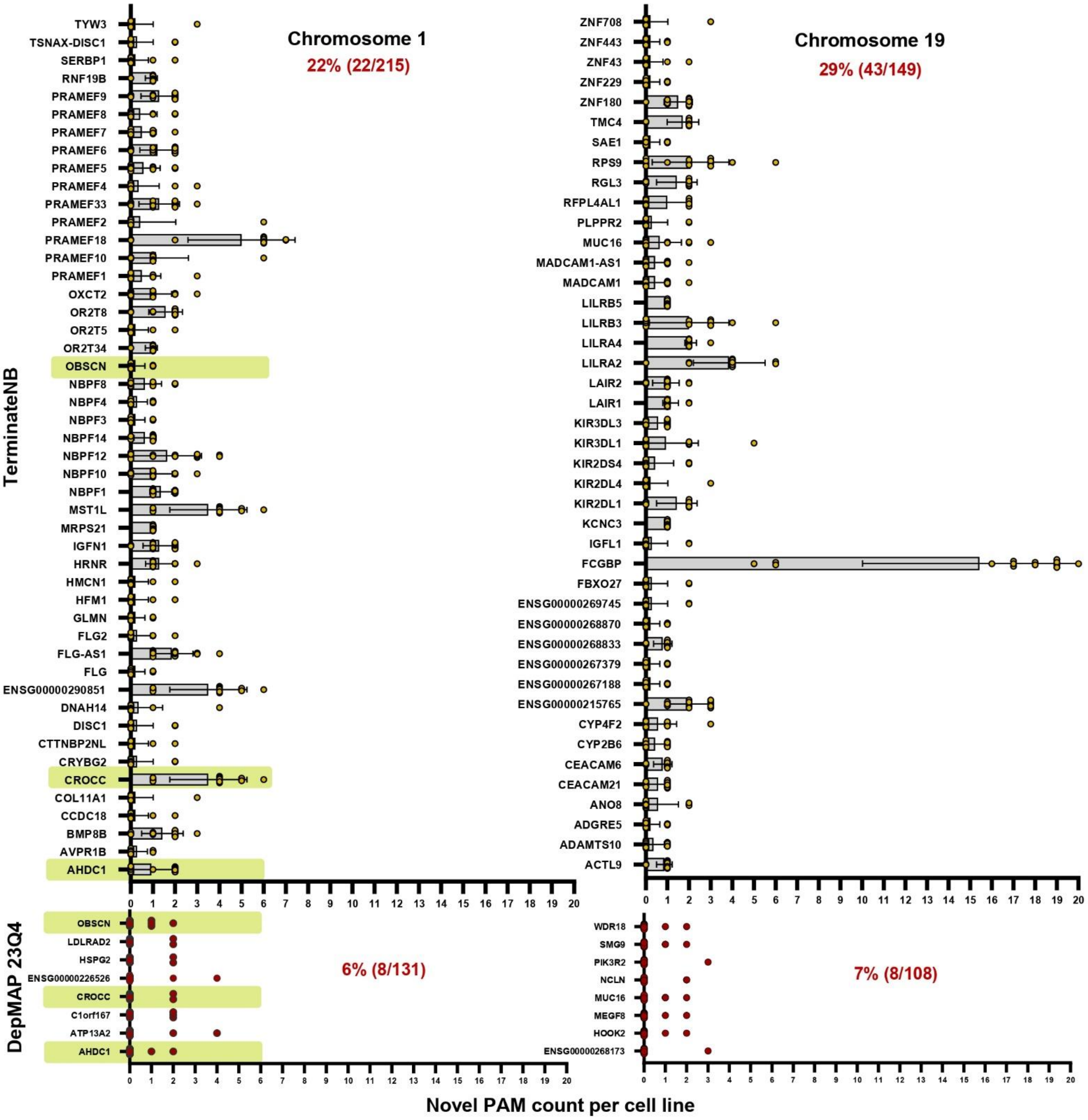

**Supplementary Fig. S4. Chromosomal novel PAM distribution.** Visualization of all genes with at least two novel PAMs on Chromosomes 1 and 19 found in at least one cell line from the terminate NB cell line dataset (n = 14) and the DepMap 23Q4 dataset (n = 48). In the terminate NB dataset, 22% (22/215) and 29% (43/149) of the genes on Chromosomes 1 and 19, respectively, with novel PAMs contained at least two novel PAMs in at least one analyzed cell line. For the DepMap 23Q4 dataset, this proportion was 6% (8/131) and 7% (8/108) for Chromosomes 1 and 19, respectively. Green bars indicate genes with at least two novel PAMs in at least one cell line that were shared between both datasets.

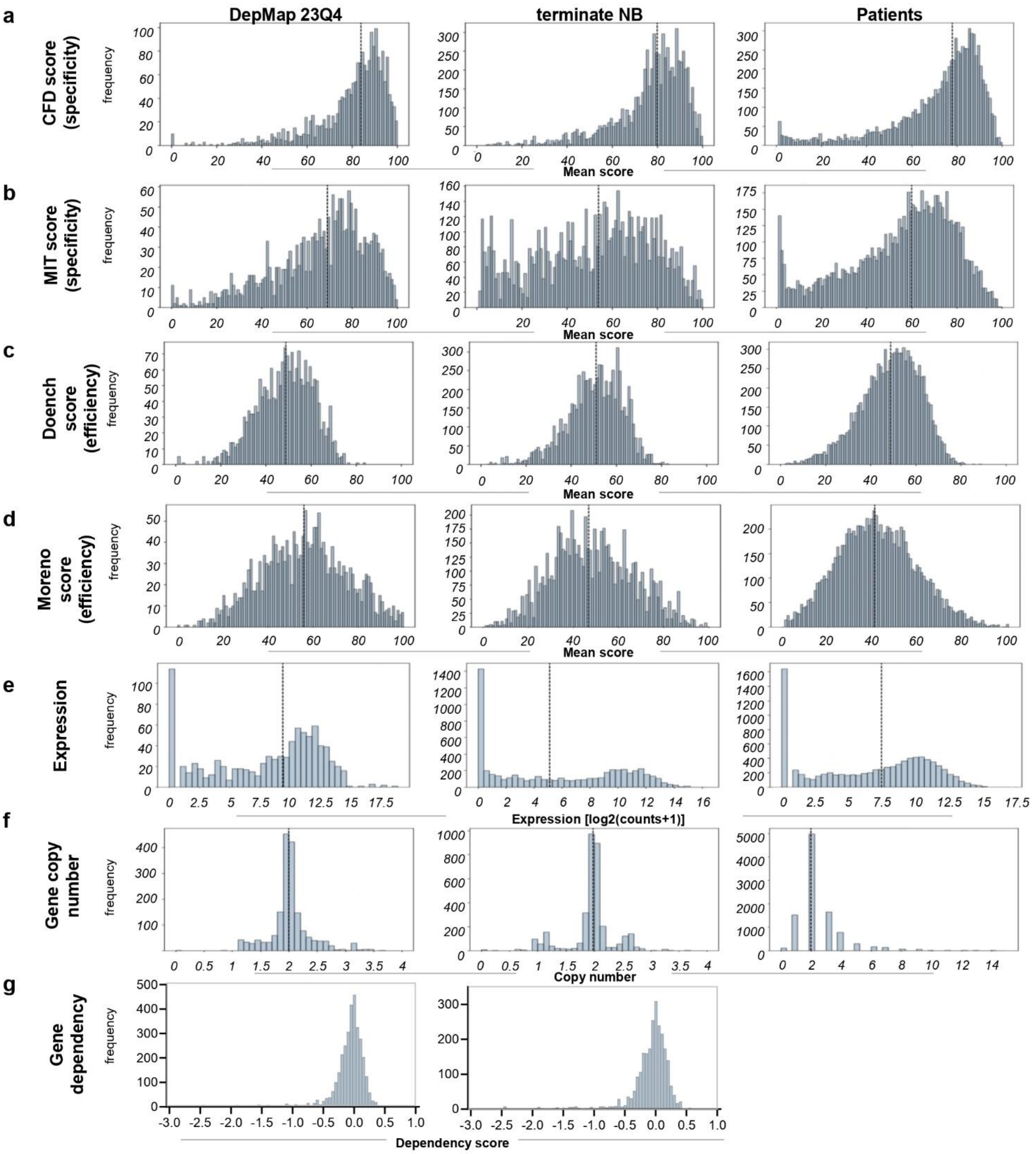

**Supplementary Fig. S5. Novel PAM feature characteristics distribution. (a-g)** Frequency distribution of values for various features annotated to the identified novel PAM sites in the DepMap 23Q4, terminate NB and patient datasets. The DepMap 23Q4 dataset includes data from 48 cell lines, terminate NB includes 14 cell lines and the patient dataset consists of 54 patients. Analyzed features include the CRISPR specificity scores CFD (**a**) and MIT (**b**), the CRISPR efficiency scores Doench (**c**) and Moreno (**d**), as well as gene expression levels **(e),** copy number **(f)** and Gene dependency scores **(g)** associated with novel PAM sites.

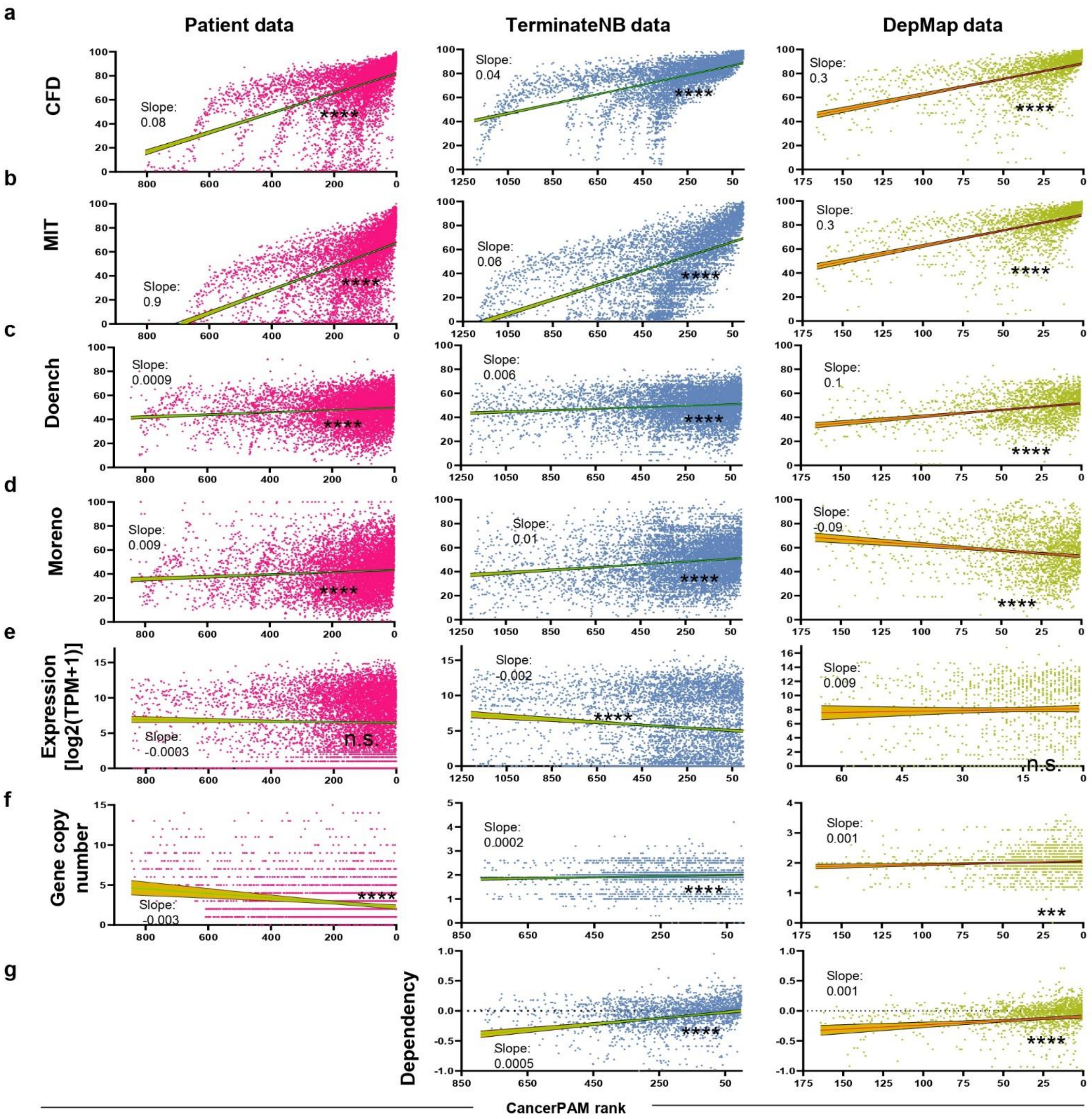

**Supplementary Fig. S6. CancerPAM ranking validation. (a-g)** Correlation analysis of novel PAM feature values against the CancerPAM rank of the identified novel PAM sites. Analyzed features include the CRISPR specificity scores CFD (**a**) and MIT (**b**), the CRISPR efficiency scores Doench (**c**) and Moreno (**d**), gene expression levels (**e**), copy number **(f)** and DepMap gene dependency (**g**, not available for patient data) from genes associated with the novel PAM sites. *Statistical analysis: Linear regression for curve fitting and slope calculation, followed by Spearman correlation analysis. p values: *<0.05, **<0.01, ***<0.001, ***<0.0001; n.s.: not significant.*

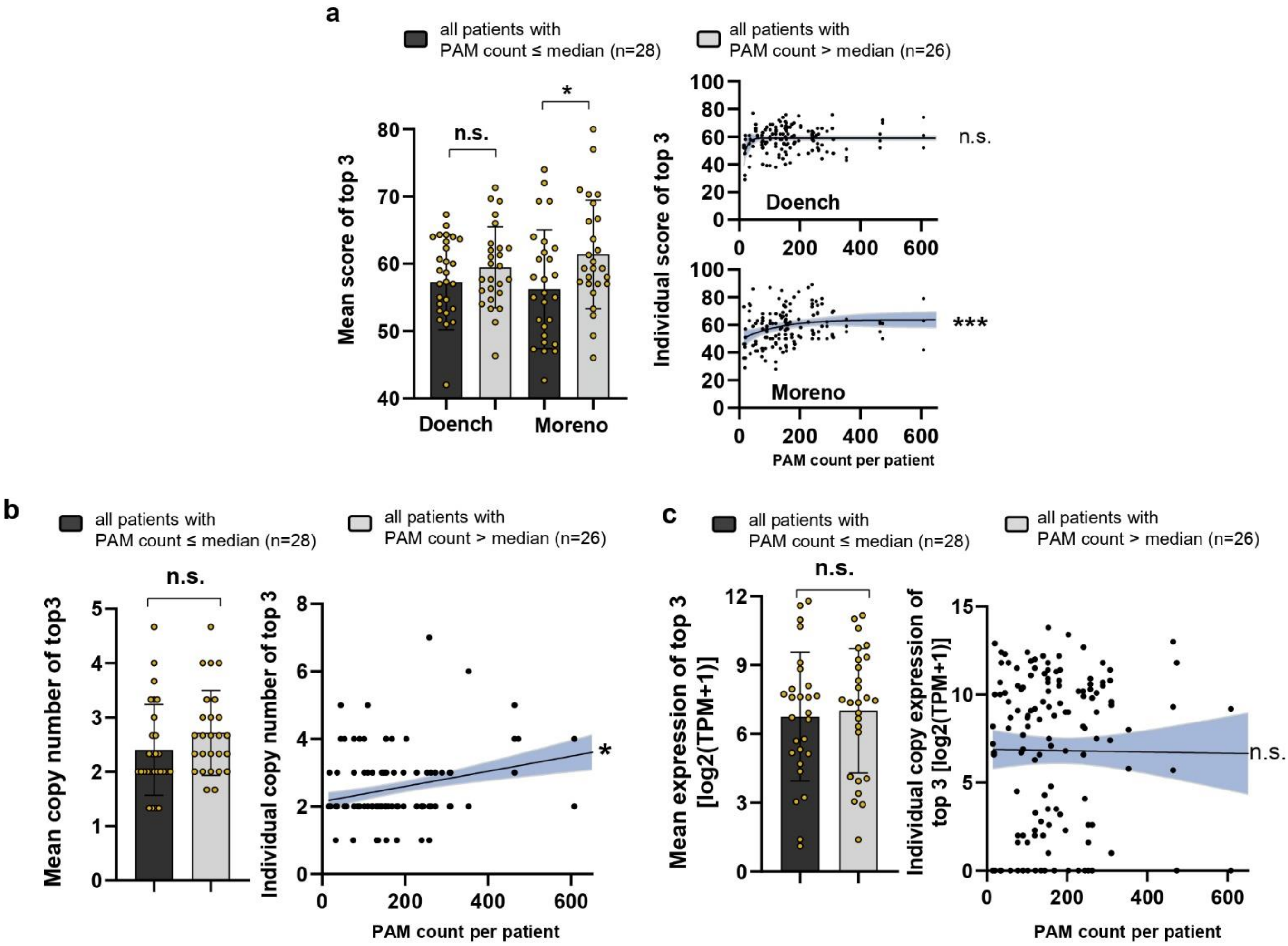

Supplementary Fig. S7. Feature correlation with PAM count. (a-c) Bar graphs comparing patients with a high (>median; n = 26) versus low (≤median; n = 28) PAM count in terms of the top three ranked PAM mean Doench and Moreno efficiency scores (a), copy number (b), or expression levels (c) X/Y scatter plots depict the correlation of individual scores for the top three ranked novel PAM sites across patients (n = 162) against total tumor-specific PAM counts per patient. *Data presentation: (a-c) Means ± SD. Statistical tests: (a-c) Bar graphs: Kruskal-Wallis with Dunn's post hoc test; X/Y scatter plots: logistic and linear regression for curve fitting and Spearman correlation. p values: *<0.05, **<0.01, ***<0.001, ****<0.0001; n.s.: not significant.*

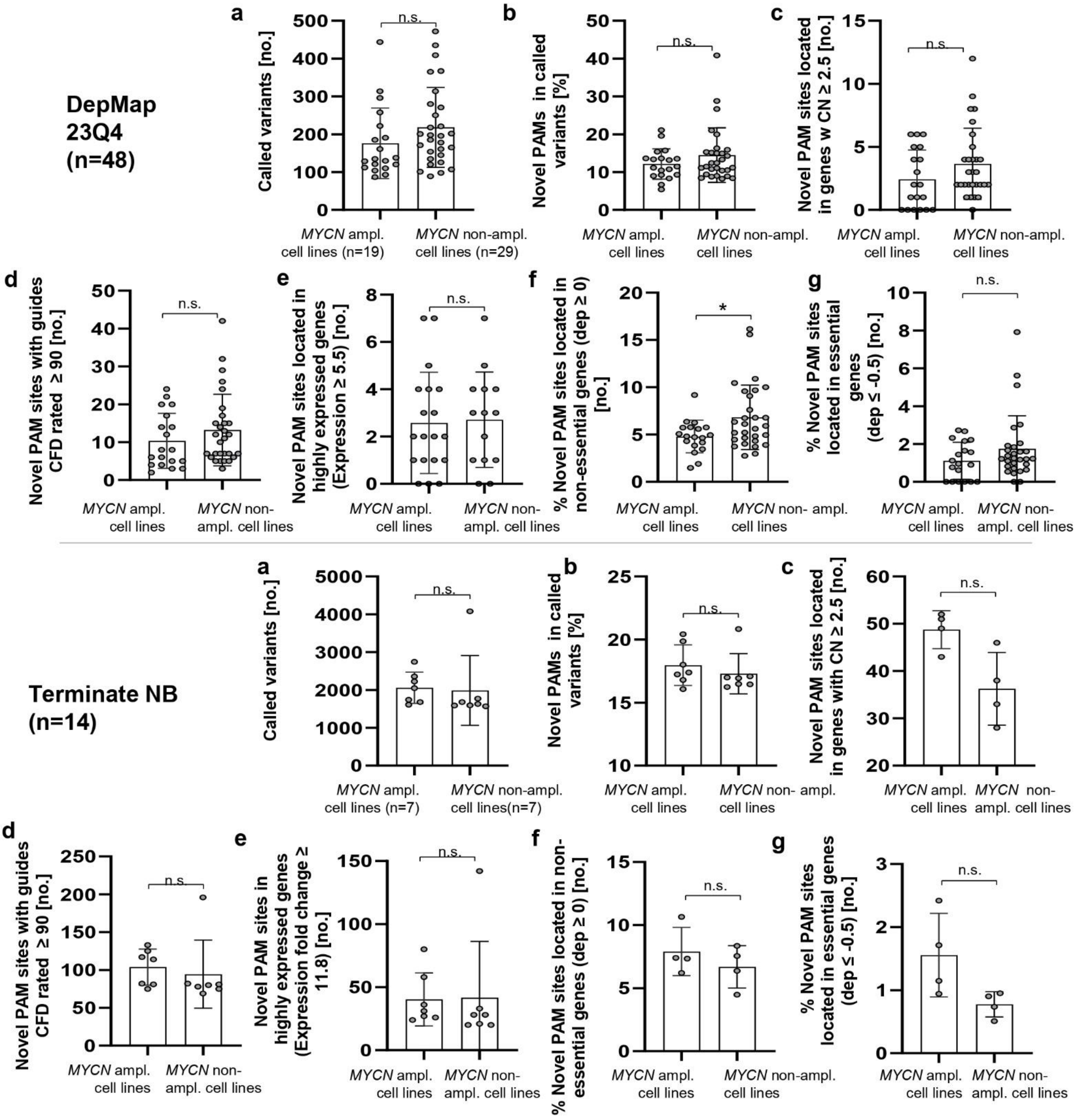

Supplementary Fig. S8. Influence of MYCN amplification status on novel PAM counts and features. (a-g) Group comparisons of novel PAM counts and feature values between MYCN-amplified and non-amplified cell lines for the terminate NB and DepMap 23Q4 datasets. Comparisons include the number of called variants per cell line **(a)**, the percentage of novel PAMs among all called variants **(b)**, the number of novel PAMs located in genes with a copy number (CN) ≥ 2.5 **(c)**, the number of novel PAM sites with guides showing a CFD score ≥ 90 **(d)**, the number of novel PAM sites located in highly expressed genes with log2(counts+1) ≥ 5.5 **(e)**, the number of novel PAM sites located in non-essential genes with gene dependency (dep) values ≥ 0 **(f)** and the number of novel PAM sites located in essential genes with gene dependency (dep) values ≤ -0.5 **(g)**. *Data presentation: (a-g) Means ± SD. Statistical analysis: (a-g): Mann-Whitney test. p values: *<0.05, **<0.01, ***<0.001, ***<0.0001; n.s.: not significant.*

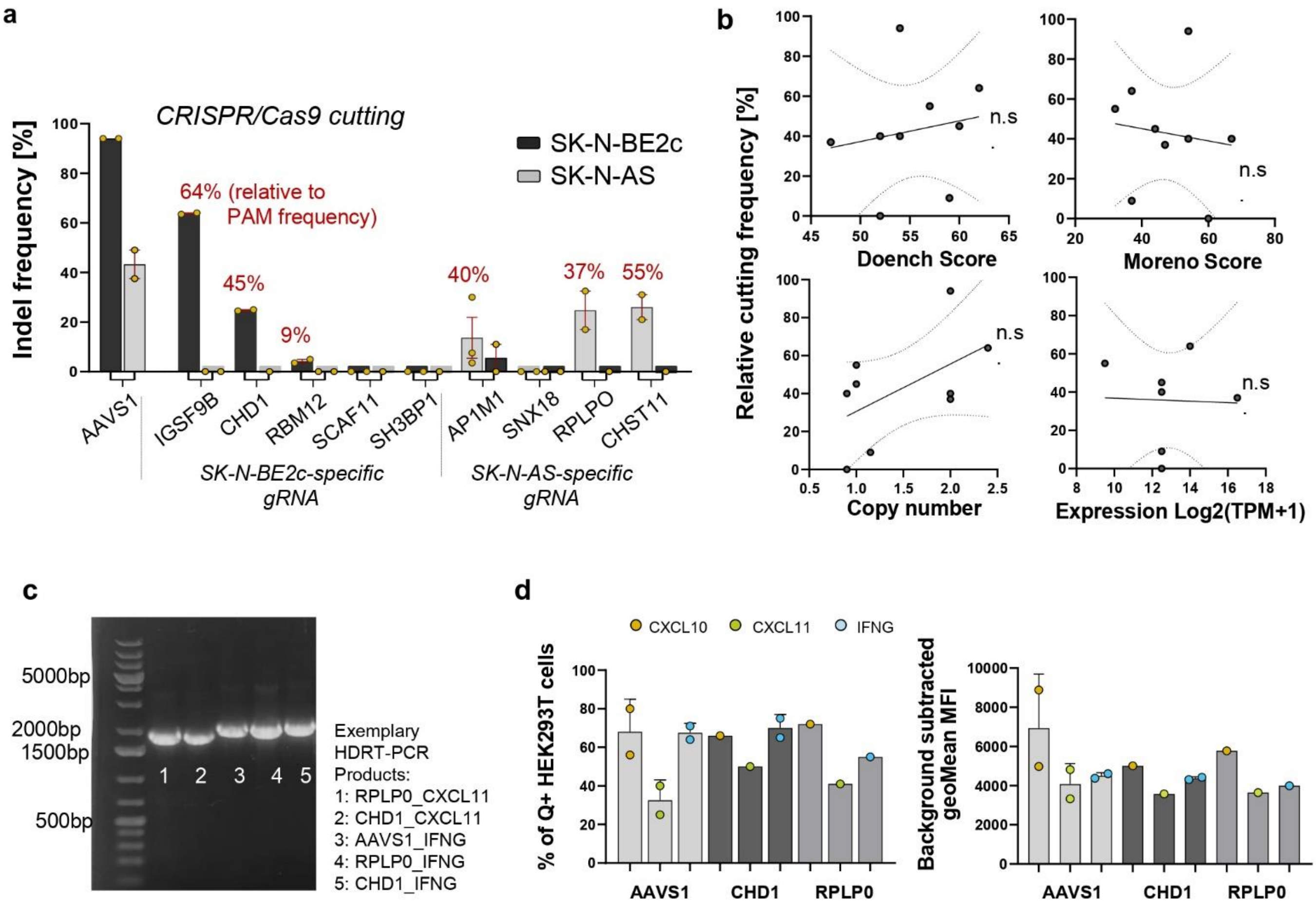

**Supplementary Fig. S9. In vitro CRISPR cutting and transgene vector production and testing. (a)** CRISPR/Cas9 cutting efficiency at the chosen target sites and the AAVS1 control locus, represented by indel frequency determined via Sanger sequencing and Synthego ICE analysis. **(b)** Correlation between relative cutting frequency (INDEL% / target site frequency) and the CRISPR efficiency scores (Doench and Moreno), as well as target gene copy number and expression. **(c)** Exemplary DNA gel electrophoresis result after homology directed repair template (HDRT) production using PCR and bead-based clean up. **(d)** Transgene expression from the produced HDRTs as represented by the percentage of positive cells and mean fluorescence intensity (MFI) in a preliminary vector expression test using HEK293T cells and the Effectene transfection reagent. *Data presentation: (a, d) Means ± SD. Statistical analysis: (e) Linear regression for curve fitting and Spearman correlation. n.s., not significant.*

**a**

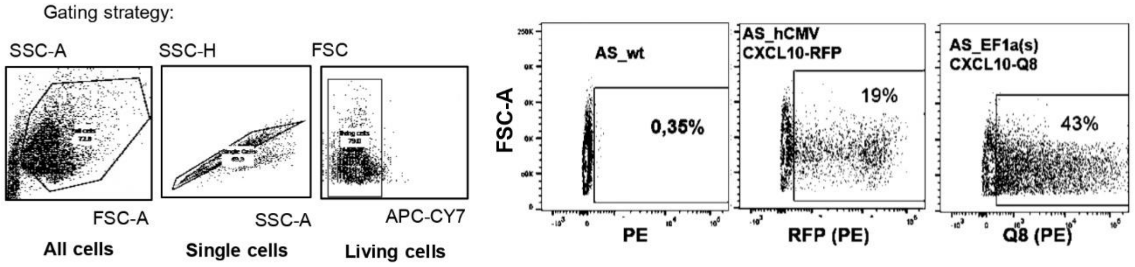

**b**

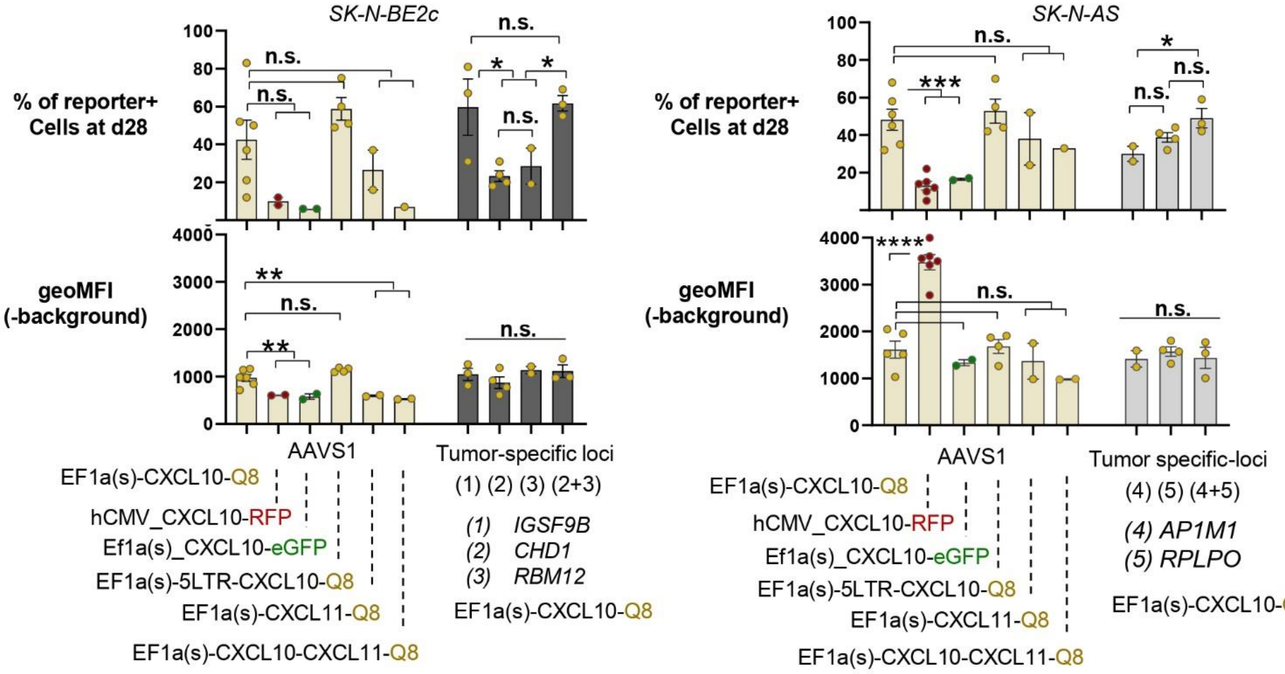

**c**

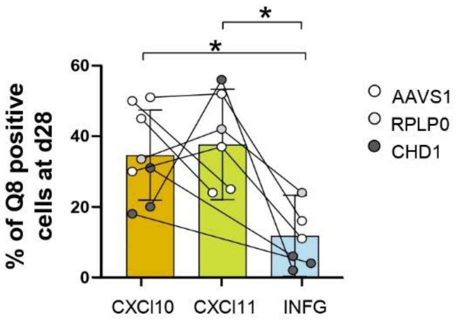

**Supplementary Fig. S10. In vitro CRISPR knock-in. (a)** Exemplary flow cytometry analysis and gating strategies of tumor cell lines 28 days after RNP + dsDNA template electroporation. Q8 reporter positivity was measured using PE-labeled CD34 Monoclonal Antibody QBEND/10 (Thermo Fisher). **(b)** Knock-in rates (%) and reporter expression strengths, represented by background-subtracted geometric mean fluorescence intensity (geoMFI), 28 days post-CRISPR KI for different constructs and loci in SK-N-BE2c and SK-N-AS neuroblastoma cell lines. Compared promoters include human CMV and a custom EF1a-derived promoter with or without a lentivirus-derived 5′ long terminal repeat (LTR) sequence. HDRT templates were designed for cytokines with a Q8 reporter and for CXCL10 with red (RFP) or green fluorescent protein (GFP) reporters. Additional tests included a tri-cistronic vector with two cytokines and a combinatorial knock-in using two HDRTs (differing homology arms) and guide RNAs (two tumor-specific loci) simultaneously without increasing total DNA or gRNA dose. In SK-N-BE2c, CHD1 and RBM12 were targeted; in SK-N-AS, AP1M1 and RPLP0. **(c)** Cumulative knock-in rate comparison for cytokine transgenes across three loci: AAVS1 (SK-N-BE2c and SK-N-AS), CHD1 (SK-N-BE2c only) and RPLP0 (SK-N-AS only). *Data presentation: (b, c) Means ± SD. Statistical analysis: (b) Kruskal-Wallis with Dunn's post hoc test; (c) Mixed-effects analysis with Tukey's multiple comparison test. p values: *<0.05, **<0.01, ***<0.001, ***<0.0001; n.s., not significant.*

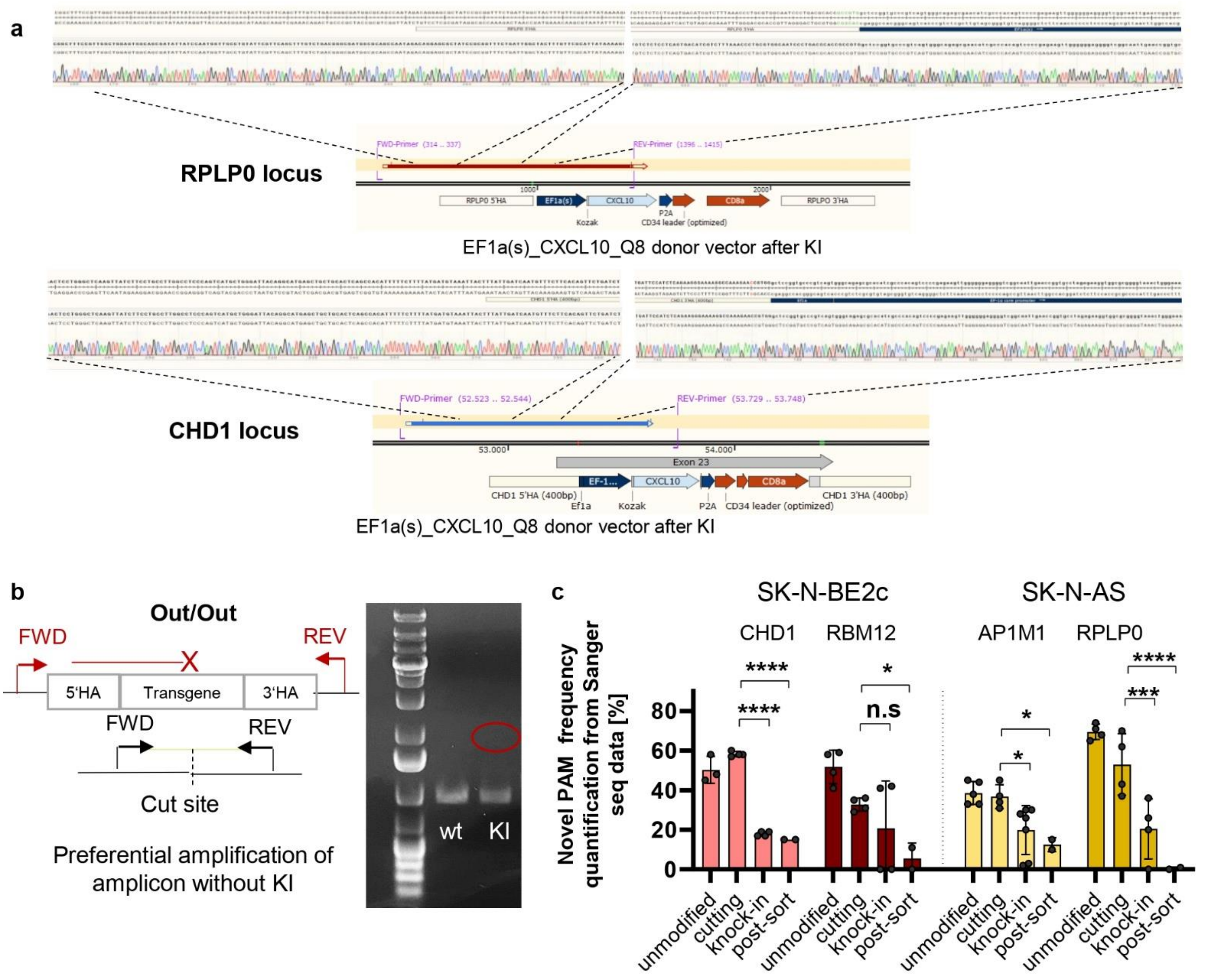

**Supplementary Fig. S11. Site-specific knock-in confirmation via Sanger sequencing and PAM frequency analysis.** *(a)* Schematic representation of on-target knock-in confirmation of an EF1a(s)_CXCL10_Q8 donor vector via PCR and Sanger sequencing with sequence alignment using SnapGene at the RPLP0 (SK-N-AS) and CHD1 (SK-N-BE2c) loci. **(b)** Schematic illustrating the preferential amplification of the non-knock-in amplicon using an Out/Out PCR, where forward and reverse primers are located upstream and downstream of the homology arms and cut site, complemented by representative gel electrophoresis results. **(c)** Quantification of novel PAM frequency from Sanger sequencing data using EditR at the CHD1, RBM12, AP1M1 and RPLP0 loci in SK-N-BE2c and SK-N-AS cell lines. Data are shown for unmodified cells, cells treated with RNP only (cutting), and cells treated with RNP+HDRT (knock-in), with or without enrichment for reporter-positive cells. *Data presentation: (c) Means ± SD. Statistical analysis: (c) Two-way ANOVA with Dunnets post hoc test. p values: *<0.05, **<0.01, ***<0.001, ****<0.0001; n.s., not significant.*

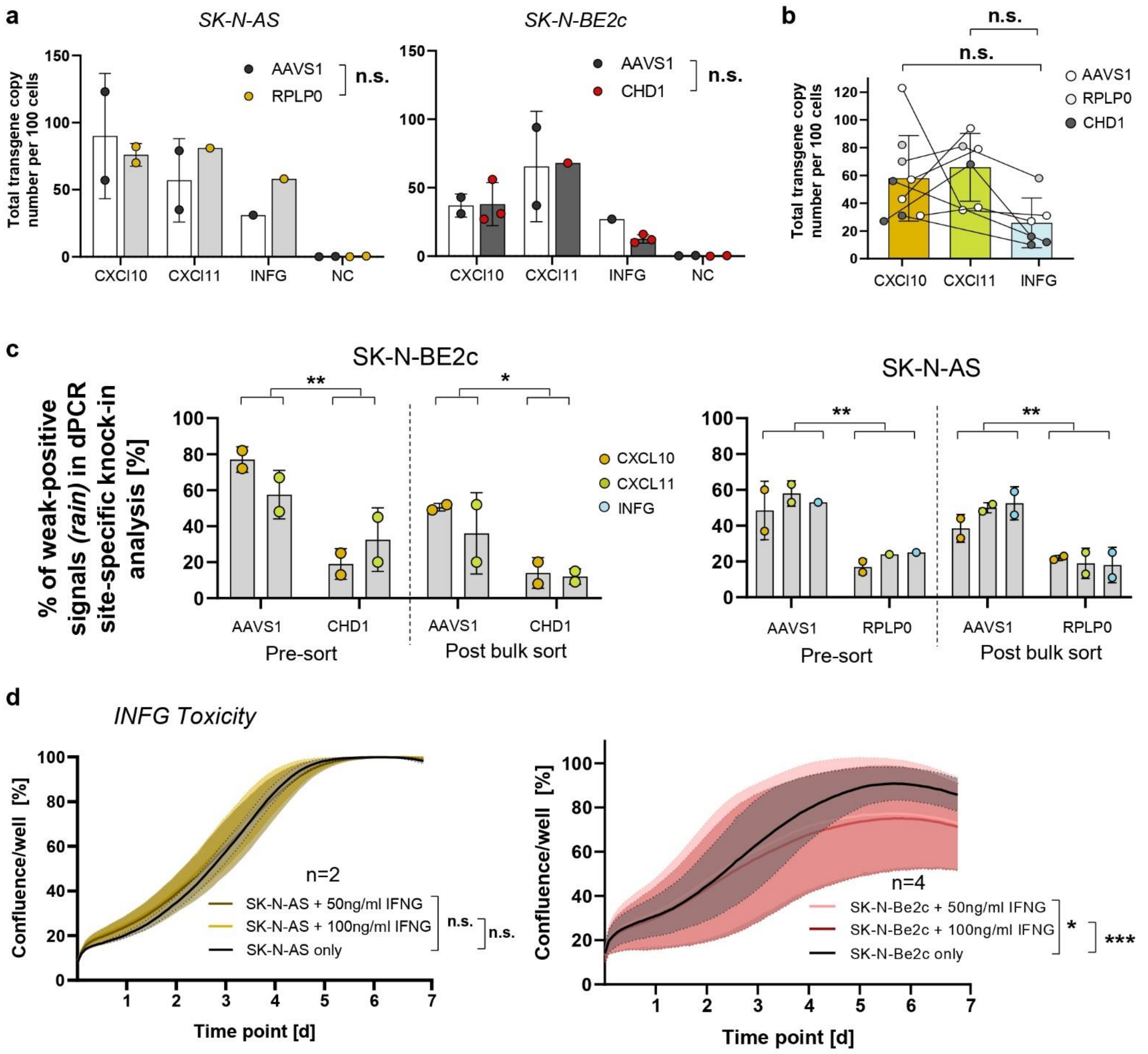

**Supplementary Fig. S12. Site-specific knock-in analysis via dPCR and IFNG toxicity testing. (a)** Total transgene copy number per 100 cells, 7 days after RNP/HDRT electroporation in SK-N-AS and SK-N-BE2c cell lines for different transgenes and target loci. **(b)** Paired analysis of cumulative total transgene copy number for the different cytokine transgenes CXCL10, CXCL11, IFNG. **(c)** Quantification of weak-positive signals ("rain") in digital PCR (dPCR) Out/In site-specific knock-in analysis for SK-N-BE2c and SK-N-AS neuroblastoma cell lines after HDRT knock-in at the AAVS1 control locus or novel PAM sites, before and after enrichment for reporter-positive cells.. **(d)** 7 day growth curves of unmodified SK-N-AS and SK-N-BE2c cell lines cultivated with varying concentrations of IFNG represented by confluence development recorded using Incucyte live imaging. *Data presentation: (a-d) Means ± SD. Statistical analysis: (a-d) Two-Way-ANOVA with Tukey test. p values: *<0.05, **<0.01, ***<0.001, ****<0.0001; n.s., not significant.*

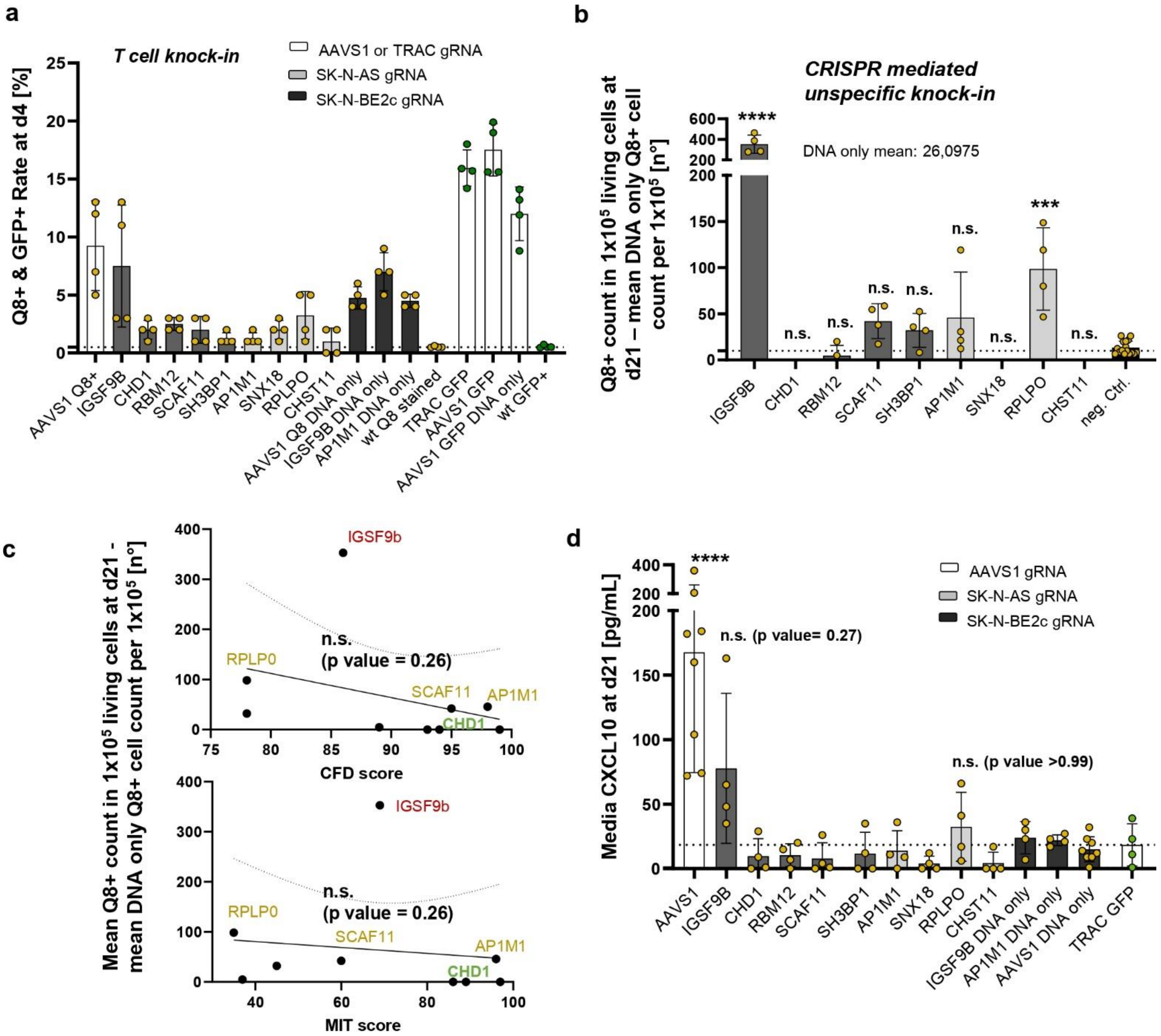

**Supplementary Fig. S13. Unspecific T cell knock-in. (a)** Comparison of the percentage of reporter (Q8/GFP)-positive cells 4 days after RNP/HDRT electroporation in primary T cells from two donors using gRNAs targeting neuroblastoma-specific novel PAM sites or AAVS1 and TRAC control loci, measured via flow cytometry. **(b)** Analysis and comparison of CRISPR-mediated unspecific knock-in rates, calculated as the $Q8^{+}$ count per 100,000 cells minus the mean $Q8^{+}$ count per 100,000 cells in DNA-only control samples, via high-throughput flow cytometry (>300,000 cells analyzed per sample) for Q8-reporter harboring HDRTs targeted at neuroblastoma-specific PAM sites. **(c)** Correlation of the mean CRISPR-mediated unspecific T cell knock-in rates, analyzed via high-throughput flow cytometry and DNA-only background subtraction (b), with CRISPR efficiency scores (CFD and MIT). **(d)** Quantification of secreted CXCL10 two days after media change at day 21 post-RNP/HDRT electroporation in primary T cells from two donors using gRNAs targeting neuroblastoma-specific novel PAM sites or AAVS1 and TRAC control loci via ELISA. *Data presentation: (a,b,d) Means ± SD. Statistical analysis: (b,d) Kruskal-Wallis with Dunn's post hoc test; (c) Linear regression for curve fitting and Spearman correlation. p values: *<0.05, **<0.01, ***<0.001, ****<0.0001; n.s., not significant.*

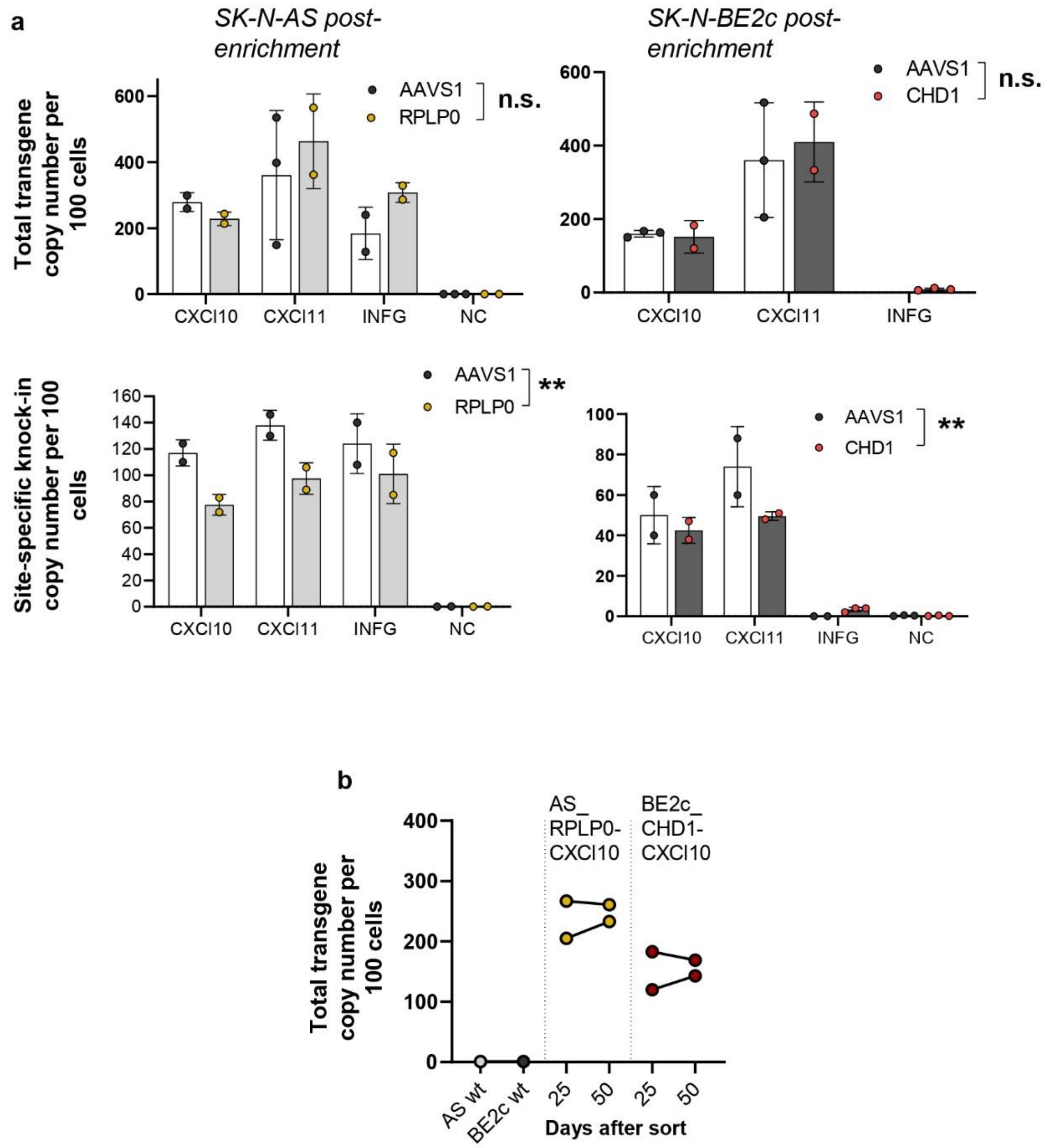

Supplementary Fig. S14. dPCR analysis after FACS enrichment. (a) Total transgene copy number and site-specific knock-in copy number per 100 cells after enrichment of SK-N-AS and SK-N-BE2c knock-in cell lines for different transgenes and target loci, determined via digital PCR (dPCR). (b) Total transgene copy number 25 and 50 days after enrichment in SK-N-AS CXCL10 RPLP0 and SK-N-BE2c CXCL10 CHD1 knock-in cell lines. *Data presentation: (a) Means ± SD. Statistical tests: (a) Two-way ANOVA. p values: *<0.05, **<0.01, ***<0.001, ****<0.0001; n.s.: not significant.*

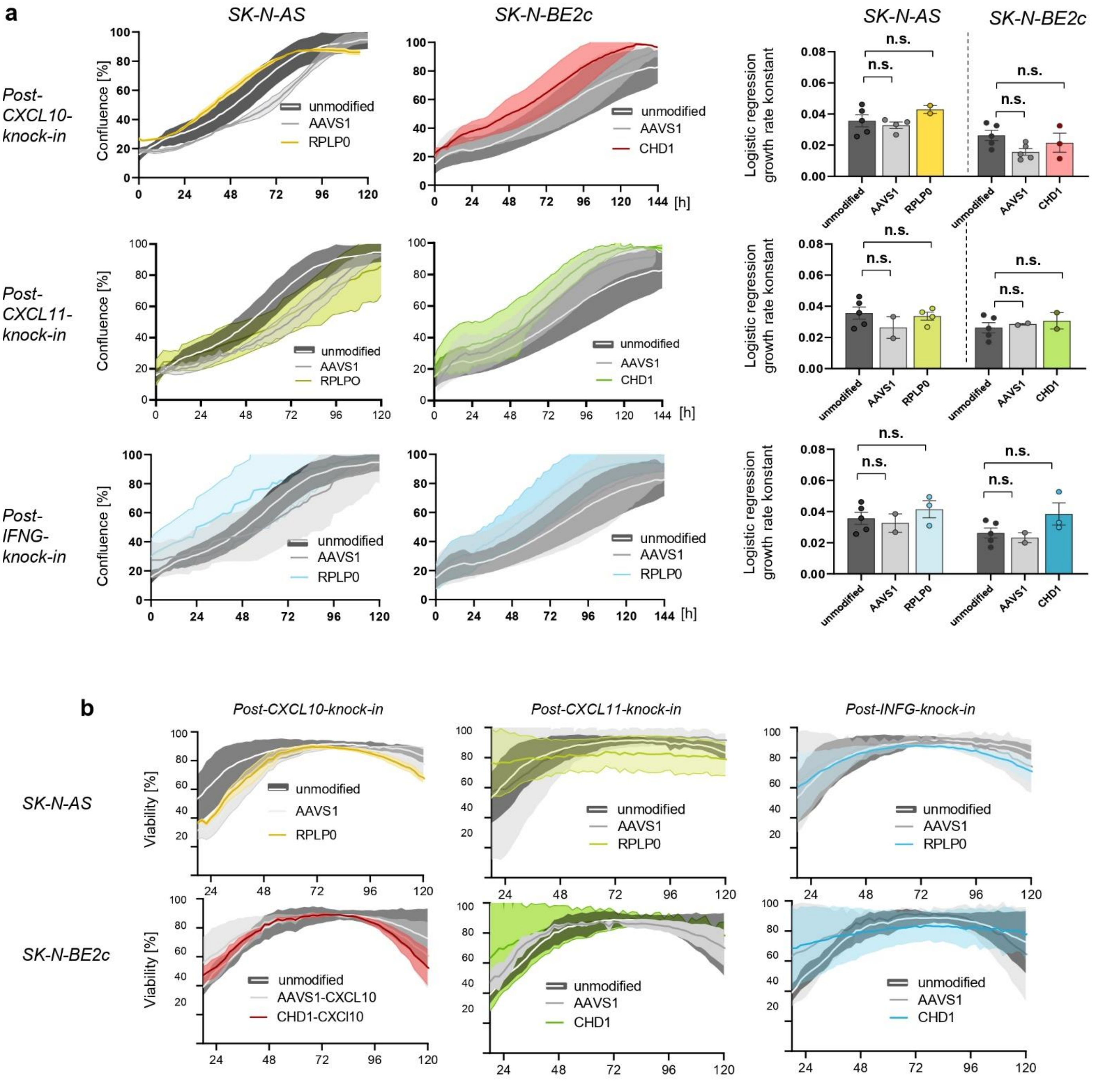

Supplementary Fig. S15. Characterization of enriched transgenic tumor cell lines. (a) Growth curves of unmodified or enriched SK-N-AS and SK-N-BE2c knock-in cell lines, represented by confluence development recorded using Incucyte live imaging, followed by growth rate comparison. (b) Viability curves of unmodified or enriched SK-N-AS and SK-N-BE2c knock-in cell lines, measured using Incucyte Cytotox Dye and Incucyte live imaging. *Data presentation: (a,b) Means ± SD. Statistical analysis: (a) Kruskal-Wallis with Dunn's post hoc test; n.s., not significant.*

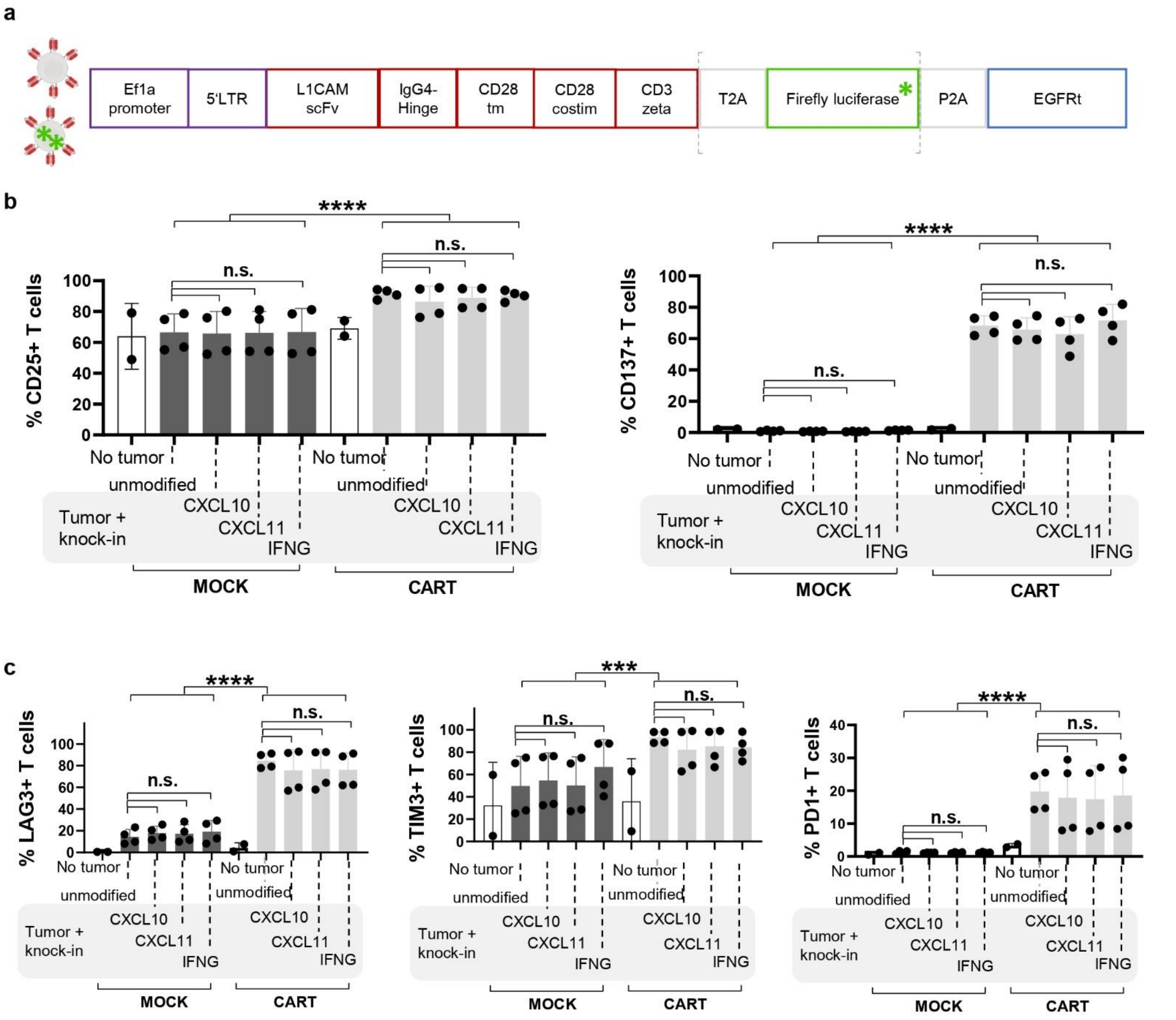

**Supplementary Fig. S16. CAR T cell activation and exhaustion in tumor cell line–CAR T cell co-culture experiments. (a)** Schematic overview of the two CAR constructs used in the study, featuring an EF1α promoter followed by a lentiviral 5′ long terminal repeat (LTR), a codon optimized L1CAM-targeting single-chain variable fragment (scFv), a short 12 amino acid long spacer domain from the immunoglobulin G4 (IgG4) Fc hinge, a CD28 transmembrane (tm) domain, a CD28 costimulatory (costim) domain, a CD3-zeta signaling domain, a T2A linker, firefly luciferase, a P2A linker and a truncated non-functional epidermal growth factor receptor (EGFRt). **(b)** T cell activation, represented by CD25 and CD137 positivity, measured via flow cytometry 24 hours after co-culture initiation (see Fig. 4) for different tumor cell line–CAR T cell combinations. **(c)** T cell exhaustion, represented by LAG3, TIM3 and PD1 positivity, measured via flow cytometry under the same conditions. *Data presentation: (a-c) Means ± SD. Statistical analysis: (a-c) Kruskal-Wallis with Dunn's post hoc test. p values: *<0.05, **<0.01, ***<0.001, ***<0.0001; n.s., not significant.*

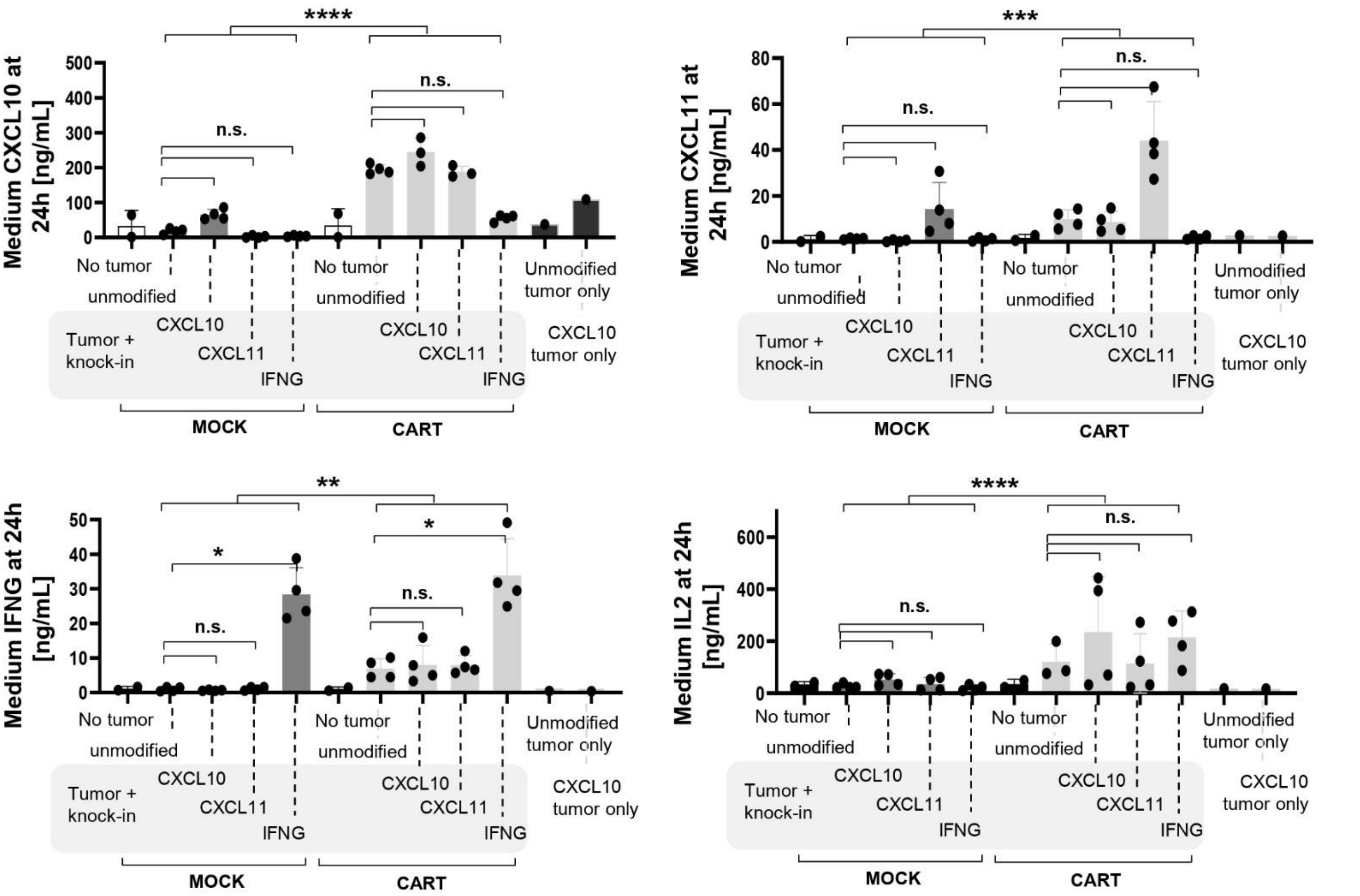

Supplementary Fig. S17. Cytokine secretion in tumor cell line–CAR T cell co-culture experiments. CXCL10, CXCL11, IFNG and IL2 cytokine concentrations in the supernatant, determined via ELISA 24 hours after co-culture initiation. *Data presentation: Means ± SD. Statistical analysis: Kruskal-Wallis with Dunn's post hoc test. p values: *<0.05, **<0.01, ***<0.001, ***<0.0001; n.s., not significant.*

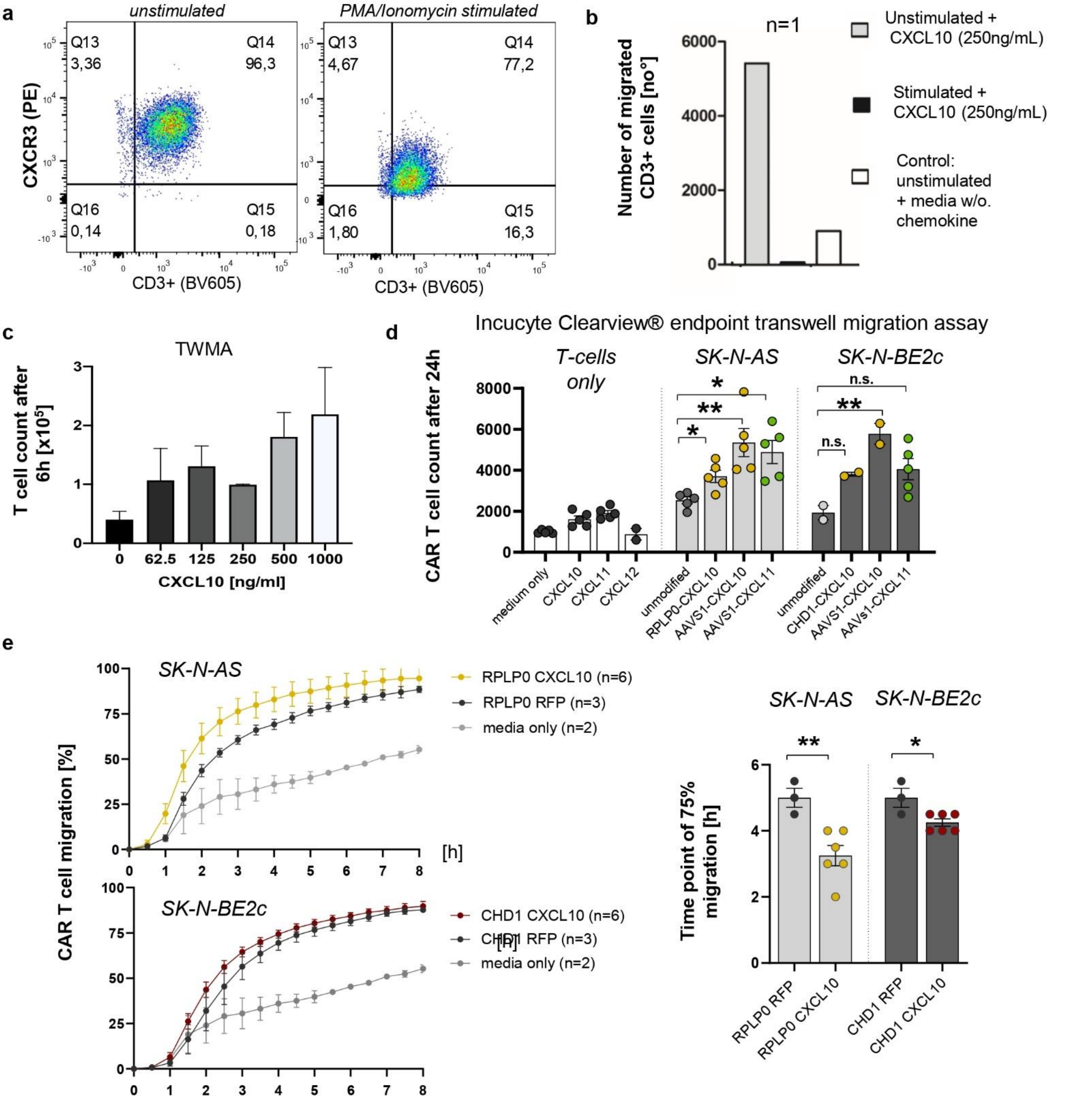

**Supplementary Fig. S18. Cytokine-mediated T cell migration in vitro. (a)** Representative flow cytometry data showing CXCR3 expression, the receptor for CXCL10 and CXCL11, on unstimulated and PMA (Phorbol 12-Myristate 13-Acetate)/Ionomycin-stimulated T cells. **(b)** Preliminary Boyden chamber transwell migration assay (Corning) comparing the number of migrated $CD3^+$ T cells between unstimulated and PMA/Ionomycin-stimulated conditions. **(c)** Boyden chamber transwell migration assay comparing the number of migrated $CD3^+$ T cells in response to increasing CXCL10 concentrations. **(d)** Incucyte Clearview endpoint transwell migration assay (Sartorius) measuring the number of migrated L1CAM-targeting $CD3^+$ CAR T cells after 24 hours in either media only, recombinant cytokines, or supernatants from unmodified or knock-in SK-N-AS and SK-N-BE2c cell lines. **(e)** Migration curves generated using a Boyden chamber transwell migration assay (Corning) and Incucyte live imaging (Sartorius) to determine the migration rate of $CD3^+$ CAR T cells in response to supernatants from transgenic SK-N-AS and SK-N-BE2c cell lines (CXCL10-expressing or RFP) or media only, followed by statistical comparison of the time points at which 75% of migration was reached. *Data presentation: (c-e) Means ± SD. Statistical analysis: (d, e) Kruskal-Wallis with Dunn's post hoc test. p values: *<0.05, *<0.01; n.s., not significant.*

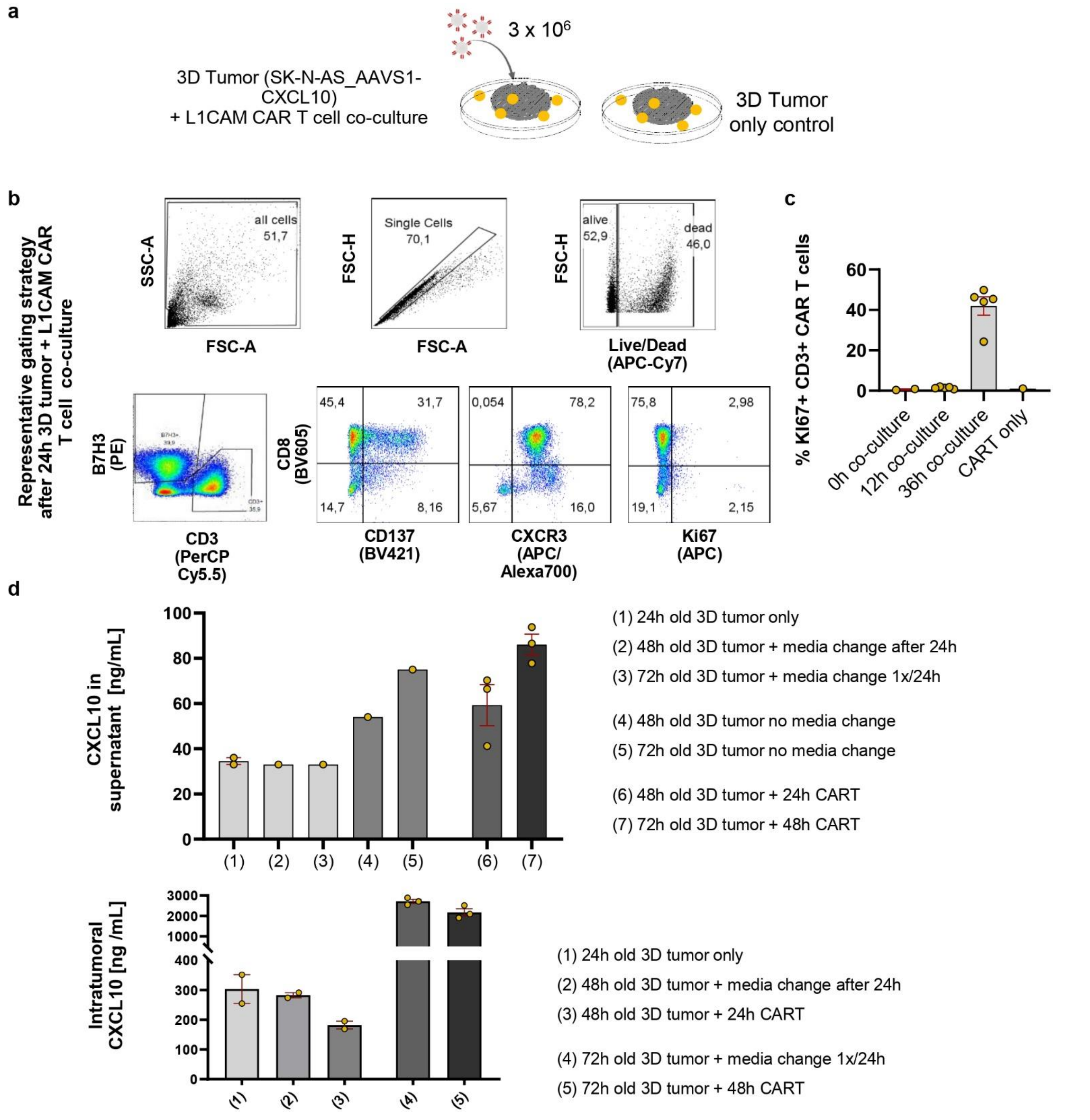

**Supplementary Fig. S19. 3D tumor infiltration assay.** (a) Schematic overview of the preliminary 3D bioprinted SK-N-AS tumor model and L1CAM-targeting CAR T cell co-culture. (b) Representative flow cytometry gating strategy 24 hours after 3D bioprinted SK-N-AS tumor–L1CAM CAR T cell co-culture, distinguishing T cells and tumor cells via B7H3 and CD3 surface markers. (c) KI67 proliferation marker positivity in CD3⁺ CAR T cells at different time points following 3D tumor–CAR T cell co-culture initiation. (d) CXCL10 concentration measured by ELISA in the supernatant or within the 3D tumor construct under various co-culture conditions. *Data presentation: (c, d) Means ± SD.*

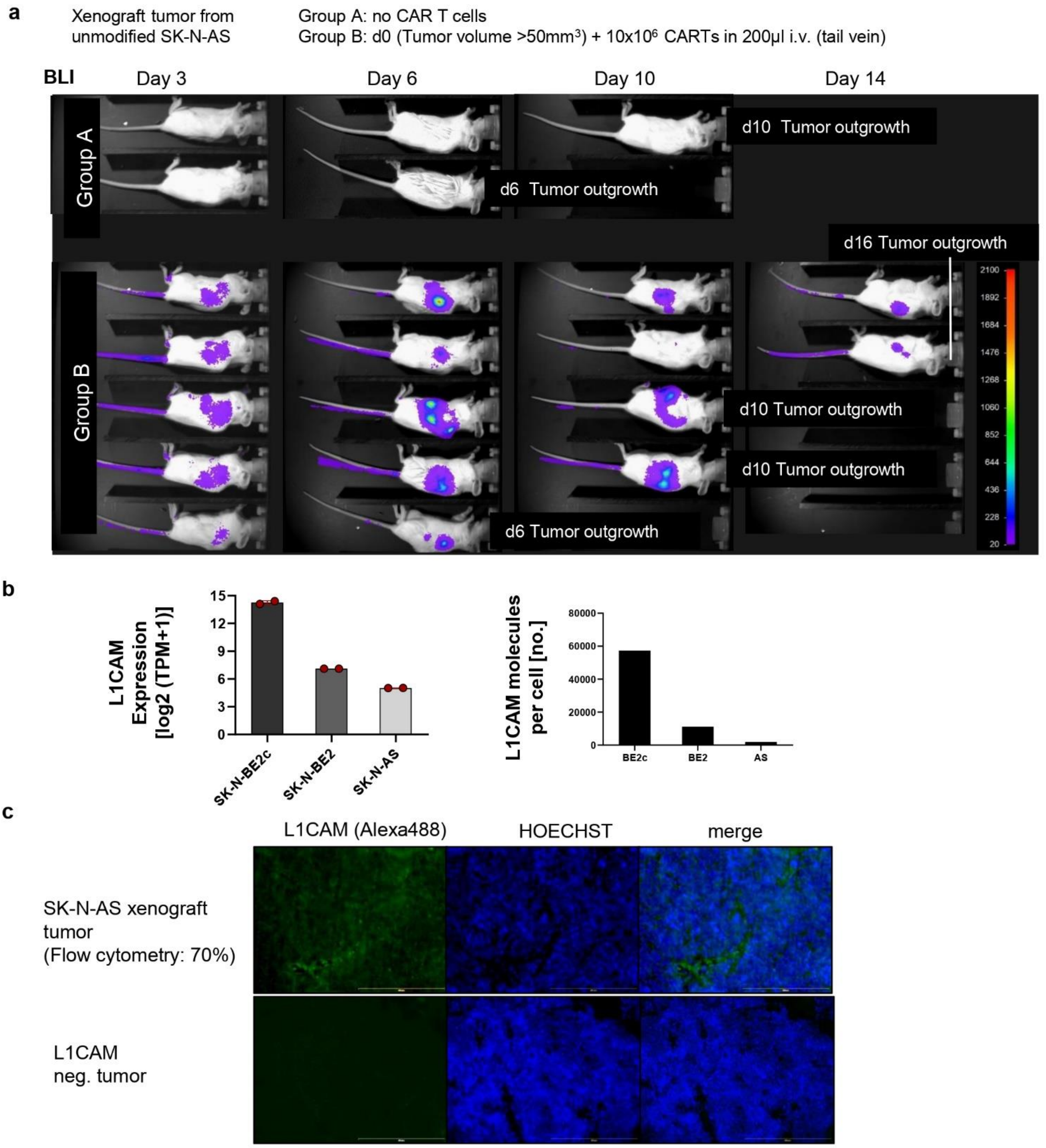

**Supplementary Fig. S20. In vivo CAR T cell infiltration model development.** (a) Preliminary in vivo CAR T cell infiltration analysis in a xenograft mouse model using unmodified SK-N-AS neuroblastoma cells and L1CAM-targeting, firefly luciferase-expressing CAR T cell treatment after tumor engraftment (tumor size >50 mm³), followed by regular monitoring and bioluminescence imaging to assess CAR T cell migration, tumor infiltration and expansion. (b) L1CAM expression levels from DepMap 24Q2 and 24Q4 datasets and L1CAM molecule per cell count as determined via QuantiBRITE PE bead surface expression analysis. (c) Immunofluorescence staining of two different neuroblastoma PDX mouse models, one L1CAM-positive and one L1CAM-negative. *Data presentation: (b) Means ± SD.*

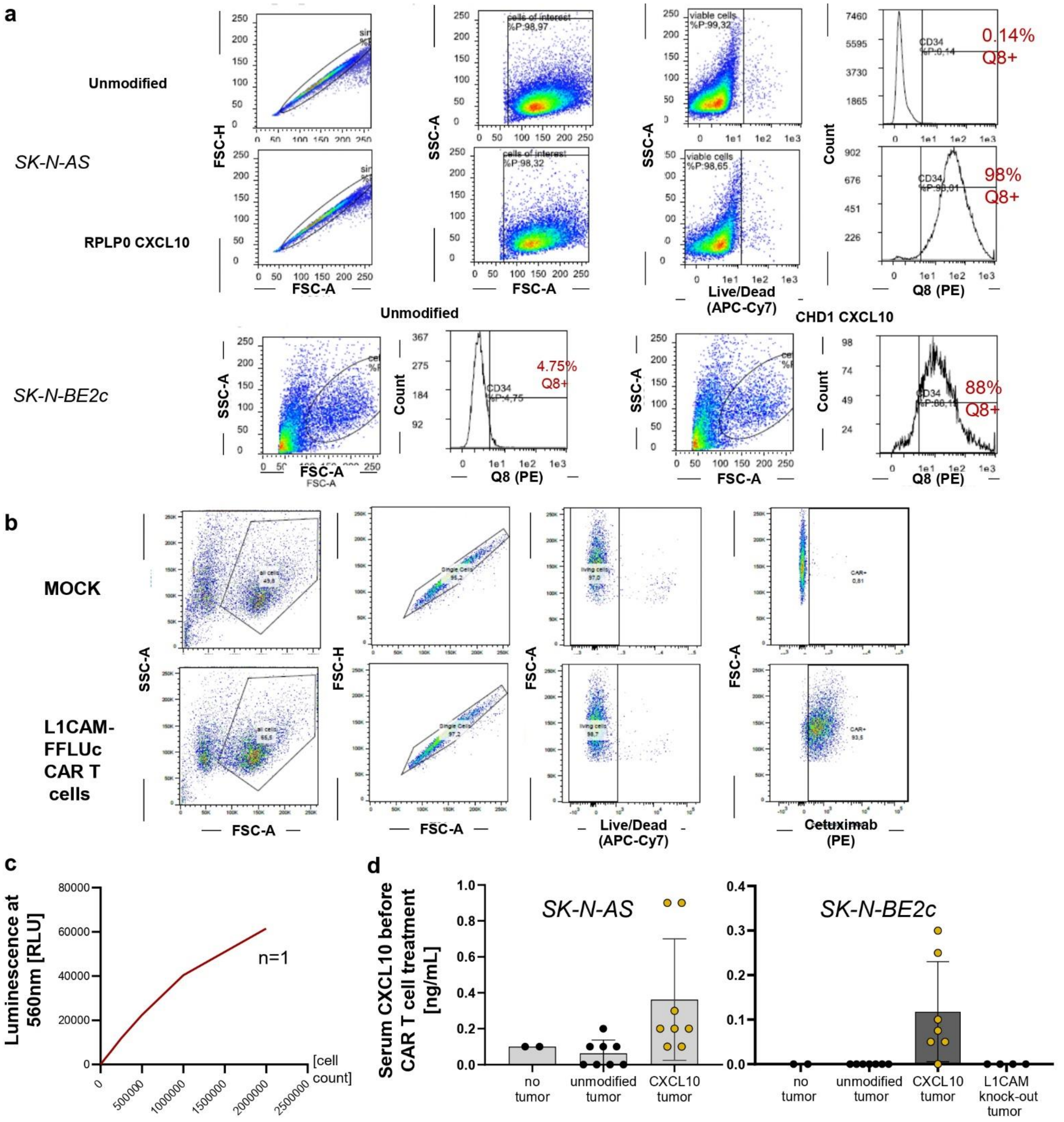

**Supplementary Fig. S21. In vivo CAR T cell infiltration experiment: pre-transplant and pre-treatment analysis.** (a) Confirmation of transgene expression in SK-N-AS and SK-N-BE2c neuroblastoma cell lines, assessed via Q8 positivity using flow cytometry. (b) CAR expression confirmation before in vivo application, determined by flow cytometry in CAR T cells after transduction with an L1CAM CAR, firefly luciferase and EGFRt reporter-expressing lentiviral vector, followed by expansion. Cetuximab staining was used for EGFRt detection on the cell surface. (c) In vitro bioluminescence intensity measured for different L1CAM-targeting, firefly luciferase-expressing CAR T cell counts after luciferin incubation. (d) Serum CXCL10 levels on day 0, measured by ELISA in NSG mice harboring unmodified or CXCL10-expressing SK-N-AS or SK-N-BE2c xenograft tumors, directly before CAR T cell treatment. *Data presentation: (b) Means ± SD.*

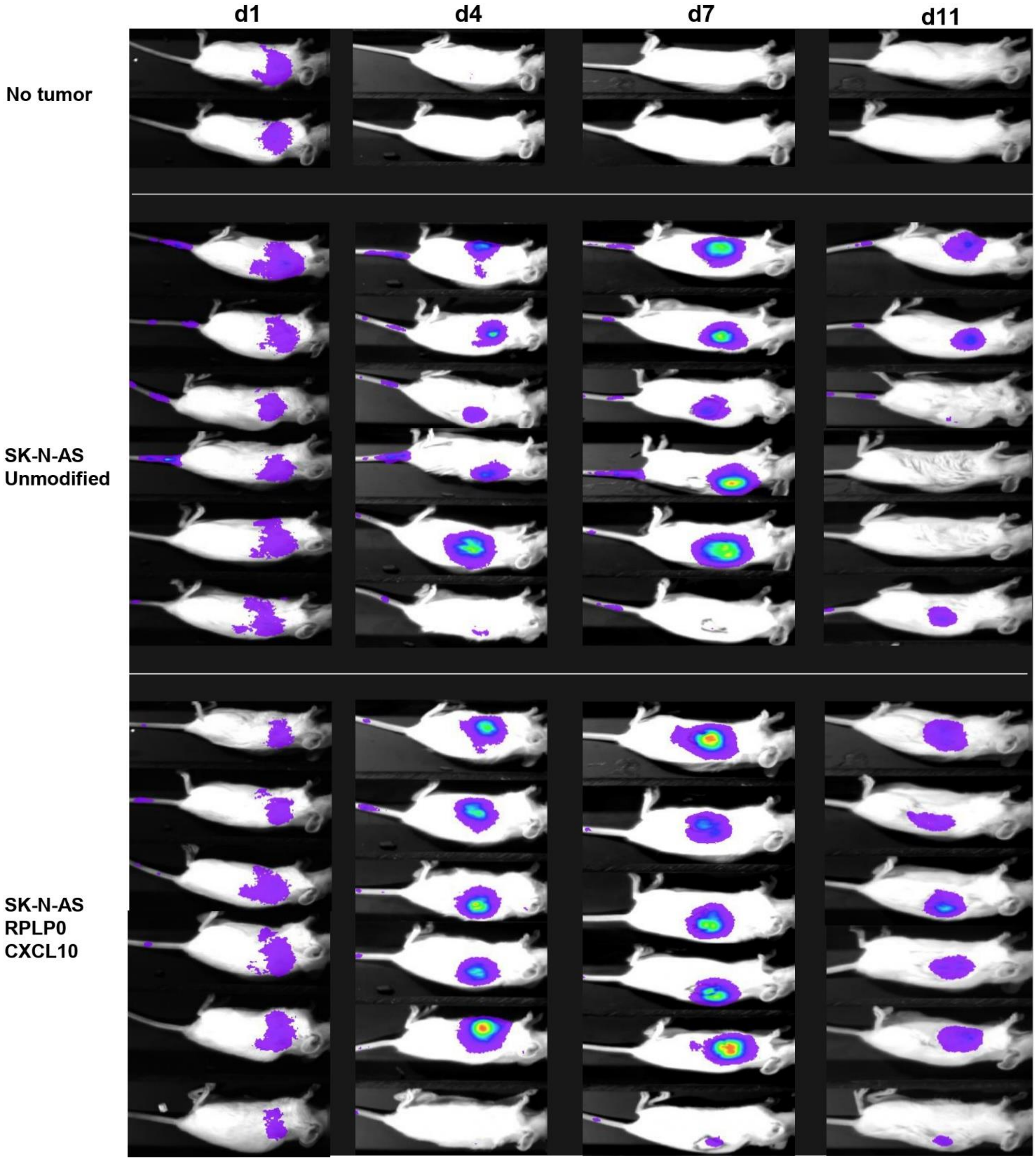

**Supplementary Fig. S22. Bioluminescence imaging of CAR T cell treated SK-N-AS xenograft neuroblasoma models.** Imaging timepoints are given relative to treatment with L1CAM targeting, luciferase expressing human CAR T cells. Mice with tumor volumes of >1200-1500mm$^3$ were removed from the study. *Data from d14 to d30 is not shown.*

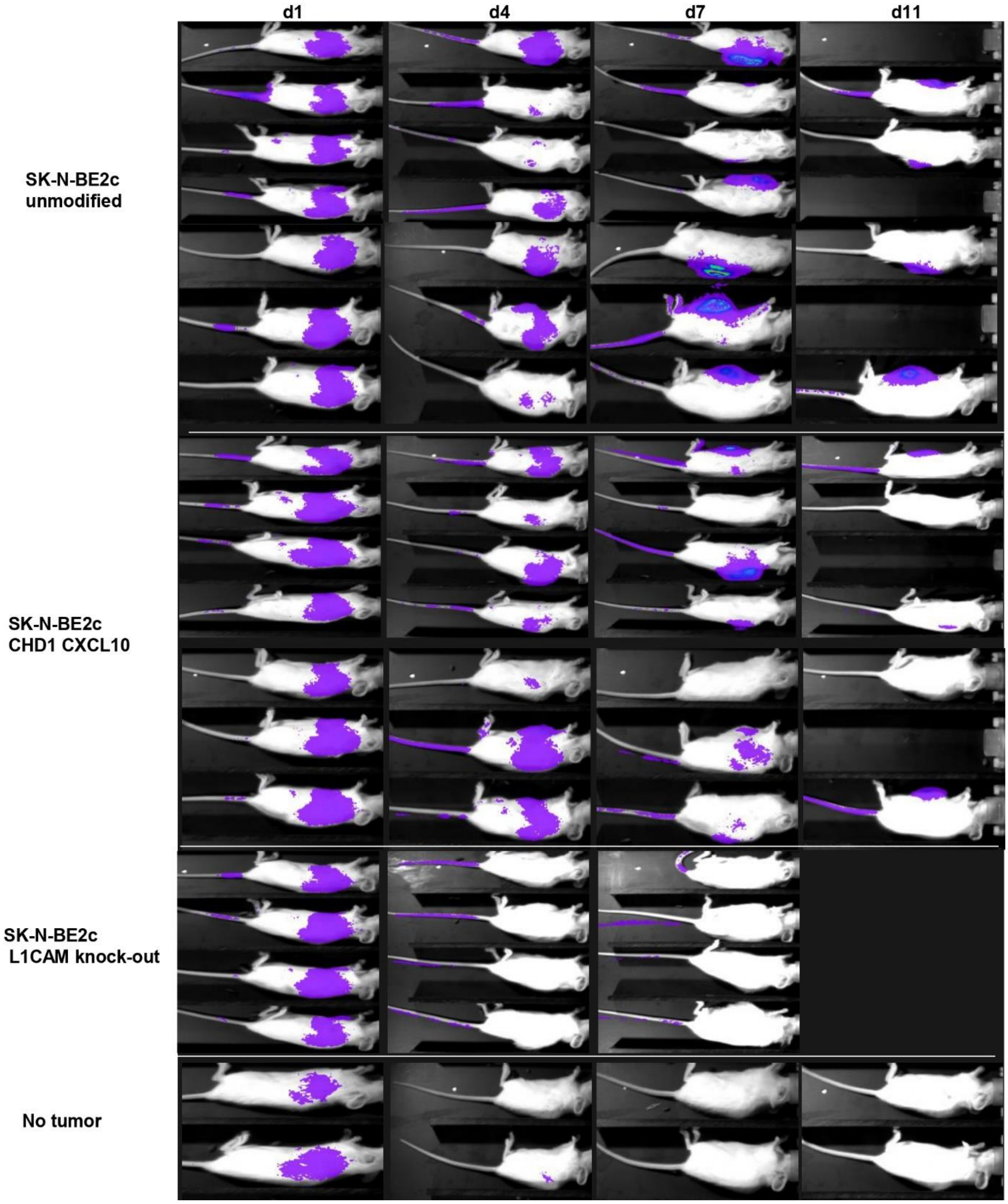

**Supplementary Fig. S23. Bioluminescence imaging of CAR T cell treated SK-N-BE2c xenograft neuroblasoma models.** Imaging timepoints are given relative to treatment with L1CAM targeting, luciferase expressing human CAR T cells. Mice with tumor volumes of >1200-1500mm$^3$ were removed from the study. *Data from d14 to d30 is not shown.*

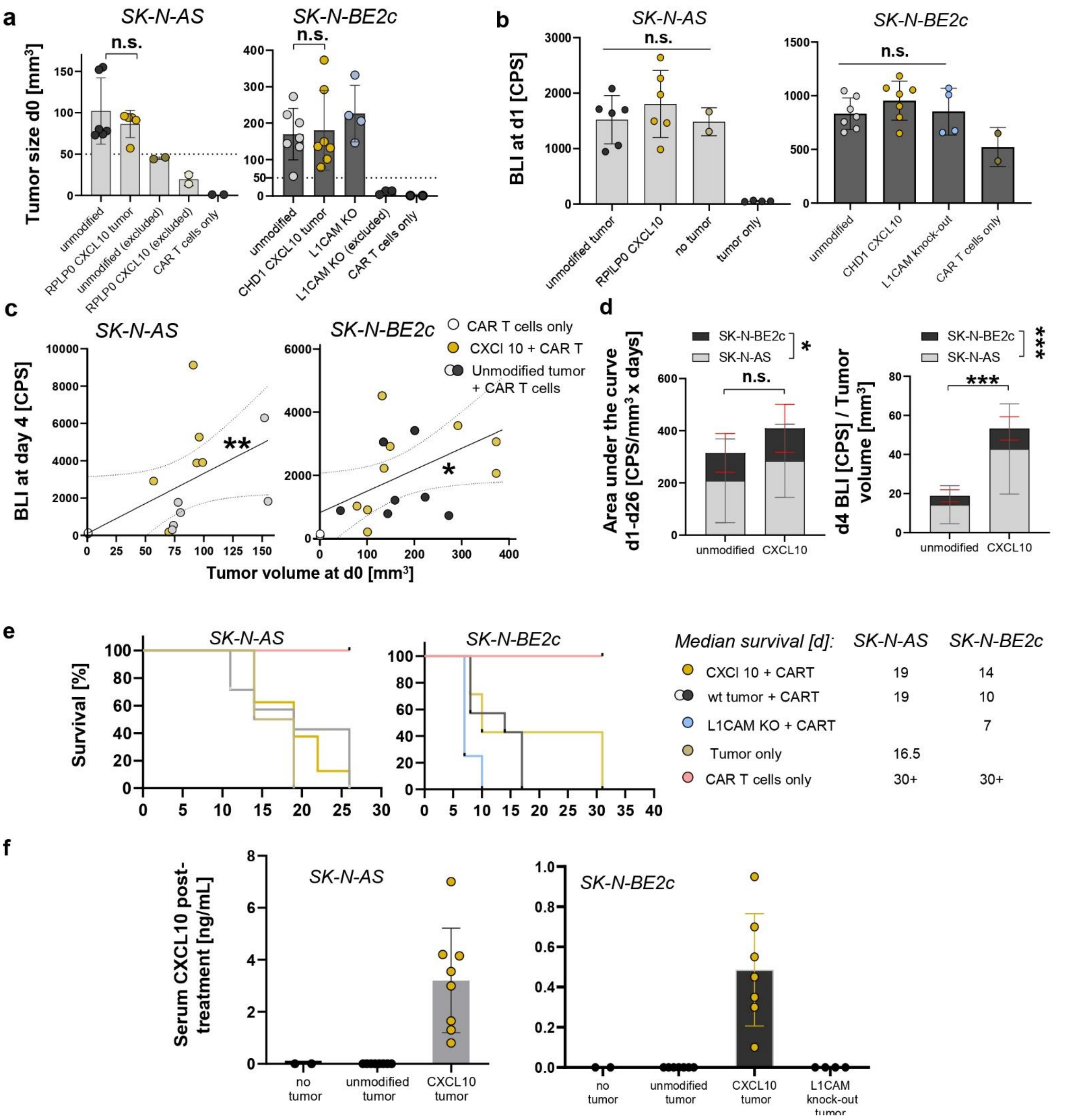

**Supplementary Fig. S24. In vivo CAR T cell infiltration in CXCL10-secreting xenograft tumors. (a)** Manually measured tumor size on the day of CAR T cell injection (d0), compared across treatment groups; only animals with tumors bigger than 50 mm³ received CAR T cell treatment. **(b)** Bioluminescence measurement in the lung on day 1 after treatment with L1CAM-targeting, firefly luciferase-expressing CAR T cells. **(c)** Correlation between the bioluminescence signal in the left flank on day 4 and tumor volume at day 0. **(d)** Cumulative comparison of total tumoral CAR T cell infiltration and expansion, represented by the area under the curve (AUC) of the bioluminescence-to-tumor volume ratio and early infiltration based on the bioluminescence-to-tumor volume ratio on day 4, combining SK-N-AS and SK-N-BE2c data. **(e)** Survival curves for different SK-N-AS and SK-N-BE2c xenograft mouse model treatment groups after CAR T cell treatment (d0), including median survival. **(f)** Serum CXCL10 levels measured on the day of tumor outgrowth and animal removal after CAR T cell treatment via ELISA. *Data presentation: (a, c, d, f) Means ± SD; (E) Kaplan-Meier survival curves. Statistical analysis: (a,c) Kruskal-Wallis with Dunn's post hoc test; (b) Linear regression for curve fitting and Spearman correlation. (d) Two-way ANOVA. p values: *<0.05, **<0.001; n.s., not significant.*

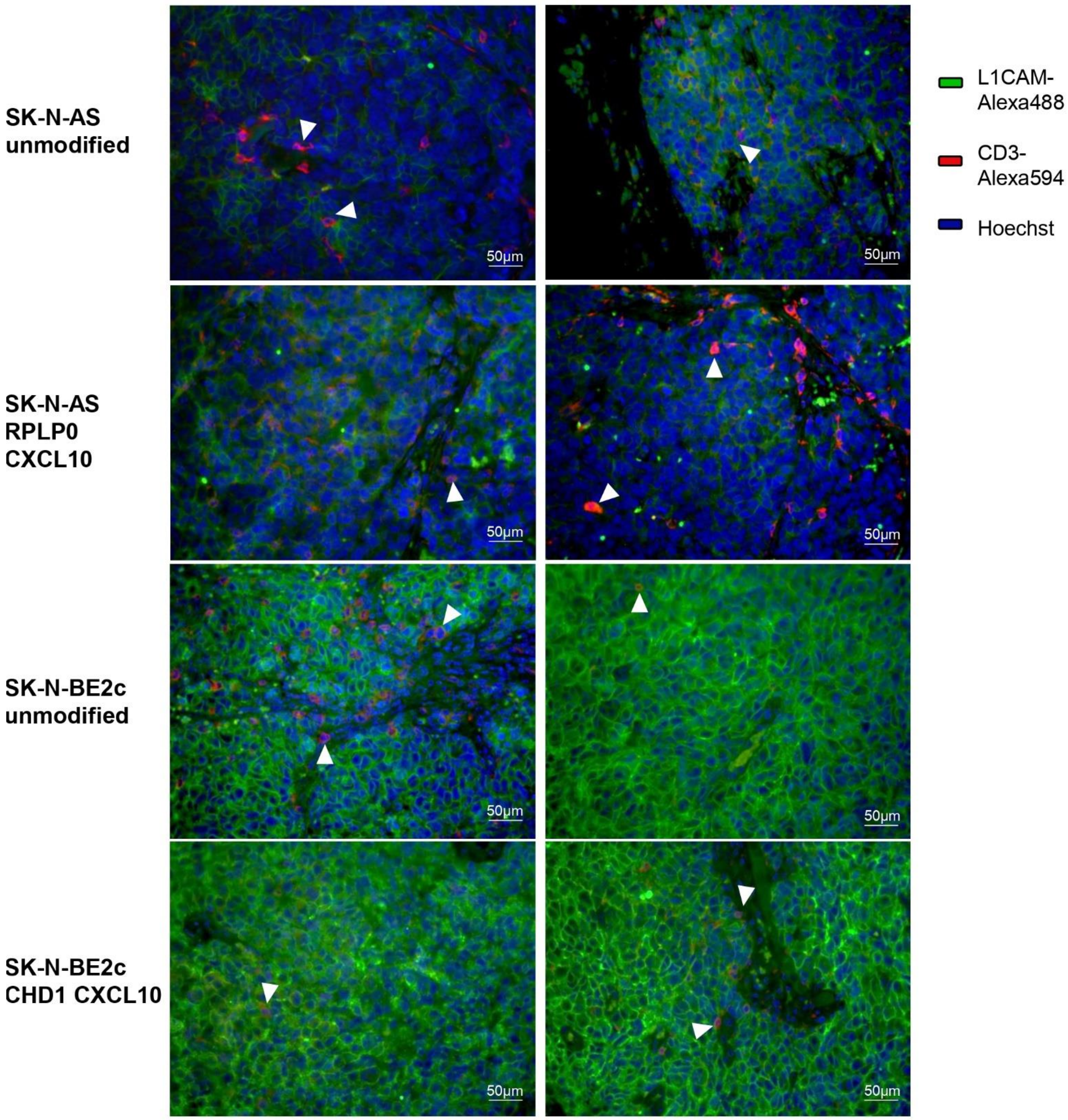

**Supplementary Fig. S25. Immunofluorescence staining of tumor sections.** Representative 40x magnification images of L1CAM and CD3 immunofluorescence staining of tumor sections at the time point of outgrowth and animal removal after treatment with L1CAM-targeting, firefly luciferase-expressing CAR T cells. White arrows indicate examples of intratumoral CD3⁺ T cells.

#### Supplementary Tables

##### Supplementary Table 1. Exemplary patient data for top and bottom rank novel PAM sites identified via CancerPAM

| | Gene | Location | Variant | Expression [log 2 + 1 norm] | gRNA + PAM [5'-3'] | CRISPR efficiency (Doench / Moreno Mateos) | CRISPR specificity (CFD / MIT) | rated position | copy number |
|---|---|---|---|---|---|---|---|---|---|
| Patient A | | | | | | | | out of 110 | |
| | QKI | Chr6:163575669 | T>C | 11.1 | accagacaccgattgtgtgt AGG | 69 / 44 | 94 / 89 | 1 | 2 |
| | SARM1 | Chr 17:28392620 | A>G | 11.9 | tggactctcgaggcttgagt CGG | 62 / 50 | 91 / 85 | 2 | 5 |
| | PIN1 | Chr19:9846446 | C>G | 9.1 | cacaagactgtcaggcggag GGG | 62 / 61 | 86 / 84 | 3 | 3 |
| | EMCN | Chr4:100684664 | A>C | 7.6 | cagctttgttctttctactt AGG | 33 / 43 | 27 / 1 | 96 | 2 |
| Patient B | | | | | | | | out of 289 | |
| | GDAP1L! | Chr20:44250939 | T>C | 10.5 | aggcacgttgtatcaaacat CGG | 64 / 62 | 95 / 91 | 1 | 3 |
| | CDYL | Chr6:4844537 | A>G | 10.7 | cacgctgctctgtccgtccg AGG | 49 / 50 | 94 / 88 | 2 | 3 |
| | PTCD3 | Chr2:86133390 | T>C | 11.8 | tagccgattggccacatcca AGG | 49 / 59 | 93 / 82 | 3 | 3 |
| | LINC02253 | Chr15:97289166 | T>G | 0 | tattaacttgattttttaaa AGG | 19 / 23 | 23 / 10 | 263 | 1 |

##### Supplementary Table 2. Neuroblastoma cell line target loci

| Gene | Variant | Variant RNA reads | gRNA + PAM [5'-3'] | PAM Copy number | Expres-sion [TPM] | depMap CRISPR score | Doench | Moreno | CFD | MIT | CancerPAM Rank |
|---|---|---|---|---|---|---|---|---|---|---|---|
| SK-N-AS | | | | | | | | | | | out of 406 |
| AP1M1 | T>G | 59% | CCGCCAGCGCCGTCTACGTG CGG | 0.9 | 12.5 | -0.06 | 67 | 67 | 98 | 96 | 6 |
| SNX18 | C>G | 22% | GCGCTACAAGCACTTCGACT GGG | 0 | 9.5 | -0.02 | 52 | 52 | 99 | 97 | 8 |
| RPLP0 | A>G | 62% | GCCGTGATGCCCAGGGAAGA CGG | 2 | 16.5 | -1,6 | 47 | 47 | 78 | 35 | 304 |
| CHST11 | T>C | 50% | TTGACATCGTCCCCTTTGCG CGG | 1 | 9.5 | 0 | 32 | 32 | 93 | 86 | 34 |
| SK-N-BE2c | | | | | | | | | | | out of 426 |
| IGSF9B | C>G | 99% | ACGGCAGCCTGACAGTGACA TGG | 2.4 | 14 | 0.05 | 37 | 37 | 86 | 69 | 104 |
| CHD1 | A>C | 33% | GAATAGTCCGTGGTCTTCCA CGG | 1 | 12.5 | 0.23 | 44 | 44 | 94 | 89 | 36 |
| RBM12 | T>G | 42% | AATAATCCATCACCCAGTGT AGG | 1.2 | 12.5 | -0.33 | 37 | 37 | 89 | 37 | 59 |
| SCAF11 | G>C | 33% | GTAAGTGCCTCTGCCACGGT TGG | 0.9 | 12.5 | 0.04 | 60 | 60 | 95 | 60 | 26 |
| SH3BP1 | T>C | 45% | GAATTCTTGGTTGCCTGACT GGG | 0 | 9.5 | -0.06 | 45 | 45 | 78 | 45 | 243 |

#### Supplementary Table 3. HDRT sequences

**AAVS1 5‘homology arm (399bp)**

ACCGTTTTTCTGGACAACCCCAAAGTACCCCGTCTCCCTGGCTTTAGCCACCTCTCCATCCTCTTGCTTTCTTTGCCTGGACACCCCGTTCTCCTGTGGATTCGGGTCACCTCTCACT
CCTTTCATTTGGGCAGCTCCCCTACCCCCCTTACCTCTCTAGTCTGTGCTAGCTCTTCCAGCCCCCTGTCATGGCATCTTCCAGGGGTCCGAGAGCTCAGCTAGTCTTCTTCCTCCA
ACCCGGGCCCCTATGTCCACTTCAGGACAGCATGTTTGCTGCCTCCAGGGATCCTGTGTCCCCGAGCTGGGACCACCTTATATTCCCAGGGCCGGTTAATGTGGCTCTGGTTCTG
GGTACTTTTATCTGTCCCCTCCACCCCACAGTGGGGCCACTAGGGACAG

**AAVS1 3’homology arm (400bp)**

GATTGGTGACAGAAAAGCCCCATCCTTAGGCCTCCTCCTTCCTAGTCTCCTGATATTGGGTCTAACCCCCACCTCCTGTTAGGCAGATTCCTTATCTGGTGACACACCCCCATTTCC
TGGAGCCATCTCTCTCCTTGCCAGAACCTCTAAGGTTTGCTTACGATGGAGCCAGAGAGGATCCTGGGAGGGAGAGCTTGGCAGGGGGTGGGAGGGAAGGGGGGGATGCGT
GACCTGCCCGGTTCTCAGTGGCCACCCTGCGCTACCCTCTCCCAGAACCTGAGCTGCTCTGACGCGGCCGTCTGGTGCGTTTCACTGATCCTGGTGCTGCAGCTTCCTTACACTTC
CCAAGAGGAGAAGCAGTTTGGAAAAACAAAATCAGAATAAGTTGGTCCTGAGTTCT

**AP1M1 5’homology arm (400bp)**

GTCGGTTTCACCACCAGGCTCTCTCGCCTAACTCGGCCCACGGGAGGCCGTTTCCAGCTGCTCTGGCTCCATAGCTCAGCCTGGCAGCGGCCACTGGGTGCCCTGATTTGGTCG
GCCCCGCCCCACCGGGCCTTACTTCCGGGATGCTTCTGGCGGGAGCAAGAAACATGGTTACATACGACGTAATGCGCCTGCGCACAGTCCGTCCTGCGCCTGCGCATTCCTGAC
AGCACCTCCCGGACCGGACATAACGGTCCCCGCGCGGCTCCCCGAACCGGAAGTGGAGGTGAGCTGTCGCGGGCGGCGCCCGGCCTTGCTCAACGCCCAGCAGTCCCCACCG
TCGCTGCCGACGCCACCGCCCTCGACCGCTGCCGAGGCCTCCTGCAGCCATCATGTCCGC

**AP1M1 3’homology arm (400bp)**

GACCTGAAGGGCAAGGTACTGAGGGCTCCCCACCCTCCCTGTTGCCAGGCAACCGGCAGGGGCCTCCGCCCGCGGGGACCCACCCTGTTGCTAGGTAACCGGCTTATGAGCCC
CATCCTGGTGTGAGGCAGAGAGGCCGGTAGACCTCACCTGAGAGCCCCATCCTGTTTCTGGGTAACCGGGCGGGGGTCTTAAGTGGGTAATCCCCGCCCTGTTGCTACGTAAC
GCGGTGGGCTGCAGCCACCGCGTTGCCAGGTAACTGACTGCACCGCCCTCTGCTCCCCAGCGCCGGGGCCAGGGGCTCGGGCCTCGGTGGAGGACCCAGAAGCGGTGCCGCA
GCCGGCCTCGGAGCCAGGCAGGGCGTCCAGGGGCTGCGCGACCCTTGGCCGGGGGTGGCGAG

**SNX18 5’homology arm (400bp)**

CTAGATCGACGGCTCGTCTTCGGCGGGTGTGGGCGCAGCCGGCCGCTACCGCCTGTCCACGCGCTCCGACCTGTCCCTGGGTTCCCGCGGCGGCTCGGTCCCCCCGCAGCACC
ACCCGTCGGGGCCCAAGAGCTCGGCCACCGTGAGCCGCAACCTCAATCGCTTCTCCACCTTCGTCAAGTCCGGCGGGGAGGCCTTCGTGCTGGGGGAGGCGTCAGGCTTCGTG
AAGGACGGGGACAAGCTGTGCGTGGTGCTGGGGCCCTATGGCCCCGAGTGGCAGGAGAACCCCTACCCGTTCCAGTGCACCATCGACGACCCCACCAAGCAGACCAAGTTCA
AGGGCATGAAGAGCTACATCTCCTACAAGCTGGTGCCCACGCACACGCAGGTGCCGGTGCAT

**SNX18 3’homology arm (395bp)**

TGTACGCGCGCCTGGCGGAGAAGTTCCCGGTCATCTCCGTGCCCCACCTGCCCGAGAAGCAGGCCACCGGCCGCTTCGAGGAGGACTTCATCTCTAAGCGCAGGAAGGGCCTG
ATCTGGTGGATGAACCACATGGCCAGCCACCCAGTGCTGGCGCAGTGCGACGTCTTCCAGCACTTCCTGACGTGCCCCAGCAGCACCGACGAGAAAGCCTGGAAGCAGGGCAA
GAGGAAGGCCGAGAAGGACGAGATGGTGGGCGCCAACTTCTTCCTGACCCTTAGCACGCCCCCCGCCGCTGCCCTTGACCTGCAGGAGGTGGAGAGCAAGATCGACGGCTTC
AAGTGCTTCACCAAGAAGATGGACGACAGCGCGCTGCAGCTCAACCACACGGCCAAC

**RPLP0 5’homology arm (400bp)**

ACAGGAGCGCTATCCGCGGTTTCTGATTGGCTACTTTGTTCGCATTATAAAAGGCACGCGCGGGCGCGAGGCCCTTCTCTCGCCAGGCGTCCTCGTGGAAGGTTCGTGTGCTAG
TTAGATGGGCGCCAGGGGTCGCCGGCGGGAAGCATGGAGGGGTCTTTGGGGGCCTTTGGGAACATGGAGTCCTATTCTGTTCCGCCTGGGGCCTCGGTGGCGGCTTGCACGC
CCCGAGATGACGGCCGCTGCCCTAGGCAGGGCCGGCGGGCGATTGCGCGTGTCCTGCTCCTCTTAGGCCCGGGACCGCGGGATGGGTGTCGGCGTGACCAGGCCTGAGCTCC
CTGTCTCTCCTCAGTGACATCGTCTTTAAACCCTGCGTGGCAATCCCTGACGCACCGCCGTG

**RPLP0 3’homology arm (400bp)**

GGCGACCTGGAAGTCCAACTACTTCCTTAAGATCATCGTAAGTGCAGGGTGGGTCGCCTCTGCTCTTCATGTTGCCCCAGCGCAAATAGGGACAGTCAGCTGCTATGTGCTGAG
GGTCTACTCACACCGGCTACTGAATTAGGCCATTTTTGGGAAAATACTGTTTAGTTAACAATTTCCTGAGATAGGTCCCTTCTGTTGCAGATAAACGGGCTCAGGCAAGTTAAGT
GGGTCCTAAGATGACAGCATTCGTATCCAGGTCTGTCTGGCTTCTAAAAGAGCGCGCTTTATACTTTTTATTTTTTAAACGGAGTCTCGCTCTGTTGTCCAGGCTGGAGTGCAGTG
GCTTGATCTTGGCTTACTGCAACCTCCGCTTCCCAGGTTCAAGCGATTCTCCTGC

**CHST11 5’homology arm (400bp)**

AAGCCGTAAGCGGAGGGTGCTGACCCCCAACGACCTGAAGCACTTGGTGGTGGATGAGGACCACGAGCTCATCTACTGCTACGTGCCCAAGGTGGCCTGCACCAACTGGAAGC
GGCTCATGATGGTCCTGACCGGGCGGGGGAAGTACAGCGACCCCATGGAGATCCCGGCCAACGAGGCACACGTCTCCGCCAACCTGAAGACCCTGAACCAGTACAGCATCCCA
GAAATCAACCACCGCTTGAAAAGCTACATGAAGTTCCTGTTTGTCCGGGAGCCCTTCGAGAGGCTAGTGTCCGCCTACCGCAACAAGTTCACCCAGAAGTACAACATCTCCTTCC
ACAAGCGGTACGGCACCAAGATCATCAAACGCCAGCGGAAGAACGCCACCCAGGAGGCC

**CHST11 3’homology arm (400bp)**

ACGATGTCAAATTCGAGGAGTTTGTGGCCTATCTCATCGACCCACACACCCAGCGGGAGGAGCCTTTCAACGAACACTGGCAAACCGTCTACTCACTCTGCCATCCCTGCCACAT
CCACTATGACCTCGTGGGCAAGTACGAGACACTGGAAGAGGATTCTAATTACGTCCTGCAGCTGGCAGGAGTGGGCAGCTACCTGAAGTTCCCCACCTATGCAAAGTCTACGAG
AACTACTGATGAAATGACCACAGAATTCTTCCAGAACATCAGCTCAGAGCACCAAACGCAGCTGTACGAAGTCTACAAACTCGATTTTTTAATGTTCAATTACTCAGTGCCAAGCT
ACCTGAAATTGGAATAAAGGGGGTGGGGAGAGGGAGAGAATCATGCTTTTTAATT

**IGSF9b 5’ homology arm (400bp)**

AAGCCCATTGTCACCTGGCTCAAGGAGGGGACGCTCCTCGGTGCTAGTGGGAAATACCAGGTGAGTGTGGTTCTAGGTTGGGGCCGCTGCGGTCTCTTCCCCCGTTTCCCCAG
GGCAGCCTCCTACTGCAGCCGACCCCTGATGCTACAGGCACAAGCACAGAGGAGGCTGGAAAGCCGTAGCCTGGCTCAGCTGCAGGCGTCTCTGTGTCGCTGGCGGGAGATG
GGCAAGGGGGCACCTGCCCTTCGTGTCCCCCAGATGCATCTGTCCCTGTGGTCATCTCCACCTTAGTGTCTCCGACTGCATCTCCGTGTTTGAGGGTAGCAGCGGTGGGTGTGA
GCGTGGGTGCTGAGCTCCCTCTGTTTTGGGCTCCCAGGTGAGTGACGGCAGCCTGACAGTG

**IGSF9b 3’ homology arm (400bp)**

GTCAGTCGGGAGGACAGAGGTGCCTACACCTGCCGAGCGTACAGCATTCAGGGGGAGGCTGTCCACACGACTCACCTGCTTGTCCAAGGTAGGAGCCATTTCCTCAGCCTCTTT
AACCCTGGGCTCCCGCGGGGATGTCCAGCCCGGCTGAGCTACGGAGGGCGTTTTCCCAGGGCTCAGCTGGGGCTAGTGGAGGTGCAGGCGGCCATGAGGACCTGTGCGGGG
CTCCTGTGTGCTGCCTGCTGCCCTGGTCAGAGCTCCTGCACCGGCTGCCTGGGAGAGGGCCAGGCCCACAGGCCGTGGCCAGGCACCACATGGCGTGGCAGAGAGGCCCCTG
CTGTGTGGGTTTCTGGTTTCTCCTTCTGTTCAGATAGCATCTCTCCCATGTCTCCGCCTCCCT

**CHD1 5’homology arm (400bp)**

TTATTGATCAATGTTTCTTCACAGTTCTGATCTCCTCAGGAAAAATGCCTAGAGGTTATGTTGCTGCGTCAGTAGAATAACTTTTGGAGGCTTTTGATATAATGTCTGATTACCTTT
TAAATCAAAATATCCACCAGAGGTATATATTTTTCTCTGTTTGCTCTTTAATTTTGTTTATGGAATGTTTTTGATACATTTAATAACATTTTTAAGATTCCGACCATCTTTTATAATGG
TTTTTGCTCTTAAGATACAAGCTGTTTTTACACTAAGGTTGGGTAAATATTTTCCAATTCCAGATTAGTTTCAATGGAAGTGAAGGGAGGCGCAGTAGAAGTAGGAGATACTCTG
GATCTGATAGTGATTCCATCTCAGAAGGGAAAAGGCCAAAGAACCGTGG

**CHD1 3’homology arm (400bp)**

AAGACCACGGACTATTCCTCGGGAGAATATTAAAGGATTTAGTGATGCAGAAATTAGGCGGTAAGGTGTTATAAGACAATCGCTTCAACTTGGTTTTAGTAGAGTGTGAAACTA
AAATGTAAAAACGCCGTAATTTTCAAAAATTGTGATGAAGTTTTAGAAGAGGCAAATCTGTTATTTGAGTAGGAATTGGGTGGTTCTGCTTTTGGGTATTTGTTATAACTGAACAA
AACAAGTTTGGTGTCTAATCTTTGCATGTTATTTATTTATCGCTAAACTGCTTAGACTACACAATTTTTTTTATTAATTTGACTCAATAGTGATCCTTTCGAATAATAAAAGTAGTTTT
GATGCATTTGATTTTTAAAAATTCTCACGGTGTTTATGACTCCCAAAGTTA

**RBM12 5’homology arm (400bp)**

TGTGGTTTCAGCGCAGCATGGCTGTGGTCATCCGTTTGCAAGGTCTCCCAATTGTGGCGGGGACCATGGACATTCGCCACTTCTTCTCTGGATTGACCATTCCTGATGGGGGCGT
GCATATTGTAGGGGGTGAACTGGGTGAGGCTTTCATCGTTTTTGCCACTGATGAAGATGCAAGGCTTGGTATGATGCGCACAGGTGGTACAATTAAAGGGTCAAAAGTAACACT
ATTGTTGAGTAGTAAGACGGAAATGCAGAATATGATTGAACTGAGTCGTAGGCGTTTTGAAACTGCCAACTTAGATATACCACCAGCAAATGCCAGTAGATCAGGACCACCACC
TAGCTCAGGAATGAGTAGCAGGGTAAACTTGCCCACAACAGTATCCAACTTTAATAA

| |
|---|
| **RBM12 3'homology arm (400bp)** |
| TACTGCCACCACTTCTGTTCATGAAAGCAACAAAAACATACAGACATTTTCCACAGCCAGCGTAGGAACAGCTCCTCCAAATATGGGGGCTTCCTTTGGGAGCCCAACGTTTAGC<br>TCAACTGTTCCAAGCACAGCCTCTCCAATGAACACAGTCCCGCCGCCACCAATTCCTCCAATTCCAGCGATGCCATCTCTGCCACCAATGCCATCCATTCCCCCAATTCCAGTTCCT<br>CCTCCAGTACCTACATTGCCTCCTGTGCCTCCTGTGCCCCCGATTCCCCCAGTTCCTTCTGTGCCACCCATGACCCCACTGCCACCCATGTCGGGCATGCCGCCCTTGAATCCGCCA<br>CCTGTGGCACCTCTACCTGCTGGAATGAATGGCTCTGGAGCACCTATGAAT |
| **SCAF11 5'homology arm (401bp)** |
| TGTAGAACAAAAAGTAAGAGTTCATCATTTGGTAGAATTGACAGAGATAGTTACTCTCCCCGGTGGAAGGGAAGATGGGCAAATGATGGTTGGAGATGTCCACGAGGAAATGA<br>TCGGTACAGAAAGAATGACCCAGAGAAACAGAATGAAAATACAAGAAAAGAAAAAAATGACATCCATCTAGATGCTGATGATCCAAATTCTGCTGACAAACATAGAAATGACTG<br>TCCCAATTGGATAACAGAAAAAATAAACTCTGGGCCTGATCCAAGAACCAGAAATCCAGAAAAGTTGAAAGAGTCTCATTGGGAAGAAAATAGAAATGAAAATTCAGGAAATTC<br>TTGGAATAAAAACTTTGGTTCTGGTTGGGTATCTAACCGTGGTAGAGGCAGAGCCAACCG |
| **SCAF11 3'homology arm (400bp)** |
| GTGGCAGAGGCACTTACAGAAGTAGTTTTGCCTATAAAGATCAGAATGAAAATCGGTGGCAAAATCGAAAACCCCTCTCAGGGAATTCAAACAGTTCAGGGAGTGAATCTTTCA<br>AGTTTGTGGAACAGCAATCCTATAAGCGAAAAAGTGAACAGGAGTTCTCATTTGATACACCAGCAGATAGATCTGGATGGACATCTGCATCCAGCTGGGCCGTGAGAAAGACTT<br>TGCCAGCAGATGTACAAAACTACTACTCACGACGAGGCAGAAATTCTTCAGGTCCACAGTCTGGATGGATGAAACAAGAGGAGGAAACATCTGGACAGGGTAATGTATTCTTGT<br>TGGTTTATTGATACTGTTTTATTTGAAGAATTATAAATTTAATATGTTATTTGTATGA |
| **SH3BP1 5'homology arm (400bp)** |
| CCACACCAGGAGGATGCTGTGGGGTCTGCTGAGTACAGTGTGTGTACTTGTGCACGCCTGGTCCCTAGGAGCACAGGTGGTTAGCGTATCTGTGAGGGTGTGCCCGCGTGTGT<br>CACAGCTGGTGCTGGGGCAACCCTAAAGCCTGGGATCCTGTAAACAGTCCTGCGGCAGCCCTGTTCTCCCCTCTGTGGAGTGGCTGTGATCACCAAATCTGCCACCAGGGGGCA<br>ACCCAGAGCACTCATGGCCCAGCCTGGGCAGAGGCCCCGGCCAGTGGAGCTCTCAGATCTGCAGCTCTAAATCCTAGAATCCCATGTGGTCTGCTGTATGGAGGGGACACTTG<br>GCTCTGGACATGCAACAGGGCAGGTGACCAGCTCACCTTGGGATCATCTCCAGGCCCAGT |
| **SH3BP1 3'homology arm (400bp)** |
| CAGGCAACCAAGAATTCAGGCAGCAGTCAAGGCCTAGGAGGCAGCCCGGGTAGTCACAGCCATACGACCATGGCCAACAAGGTGGAGACGCTGAAGGAGGAGGAGGAGGAG<br>CTGAAGAGGAAAGTGGAGCAATGCAGGGTGAGGGCCATGGGGGTCCCCTGGATATGTAGGGGTGGCAGGGGTGGGTCCTGGCCCATCTCACAGGCAAGGAAGCTGAAGTTC<br>AGAGAGGTGACATGACTTGCCTAAGGCCACGTGGCTAGGAGCCAGCAGAGCTCCGCCATTCCCTCGTAGCCTCTTCTGTCTGTACCCCATGCATAACTTGGGAAGGCAGAGACT<br>TGGTCTGTTCCCTGAGTCTCCATGTGTTTGTTTGCTGAGCGAGCGAGTGAATGAATAAATGAAG |
| **Ef1a(s) promoter 239bp** |
| GCTCCGGTGCCCGTCAGTGGGCAGAGCGCACATCGCCCACAGTCCCCGAGAAGTTGGGGGGAGGGGTCGGCAATTGAACCGGTGCCTAGAGAAGGTGGCGCGGGGTAAACT<br>GGGAAAGTGATGTCGTGTACTGGCTCCGCCTTTTTCCCGAGGGTGGGGGAGAACCGTATATAAGTGCAGTAGTCGCCGTGAACGTTCTTTTTCGCAACGGGTTTGCCGCCAGAA<br>CACAGGCCGCCACC |
| **CXCL10 (294bp)** |
| ATGAACCAGACCGCCATCCTGATCTGCTGCCTGATCTTCCTGACCCTGAGCGGCATCCAGGGAGTCCCCCTGTCCAGAACAGTGCGGTGCACCTGTATCAGCATCTCTAATCAAC<br>CTGTGAACCCAAGAAGCCTGGAAAAGCTCGAGATCATCCCTGCTTCTCAGTTCTGCCCTAGAGTGGAAATCATCGCCACAATGAAAAAGAAGGGCGAGAAGAGATGTCTGAACC<br>CCGAGAGCAAGGCCATTAAGAACCTGCTGAAGGCCGTGTCCAAAGAGATGAGCAAGCGGAGCCCT |
| **CXCL11 (282bp)** |
| ATGAGTGTGAAGGGCATGGCTATAGCCTTGGCTGTGATATTGTGTGCTACAGTTGTTCAAGGCTTCCCCATGTTCAAAAGAGGACGCTGTCTTTGCATAGGCCCTGGGGTAAAA<br>GCAGTGAAAGTGGCAGATATTGAGAAAGCCTCCATAATGTACCCAAGTAACAACTGTGACAAAATAGAAGTGATTATTACCCTGAAAGAAAATAAAGGACAACGATGCCTAAAT<br>CCCAAATCGAAGCAAGCAAGGCTTATAATCAAAAAAGTTGAAAGAAAGAATTTT |
| **IFNG (498bp)** |
| ATGAAGTACACCTCTTACATTCTGGCATTCCAATTGTGCATTGTGCTGGGAAGCCTCGGGTGCTATTGTCAAGATCCCTACGTGAAAGAGGCTGAGAATCTGAAGAAGTACTTCA<br>ACGCCGGGCACAGTGACGTCGCTGACAACGGTACACTGTTCTTGGGAATCTTAAAGAACTGGAAGGAAGAAAGCGATCGAAAGATCATGCAATCCCAGATCGTGAGCTTCTATT<br>TTAAGCTGTTCAAGAACTTCAAGGACGACCAATCTATACAGAAATCAGTCGAGACAATCAAAGAGGATATGAACGTGAAGTTCTTTAACTCTAATAAGAAGAAGAGGGACGATT<br>TTGAGAAACTTACAAACTACAGCGTCACGGATCTCAACGTACAGCGGAAGGCGATCCACGAGCTTATTCAGGTCATGGCCGAGTTGTCCCCGGCTGCGAAGACTGGCAAACGG<br>AAGCGTTCACAAATGCTTTTCAGAGGGAGACGAGCGTCACAA |
| **GFP (714bp)** |
| GTGAGCAAGGGCGAGGAGCTGTTCACCGGGGTGGTGCCCATCCTGGTCGAGCTGGACGGCGACGTAAACGGCCACAAGTTCAGCGTGTCCGGCGAGGGCGAGGGCGATGCC<br>ACCTACGGCAAGCTGACCCTGAAGTTCATCTGCACCACCGGCAAGCTGCCCGTGCCCTGGCCCACCCTCGTGACCACCCTGACCTACGGCGTGCAGTGCTTCAGCCGCTACCCC<br>GACCACATGAAGCAGCACGACTTCTTCAAGTCCGCCATGCCCGAAGGCTACGTCCAGGAGCGCACCATCTTCTTCAAGGACGACGGCAACTACAAGACCCGCGCCGAGGTGAA<br>GTTCGAGGGCGACACCCTGGTGAACCGCATCGAGCTGAAGGGCATCGACTTCAAGGAGGACGGCAACATCCTGGGGCACAAGCTGGAGTACAACTACAACAGCCACAACGTC<br>TATATCATGGCCGACAAGCAGAAGAACGGCATCAAGGTGAACTTCAAGATCCGCCACAACATCGAGGACGGCAGCGTGCAGCTCGCCGACCACTACCAGCAGAACACCCCCAT<br>CGGCGACGGCCCCGTGCTGCTGCCCGACAACCACTACCTGAGCACCCAGTCCGCCCTGAGCAAAGACCCCAACGAGAAGCGCGATCACATGGTCCTGCTGGAGTTCGTGACCG<br>CCGCCGGGATCACTCTCGGCATGGACGAGCTGTACAAG |
| **P2A (66bp)** |
| GGCAGTGGAGCAACCAACTTCAGCCTGCTGAAGCAGGCAGGCGACGTGGAAGAAAATCCAGGACCT |
| **Q8 reporter tag (417bp)** |
| ATGGGATTGGTTCGCCGAGGAGCGCGAGCAGGTCCCCGGATGCCCCGCGGATGGACTGCCCTGTGCCTGCTTTCCCTCCTTCCAAGCGGCTTCATGGCAGAATTGCCAACTCAA<br>GGTACTTTCTCAAATGTTTCCACGAACGTTAGTCCAGCTAAACCCACTACAACACCTGCACCTAGACCCCCGACGCCAGCCCCTACTATAGCTTCTCAACCACTGAGTCTTCGGCC<br>AGAGGCATGCAGGCCCGCAGCAGGTGGGGCAGTGCACACAAGGGGTCTCGATTTTGCTTGTGACATTTATATTTGGGCTCCTTTGGCAGGGACCTGTGGGGTACTTCTGCTGA<br>GTCTGGTGATCACGTTGTATTGTAACCACAGGAATAGGCGGCGGGTTTGTAAATGTCCACGACCGGTTGTCTGA |
| **sPA (49bp)** |
| AATAAAAGATCTTTATTTTCATTAGATCTGTGTGTTGGTTTTTTGTGTG |

*HDRT: Homology Directed Repair Template, bp: base pairs*

#### Supplementary Table 4. Primer and probe sequences

| HDRT amplification primers | Orientation | Sequence |
|---|---|---|
| AAVS1_5‘HA_FWD | FWD | ACCGTTTTTCTGGACAACCCC |
| AAVS1_3‘HA_REV | REV | AGAACTCAGGACCAACTTATTCTGATTTTGT |
| AP1M1_5‘HA_FWD | FWD | GTCGGTTTCACCACCAGGC |
| AP1M1_3‘HA_REV | REV | CTCGCCACCCCCGGC |
| CHD1_5‘HA_FWD | FWD | TTATTGATCAATGTTTCTTCACAGTTCTGATCTCC |
| CHD1_3‘HA_REV | REV | TAACTTTGGGAGTCATAAACACCGTGAG |
| CHST11_5'HA_FWD | FWD | AAGCCGTAAGCGGAGGGT |
| CHST11_3'HA_REV | REV | GCATGATTCTCTCCCTCTCCCC |
| IGSF_5'HA_FWD | FWD | AAGCCCATTGTCACCTGGC |
| IGSF_3'HA_REV | REV | AGGGAGGCGGAGACATGG |
| RPlP0_5'HA_FWD | FWD | AGCGCTATCCGCGGTTTCT |

| | | |
|---|---|---|
| RPlP0_5'HA_REV | REV | GCAGGAGAATCGCTTGAACCTG |
| RBM12_5'HA_FWD | FWD | TGTGGTTTCAGCGCAGCATG |
| RBM12_3'HA_REV | REV | ATTCATAGGTGCTCCAGAGCCATT |
| SCAF_5'HA_FWD | FWD | GGTCACCCAACAAAGACACTCCAC |
| SCAF_3'HA_REV | REV | TCATACAAATAACATATTAAATTTATAATTCTTCAAATAAAACAGTATCAATAAACCAACAAG |
| SH3BP1_5'HA_FWD | FWD | CCACACCAGGAGGATGCTG |
| SH3BP1_3'HA_REV | REV | CTTCATTTATTCATTCACTCGCTCGCTC |
| SNX18_5'HA_FWD | FWD | CTAGATCGACGGCTCGTCTTC |
| SNX18_3'HA_REV | REV | GTTGGCCGTGTGGTTGAGC |
| PAM frequency/ CRISPR cutting (out/out) | Orientation | Sequence |
| AAVS1_PAM freq. /Cutting_FWD | FWD | GGCATCTCTCCTCCCTCA |
| AAVS1_PAM freq./Cutting_REV | REV | TGGGGACTAGAAAGGTGAAG |
| AP1M1_PAM freq./Cutting_FWD | FWD | TCCAAACGCCAAAACGGAGA |
| AP1M1_PAM freq./Cutting REV | REV | CGGCATTAGTAGGTGCTCCC |
| CHD1_PAM freq./Cutting_FWD | FWD | GCTCAAGTTATCTTCCTGCCTTGGC |
| CHD1_PAM freq./Cutting_REV | REV | GCCTGGCCAATCTTACCTCATCTC |
| CHST11_PAM freq./Cutting_FWD | FWD | AGCCAAGTGGAAATGTTTCAGG |
| CHST11_PAM freq./Cutting_REV | REV | AGAATGGTGGCATCCATACACA |
| GBA2_PAM freq./Cutting_FWD | FWD | TGCCTGGGAATATGGGGAGA |
| GBA2_PAM freq./Cutting_REV | REV | CTAGTCCACACCCAACACCC |
| IGSF9b_PAMfreq./Cutting_FWD | FWD | CCCTGATGCTACAGGCACAA |
| IGSF9b_PAM freq. _REV | REV | CACAGGTCCTCATGGCCG |
| IGSF9b_Cutting_REV | REV | CTCTGCCAAATGCCCACTGT |
| RBM12_PAM freq. /cutting_FWD | FWD | GCAAGGCTTGGTATGATGCG |
| RBM12_PAM freq. _REV | REV | GGAGGAACTGGAATTGGGGG |
| RBM12_Cutting_REV | REV | GGCAGGGCTAACTTCCACAT |
| RPLP0_PAM freq./Cutting_FWD | FWD | AATTATGTCTCAGCTCCACGTCAT |
| RPLP0_PAM freq./Cutting_REV | REV | CTCAAAGGCCATCCAAAAGTCA |
| SCAF11_PAM freq. /Cutting_FWD | FWD | GTCCACGAGGAAATGCGGT |
| SCAF11_PAM freq._REV | REV | AAGTCTTTCTCACGCCCAG |
| SCAF11_Cutting_REV | REV | TTTGGCAAGGTTTGGAGCAG |
| SNX18_PAM freq./Cutting_FWD | FWD | GAGCTCTCATTCACCGCACT |
| SNX18_PAM freq./Cutting_REV | REV | GATGTCCGGGAAGTTAGCCA |
| **dPCR Primers and probes: Transgene CNV (In/In) Assays** | | |
| CXCL10_Transgene_FWD | FWD | GGAAAAGCTCGAGATCATCC |
| CXCL10_Transgene_REV | REV | TCTTTTTCATTGTGGCGATGATT |
| CXCL10_Transgene_Probe | Probe | 6FAM-CTGCTTCTCAGTTCTGCCCTAGAGTGG-BHQ1 |
| CXCL11_Transgene_FWD | FWD | CTACAGTTGTTCAAGGCTTC |
| CXCL11_Transgene_REV | REV | GGTACATTATGGAGGCTTTC |
| CXCL11_Transgene_Probe | Probe | 6FAM-ACGCTGTCTTTGCATAGGCCC-BHQ1 |
| IFNG_Transgene_FWD | FWD | AAGGAAGAAAGCGATCGAAA |
| IFNG_Transgene_REV | REV | ATAGATTGGTCGTCCTTGAA |
| IFNG_Transgene_Probe | Probe | 6FAM-TGCAATCCCAGATCGTGAGCTTCT-BHQ1 |
| **dPCR Primers and probes: Site-specific knock-in (Out/In) Assays** | | |
| Ef1a_ Out/In _REV | REV | GCACTTATATACGGTTCTCC |
| AAVS1_Out/In_FWD | FWD | GCCGTCTTCACTCGCTG |
| AAVS1_Out/In _Probe | Probe | 6FAM-TCCCTTGCGTCCCGCCTCCCCTT-BHQ1 |
| CHD1_Out/In_FWD | FWD | GAACTCCTGGGCTCAAGTTATC |
| CHD1_Out/In_Probe | Probe | 6FAM-ATGCTGGGATTACAGGCATGAGCT-BHQ1 |
| RPLP0_Out/In_FWD | FWD | TATCCAATGGTTGCCTGTAT |
| RPLP0_ Out/In_Probe | Probe | 6FAM-CTATTGGCTGCGCCATCGCCCGT-BHQ1 |
| **dPCR Primers and probes: Endogenous Control Assay** | | |
| AFF3 FWD | FWD | CACCTAGCATGTGTGGCATT |
| AFF3 REV | REV | GCAGATCCAGGTCGTTGAAG |
| AFF3 probe | Probe | HEX-AACAACTCTTTCTGTCCCCCT-BHQ1 |

*HDRT: Homology Directed Repair Template, FWD: forward, REV: reverse, freq.: Frequency, dPCR: digital PCR, BHQ1: Black Hole Quencher 1.*

#### Supplementary Table 5. Equipment and Consumables used

| Item | Catalogue n° | Provider |
|---|---|---|
| Equipment | | |
| 4D-Nucleofector® core unit | | Lonza Bioscience |
| 4D-Nucleofector® X Unit | | Lonza Bioscience |
| C1000 Touch Thermal cycler | | Bio-Rad |
| Centrifuge 5415R | | Eppendorf GmbH |
| Centrifuge 5424 | | Eppendorf GmbH |
| Centrifuge 5427R | | Eppendorf GmbH |
| Centrifuge 5810 R | | Eppendorf GmbH |
| Duomax 1030, platform shaker | | Heidolph instruments |
| Electronic Balance ABS 80-4 | | Kern & Sohn GmbH |

| | | |
|---|---|---|
| Electronic Rotary Microtome HM340E | | Biotek |
| Erlenmeyer flasks | | Thermo Scientific |
| FlowCytometer LSR-Fortessa X-20 | | BD Biosciences |
| Fortessa Aria Cell sorter | | BD Bioscience |
| Fortessa X-20 | | BD Bioscience |
| Freezer (-20°C) LCv4010 | | Carl Roth |
| Freezer (-80°C), HeraFreeze T series HFU400TV63 | | Thermo Scientific |
| GloMaxR Multi | | Promega |
| Heat controlled pressure cooker | | Dako |
| Ice machine Manitowoc SOTTO | | Manitowoc |
| Incubator HERAcell 240i CO2 | | Thermo Scientific |
| Laminar Airflow Bench HERA safe 2020 KSP18 | | Thermo Scientific |
| Luminometer GloMaxR- Multi+Microplate Multimode Reader with InstinctR E8032 | | Promega |
| Magnetic Shaker RH basic | | IKA Laboratoy Equipment |
| Manual system microscope Olympus BX43 | | Olympus |
| Microscope Axio Vert.A1 | | Zeiss |
| Millipore Barnstead MicroPure | | Thermo Scientific |
| NanoDrop 2000 | | Thermo Scientific |
| Neubauer counting chamber | | Brand |
| PCR workstation Pro | | VWR Peqlab |
| pH-meter Five Easy Le409 | | Mettler Toledo |
| Pipette filler pipetusR | | Hirschmann Labortechnik |
| Pipette multipipetteR stream | | Eppendorf GmbH |
| Pipettes (2.5 – 1000 µl) | | Eppendorf GmbH |
| QuadroMACS | | Miltenyi |
| Qubit fluorometer | | Thermo Scientific |
| QIAcuity Digital PCR System (6-plex) | | Qiagen |
| Shandon Excelsior ES | | Thermo Scientific |
| Spectrophotometer EPOCH | | BioTek Instruments |
| StepOnePlus Real-Time PCR system | | Thermo Scientific |
| Suction pump AC02 | | Carl Roth |
| Table centrifuge mini star silverline | | VWR |
| ThermoMixer C | | Eppendorf GmbH |
| Transwell Permeable supports | | Costar |
| Ultracentrifuge Optima L90K | | Beckman Coulter |
| Vortexer Reax top | | Heidolph instruments |
| Vortexer VWRR Galaxy Mini Star | | VWR International bvba |
| Waterbath GFL 1086 | | GFL Technology |
| **General consumables** | | |
| 8-Well PCR tube strips plus 8 domed caps | strips: 72.985.002, caps: 65.989.002 | Sarstedt |
| 96-Well Plate Advanced TC (flat bottom) | 655983 | Greiner Bio-one |
| Biosphere® Filter Tip 10 µl | 70.1130.210 | Sarstedt |
| Biosphere® Filter Tip 1000 µl | 70.762.211 | Sarstedt |
| Biosphere® Filter Tip 100 µl | 70.760.212 | Sarstedt |
| Cell culture multiwell plate, 6 well, PS, clear, sterile | 657160 | Greiner Bio-one |
| Cell culture plates CELLSTAR®, sterile, white-96-well plates | KL43.1 | Roth |
| CELLSTAR®, TC, lid with condensation rings, sterile | 655180 | Greiner Bio-one |
| CryoPure Tube 1.6 ml yellow | 72.380.004 | Sarstedt |
| Disposable needles Sterican® long bevel facet, 0.30x12 mm | 4656300 | B.Braun |
| Disposable Syringe, Luer, 1 ml | CH030001L | Charina |
| FACS tubes | 352052 | BD |
| Falcon® 10 ml Serological Pipet | 357551 | Corning |
| Falcon® 12-well Clear Flat Bottom TC-treated Multiwell Cell Culture Plate | 353043 | Corning |

| | | |
|---|---|---|
| Falcon® 12-well Clear Flat Bottom TC-treated Multiwell Cell Culture Plate | 353043 | Corning |
| Falcon® 15 ml High Clarity PP Centrifuge Tube | 352096 | Corning |
| Falcon® 2 ml Serological Pipet | 357507 | Corning |
| Falcon® 24-well Clear Flat Bottom TC-treated Multiwell Cell Culture Plate | 353047 | Corning |
| Falcon® 24-well Clear Flat Bottom TC-treated Multiwell Cell Culture Plate | 353047 | Corning |
| Falcon® 25 ml Serological Pipet | 357525 | Corning |
| Falcon® 35 mm TC-treated Easy-Grip Style Cell Culture Dish | 353001 | Corning |
| Falcon® 40 µm Cell Strainer | 352340 | Corning |
| Falcon® 48-well Clear Flat Bottom TC-treated Multiwell Cell Culture Plate | 353078 | Corning |
| Falcon® 5 ml Round Bottom Polystyrene Test Tube | 352052 | Corning |
| Falcon® 5 ml Round Bottom Polystyrene Test Tube, with Cell Strainer Snap Cap | 352235 | Corning |
| Falcon® 5 ml Serological Pipet | 357543 | Corning |
| Falcon® 50 ml High Clarity PP Centrifuge Tube | 352070 | Corning |
| FrameStar Fast Plate 96-well semi skirted | 4ti-1200 | 4titude |
| Injekt Solo-2-piece single-use syringe, 10 ml, Luer Lock | 201235 | B.Braun |
| MACS LS Columns | 130-042-401 | Miltenyi Biotec |
| MACS MS Columns | 130-042-201 | Miltenyi Biotec |
| Pasteurpipette glass, 145mm | 500635 | Brand |
| Pasteurpipette glass, 230mm | 500636 | Brand |
| Reagent Reservoirs, 25 ml | EKT8.1 | Carl Roth |
| Rotilab®-syringe filters, CA, sterile, 0.45µm | KC71.1 | Carl Roth |
| Sodium butyrate | B5887-1g | Sigma Aldrich |
| Spitzen Filter Surphob 100 µl (10x96) | VT0230 | Biozym |
| Spitzen Filter Surphob 10 µl lang (10x96) | VT0200 | Biozym |
| Spitzen Filter Surphob 1,250 µl (10x96) | VT0270 | Biozym |
| Stericup | CT92.1 | Carl Roth |
| Sterile, Stainless Steel, Premium Disposable Scalpel, 11PA | 03025 | Razormed |
| Transwell Permeable supports | 10107341 | Costar |
| Vasco® Nitril blue glove S | 9205518 | B.Braun |
| **Chemicals and reagents (general)** | | |
| 0.05% Trypsin-EDTA (1x) | 25300-096 | Gibco |
| 2x HEPES | S0874 | Takara |
| Albumin from bovine serum Fraktion V | 8076.3 | Carl Roth |
| Anti-human CD28 antibody | 302934 | Biolegend |
| Anti-PE microbeads | 130-105-639 | Miltenyi Invitrogen |
| Brefeldin A | #347688 | BD Biosciences |
| CD3 antibody anti-human, pure-functional grade clone OKT3 | 130-093-387 | Miltenyi |
| CD8 microbeads, human - lyophilized | 130-097-057 | Miltenyi |
| CS&T Research Beads | 655050 | Becton Dickinson |
| Cut Smart Buffer | NEB#B6004 | New England Biolabs |
| D-Luciferin | 122799 | Perkin Elmer Inc. |
| Dimethyl sulfoxide (DMSO) | A994.1 | Carl Roth |
| DISPASE II | D4693-1G | Sigma Aldrich |
| Dnase I | A3778.0100 | AppliChem |
| Dynabeads Human T-Activator CD3/CD28 | 11131D | ThermoFisher |
| Dynabeads Mouse T-Activator CD3/CD28 | 114530 | ThermoFisher |
| Ethanol Rotipuran 99.8% p.a. | 2065.2 | Carl Roth |
| Fetal Bovine Serum (FBS) SUPERIOR | S0615-500ML | Merck |
| Ficoll Paque Plus | 17144002 | Cytiva |
| Flow-Set Pro Fluorospheres | A62492 | Beckman Coulter |

| | | |
|---|---|---|
| FlowCheck Beads | A63493 | Beckman Coulter |
| FlowClean Cleaning Agent | A64669 | Beckman Coulter |
| Glycine Buffer ≥ 99% p.a. | 3908.3 | Carl Roth |
| $H_2O_2$ (2N) | 339741 | Sigma Aldrich |
| Human IL-15, premium grade | 130-095-765 | Miltenyi |
| Human IL-7, premium grade | 130-095-362 | Miltenyi |
| Human Fc Receptor Blocking Solution | 422301 | Biolegend |
| Hydrocortisone | H0396-100MG | Sigma Aldrich |
| Ionomycin | 10364-1MG | Sigma Aldrich |
| IsoFlow Sheath Fluid | 8546859 | Beckman Coulter |
| Isoflurane (CPS) | 1214 | Cp-pharma |
| KAPA HiFi HotStart Ready Mix | 7958927001 | Roche |
| L-Glutamine | 25020-081 | Life Technologies |
| LE Agarose | 240004 | Biozym |
| Live/Dead Fixable Near IR (780) Viability Kit, for 633 nm excitation | L34975 | ThermoFisher Scientific |
| Matrigel | 356237 | Corning |
| Monensin | M5273-500MG | Sigma Aldrich |
| Nuclease-free water | 1097705 | Invitrogen |
| Papain from papaya latex buffered aqueous solution | P3128-100MG | Sigma Aldrich |
| Penicillin-Streptomycin (10,000 U/mL) | 18140122 | Gibco |
| PhosSTOP | 4906837001 | Sigma Aldrich |
| PMA | P8139-1MG | Sigma Aldrich |
| Potassium chloride (KCl) ≥ 99.5% p.a. | 6781.1 | Carl Roth |
| Potassium dihydrogen phosphate ($KH_2PO_4$) ≥ 99% p.a. | 3904.1 | Carl Roth |
| Potassium hydrogen carbonate ($KHCO_3$) ≥ 99% | X887.1 | Carl Roth |
| Powdered milk, blotting grade | T145.3 | Carl Roth |
| Precision Plus Protein Standards | 161-0374 | Bio-Rad |
| Q5 Polymerase | M0491L | New England Biolabs |
| Retronectin | T1008 | Takara |
| SDS Pellets ≥ 99.9% | CN30.3 | Carl Roth |
| TEMED ≥ 98.5% | 2367.1 | Carl Roth |
| Tris Bufferan® ≥ 99.9% p.a. | 4855.5 | Carl Roth |
| Trypan Blue Solution 0.4% | 15250061 | Gibco |
| Tween®20 | 9127.2 | Carl Roth |
| Vibrant™ Dio Cell-Labeling Solution | V22886 | ThermoFisher |
| β-Mercaptoethanol 99% p.a. | 4227 | Carl Roth |
| **Cell culture media** | | |
| RPMI Medium 1640 (1x) | 21875-034 | Life Technologies |
| DMEM (1x) Dulbecco's Modified Eagle Medium | 41966052 | Life Technologies |
| Endothelial Cell Growth Medium 2 | C-22111/39211 | PromoCell |
| Opti-MEM | 31985062 | Life Technologies |
| MCDB 131 Medium | 10372019 | ThermoFisher |
| Freezing medium | 10% DMSO, 90% fetal calf serum (FCS) | |
| Tumor cell and HEK 293T cell medium with RPMI | RPMI Medium 1640 (1x), 10% FCS | |
| Tumor cell medium with DMEM | DMEM (1x), 10% FCS | |
| T cell (human) medium | RPMI Medium 1640 (1x), 10% FCS, 2 mM L-Glutamin | |

**HDRT-generation**

| | | |
|---|---|---|
| Q5 Hot Start High-Fidelity 2X Master Mix | M0494S | NEB |
| NEBuilder HiFi DNA Assembly Master Mix | E2621L | NEB |
| XL10-Gold Ultracompetent Cells | 200315 | Agilent |
| Kapa HiFi Hot Start ready mix | KK2602/7958935001 | Roche |
| AMPure XP, 60 mL | A63881 | Beckman Coulter |

| | | |
|---|---|---|
| DynaMag2 | 12321D | Thermo Fisher Scientific |
| DpnI | R0176S | New England Biolabs |
| Effectene Transfection Reagent | 301425 | Qiagen |
| **CRISPR knock-in** | | |
| Alt-R® S.p. Cas9 Nuclease V3, 500 µg | 1081059 | IDT |
| Poly-L-glutaminsäure Natriumsalz | P4761-100MG | Sigma-Aldrich |
| SF Cell Line 4D-NucleofectorTM X Kit L | V4XC-2024 | Lonza |
| P3 Primary Cell 4D-NucleofectorTM X Kit L | V4XP-3024 | Lonza |
| **dPCR** | | |
| Pipet-Lite Multi Pipette L8-200XLS+ | 17013805 | Mettler-Toledo |
| Rainin Pipette Tips TR LTS 200 µL F 960A/10 | 17014963 | Mettler-Toledo |
| Eppendorf twin.tec® PCR Plate 96, semi-skirted, clean | 30128575 | Eppendorf |
| DNA LoBind® Tubes | 30108051 | Eppendorf |
| QIAamp DNA Mini Kit (50) | 51304 | Qiagen |
| QIAcuity Probe PCR Kit (5 ml) | 250102 | Qiagen |
| QIAcuity Nanoplate 26k 24-well (10) | 250001 | Qiagen |
| QIAcuity Nanoplate 26k 8-well (10) | 250031 | Qiagen |
| QIAcuity Nanoplate 8.5k 24-well (10) | 250011 | Qiagen |
| QIAcuity Nanoplate 8.5k 96-well (10) | 250021 | Qiagen |
| Nanoplate Seals (11) | 250099 | Qiagen |
| Nanoplate Tray | 250098 | Qiagen |
| XbaI | R0145S | New England Biolabs |
| **Incucyte** | | |
| Incucyte® Clearview 96-well Reservoir Plate | 4600 | Sartorius |
| Incucyte® Clearview 96-well Plate for Chemotaxis | 4582 | Sartorius |
| Recombinant Protein G | 101200 | Invitrogen (Thermo) |
| ICAM-1 Protein, Human, Recombinant (ECD, His & hFc Tag) | 10346-H03H | SinoBiological |
| Incucyte® Cytotox Dye for Counting Dead Cells: Green | 4633 | Sartorius |
| Incucyte® Nuclight Rapid Red Dye for Live-Cell Nuclear Labeling | 4717 | Sartorius |
| **CAR T cells and Cytokine effects** | | |
| $CD8^+$ T cell isolation kit, human | 130-096-495 | Miltenyi |
| Pan T cell isolation kit, human | 130-096-536 | Miltenyi |
| Human CXCL10/IP-10 DuoSet ELISA, 15 Plate | DY266 | R&D |
| Recombinant human CXCL10 | 300-12-35 | Peprotec |
| Human CXCL11/I-TAC DuoSet ELISA | DY672 | R&D |
| Recombinant Human I-TAC (CXCL11) | 300-46 | Peprotec |
| Human IFN-gamma Recombinant Protein, PeproTech® | 300-02-100UG | Thermo fisher |
| Human IFN-gamma ELISA Kit | DIF50C | R&D |
| Human IL-2 ELISA | 555190 | Becton Dickinson |
| pCMV-Rev2 (p13.33) | | Takara |
| Viral Packaging PCHGP-2 (p14.36) | | Takara |
| pCMV-G (p15) | | Takara |
| UltraPure™ BSA (50 mg/mL) | AM2616 | Thermo Fisher Scientific |
| ICAM-1 | #10346-H03H | Sino Biological |
| Protein G | #101200 | Thermo Fisher |
| 96-well Essen ClearView Reservoir, | 4600, 4601 | Sartorius |
| Effectene | 301425 | Qiagen |
| Optimem I | 31985070 | Gibco |

Supplementary Table 6. Antibodies used for flow cytometry and IF Staining

| Antibodies | Clone | Dilution | Catalogue no. | Provider |
|---|---|---|---|---|
| Invitrogen, CD34 Monoclonal Antibody (QBEND/10), PE | 537860 | 1:10 | #MA1-10205 | ThermoFisher Scientific |

| | | | | |
|---|---|---|---|---|
| CXCR3 (CD138) Alexa Fluro 700 human Ab, Clone G025H7, Mouse IgG1 | | | 353741 | Biolegend |
| CXCR3, PE | | 1:25 | 353706 | Biolegend |
| CXCR3, AX700 | | 1:25 | 353741 | Biolegend |
| B7H3, APC | | | | Biolegend |
| B7H3, PE | | | 331606 | Biolegend |
| L1CAM (CD171), APC | REA163 | 1:100 | 130-124-024 | Miltenyi |
| L1CAM (CD171), PE | REA163 | 1:100 | | Miltenyi |
| CD4, FITC | | 1:400 | 357406 | Biolegend |
| CD4, BV421 | RPA-T4 | 1:50 | | Biolegend |
| CD8, PerCP | | 1:400 | | Biolegend |
| CD8, BV605 | SK1 | 1:400 | | |
| CD3, Ax700 | HIT3a | 1:400 | 300324 | Biolegend |
| CD3, FITC | HIT3a | 1:400 | | Biolegend |
| CD3, PerCP-Cy5.5 | HIT3a | 1:400 | | Biolegend |
| CD137, BV421 | 4B4-1 | 1:50 | 309819 | Biolegend |
| CD137, BV605 | 4B4-1 | 1:50 | | Biolegend |
| CD25, PE-Cy7 | BC96 | 1:100 | 302611 | Biolegend |
| CD25, Ax700 | BC96 | 1:25 | | Biolegend |
| EGFRt (Cetuximab), PE | | 1:100 | 537860 | Invitrogen |

#### Supplementary Table 7. Software used

| Software | Use | Provider |
|---|---|---|
| FlowJo_v.10.6.2 | Analysis of flow cytometry data | FlowJo |
| Gen5_v2.04 | ELISA assay measurement | BioTek |
| GraphPad PRISM_v8 to v10 | Data analysis and graphical presentation | GraphPad |
| Incucyte 2021B | Analysis of live cell imaging | Essen Bioscience |
| Primer3Plus | Primer design and adjustment | Free software, Copyright (c) 2006, 2007 by Andreas Untergasser and Harm Nijveen |
| ProSort_v1.6 | FACS Sorter | Bio-Rad |
| SAP | Patient data collection | SAP SE |
| SnapGene | Development of cloning strategies, new Plasmids | Dotmatics |
| StepOne Software_v2.3 | qRT-PCR set-up and analysis | Applied Biosystems |
| VisionCapt_v16.16d | Western Blot visualization | Fusion |
| Python version 3.11.4. | CancerPAM programming | |
| Visual Studio Code (Version: 1.80.1) | source-code editor | |
| Snakemake | Pipeline workflow management system | |
